# Minimal phenotyping yields GWAS hits of reduced specificity for major depression

**DOI:** 10.1101/440735

**Authors:** Na Cai, Joana A. Revez, Mark J Adams, Till F. M. Andlauer, Gerome Breen, Edna M. Byrne, Toni-Kim Clarke, Andreas J. Forstner, Hans J. Grabe, Steven P. Hamilton, Douglas F. Levinson, Cathryn M. Lewis, Glyn Lewis, Nicholas G. Martin, Yuri Milaneschi, Ole Mors, Bertram Müller-Myhsok, Brenda W. J. H. Pennix, Roy H. Perlis, Giorgio Pistis, James B. Potash, Martin Preisig, Jianxin Shi, Jordan W. Smoller, Fabien Streit, Henning Tiemeier, Rudolf Uher, Sandra Van der Auwera, Alexander Viktorin, Myrna M. Weissman, MDD Working Group of the Psychiatric Genomics Consortium, Kenneth S. Kendler, Jonathan Flint

## Abstract

Minimal phenotyping refers to the reliance on the use of a small number of self-report items for disease case identification. This strategy has been applied to genome-wide association studies (GWAS) of major depressive disorder (MDD). Here we report that the genotype derived heritability (h^2^_SNP_) of depression defined by minimal phenotyping (14%, SE = 0.8%) is lower than strictly defined MDD (26%, SE = 2.2%). This cannot be explained by differences in prevalence between definitions or including cases of lower liability to MDD in minimal phenotyping definitions of depression, but can be explained by misdiagnosis of those without depression or with related conditions as cases of depression. Depression defined by minimal phenotyping is as genetically correlated with strictly defined MDD (rG = 0.81, SE = 0.03) as it is with the personality trait neuroticism (rG = 0.84, SE = 0.05), a trait not defined by the cardinal symptoms of depression. While they both show similar shared genetic liability with neuroticism, a greater proportion of the genome contributes to the minimal phenotyping definitions of depression (80.2%, SE = 0.6%) than to strictly defined MDD (65.8%, SE = 0.6%). We find that GWAS loci identified in minimal phenotyping definitions of depression are not specific to MDD: they also predispose to other psychiatric conditions. Finally, while highly predictive polygenic risk scores can be generated from minimal phenotyping definitions of MDD, the predictive power can be explained entirely by the sample size used to generate the polygenic risk score, rather than specificity for MDD. Our results reveal that genetic analysis of minimal phenotyping definitions of depression identifies non-specific genetic factors shared between MDD and other psychiatric conditions. Reliance on results from minimal phenotyping for MDD may thus bias views of the genetic architecture of MDD and may impede our ability to identify pathways specific to MDD.

## Introduction

There is now little doubt that a key requisite for the robust identification of genetic risk loci underlying psychiatric disease is the use of an appropriately large sample. However, while the costs of genotyping and sequencing continue to fall, the cost of phenotyping remains high^1^, limiting sample collection. One solution for reducing the burden of case identification is to use clinical information from hospital registers^2^, or reliance on subjects’ self-reported symptoms, help-seeking, diagnoses or medication, which are cheap and fast to collect. We refer to the latter strategy as “minimal phenotyping”, as it minimizes phenotyping costs and reduces data to a single or few self-reported answers.

However, apart from the detection of more and more GWAS loci^3–5^ (Supplemental Table S1), the consequences of sacrificing symptomatic information for genetic analyses has rarely been investigated. The consequences may be particularly important for major depressive disorder (MDD) because of its phenotypic and likely etiological heterogeneity^6^, high degree of comorbidity with other psychiatric diseases^7^, and substantial discrepancies between self-assessment using symptom scales and diagnoses made with full diagnostic criteria^8^. While a majority of the population self-identify as having one or two depressive symptoms at any one time, only between 9 and 20% of the population have sufficient symptoms to meet criteria for lifetime occurrence of MDD^8–10^. A number of factors affects self-reported diagnosis or prescribed treatment: among those with MDD only half consult a doctor for help^11–13^, there are high rates of false positives when diagnoses are made without applying diagnostic criteria^12^, and antidepressants are prescribed for a wide range of conditions other tham MDD^13–15^. As such, a cohort of MDD cases obtained through the use of self-report of either illness or prescribed treatment may yield a sample that is not representative of the clinical disorder, but enriched in those with non-specific sub-clinical depressive symptoms and depression secondary to a comorbid disease.

By comparing the genetic architecture of minimal phenotyping definitions of depression with those using full diagnostic criteria for MDD in UKBiobank^16^, a community-based survey of half a million men and women, we assess the implications of a minimal phenotyping strategy for GWAS in MDD. We find that MDD defined by minimal phenotyping has a large non-specific component, and if GWAS loci from these definitions are chosen for follow-up molecular characterization, they may not be informative about biology specific to MDD.

## Results

### Definitions of depression in UKBiobank

The diverse assessments of depression in the UK Biobank provide an opportunity to determine the impact that differences in diagnostic criteria have on findings of depression’s genetic architecture. We identified five ways that MDD can be defined in UKBiobank. First, using self-reports of seeking medical attention for depression or related conditions, we identified “Help-seeking” definitions of MDD (referred to as “broad depression” in a previous GWAS^3^). Second, participants were diagnosed with “Symptom-based” MDD if, in addition to meeting the help seeking criteria described above, they reported ever experiencing one or more of the two cardinal features of depression (low mood or anhedonia) for at least two weeks (this is the “probable MDD” diagnosis available in UKBiobank^17^). Third, we define a “Self-Report” form of depression based on participants’ self-reports of all past and current medical conditions to trained nurses. Fourth, information from electronic health records can be used to assign an ICD10 primary and secondary illness codes from the label as “EMR”. Finally, a “CIDI-based” diagnosis of lifetime MDD was derived from subjects who answered an online “Mental Health Follow-up” questionnaire (MHQ)^18^ based on the Composite International Diagnostic Interview Short Form (CIDI-SF)^19^, which included DSM-5 criteria for MDD (Supplemental Methods, Supplemental Figure S1, Supplemental Table S2). None of the definitions uses trained interviewers applying structured clinical interviews, and only the last applies operationalized criteria including symptoms, length of episode (more than two weeks) and impaired social, occupational or educational function. From hereon we refer to definitions one to three as ‘minimal’, the fourth as “EMR-based”, and the fifth as ‘strictly’ defined MDD (Supplemental Methods).

We also included a category of participants who met the Help-seeking based definition (part of “broad depression” in Howard et al 2018^3^) but failed to meet the symptom based definition (as they had neither of the two cardinal symptoms of depression: depressed mood or a loss of interest or pleasure in daily activities for more than two weeks). This group we refer to as “Non-MDD” (described in detail in Supplemental Methods, Supplemental Table S3), and is enriched for those who sought help for depression but do not have MDD.

All definitions are based on recall of episodes or symptoms of depression by participants in the UKBiobank. As priming of recall by current mood affects the reliability of such reports^20–22^, we emphasize that each definition is noisy, and can be interpreted as being enriched for individuals truly fulfilling its criteria. Figure 1 outlines the different diagnostic categories and the numbers of samples that each contains.

**Figure1:**
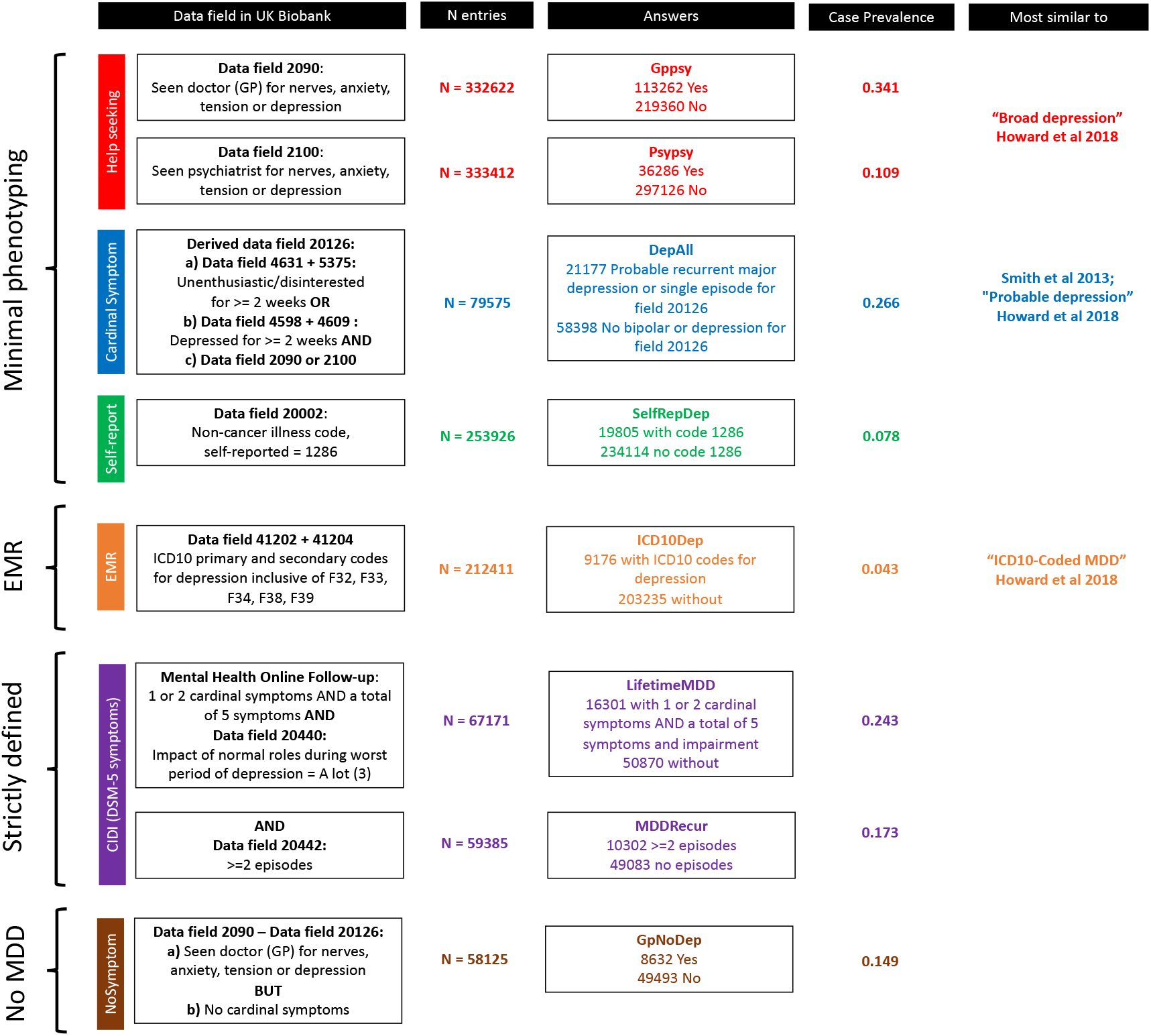
Definitions of depression in UKBiobank. This figure shows the different definitions of MDD in UKBiobank and the colour codings we use consistently in this paper. For the minimal phenotyping definitions of depression we present in this paper: Red for help-seeking based definitions derived from Touchscreen Questionnaire; blue for symptom based definitions derived from Touchscreen Questionnaire; green for self-report based definition derived from Verbal Interview. For the EMR definition of depression: orange for definitions based on ICD10 codes. For strictly defined MDD: purple for CIDI-based definitions derived from Online Mental Health Followup. For the no-MDD definition: brown for GPNoDep, containing those cases in help-seeking definitions that do not have cardinal symptoms for MDD. The data fields in UKBiobank relevant for defining each phenotype are shown in “Data field in UK Biobank”; number of individuals with non-missing entries for each definition are shown in “N entries”; the qualifying answers for cases and controls respectively are shown in “Answers”; the case prevalences in each definition are shown in “Case Prevalence”; the study and definitions of depression most similar to our definitions are shown in “Most similar to”. The similarities and differences between help-seeking, EMR, and symptom-based definitions with definitions of depression previously reported can be found in Supplemental Methods.

There are several features to these definitions of MDD we recognize as important for the comparison of their genetic architecture. First, for each diagnostic category we used controls who were asked the relevant questions but failed to meet criteria. As such, cases from one category can be controls in another, resulting in substantial overlap in both cases and controls between categories (Supplemental Figure S2), which impacts assessment of genetic architecture and genetic correlation between them^23^. Second, not all participants in UKBiobank were asked questions from all categories. For example, questions for the Symptom-based definition DepAll were asked in only 10 out of 22 assessment centers in UKBiobank (Supplemental Table S4), and the MHQ was only answered in full by 31% of the original UK Biobank participants, resulting in differences in population structure between definitions that need to be considered in analysis (Supplemental Methods, Supplemental Figure S3).

Third, different definitions of depression have different prevalence in the UKBiobank cohort (from 0.078 to 0.341, Supplemental Table S5). Though unlikely biased due to population structure (Supplemental Tables S6-7), the wide range of prevalence may be due to self-ascertainment biases. In general, the UKBiobank is known for higher participation rates from women who are older, more well-off, and better educated^24^. One example of self-ascertainment of particular interest to our analysis is the voluntary participation in the MHQ, which has been shown to have a genetic component that can be genetically correlated with that of mental health conditions^25^. All such biases can have confounding effects on genetic studies of depression phenotypes derived from it. We verified that, though ascertainment biases exist, they cannot account for the results we present in this paper (Supplemental Methods, Supplemental Figure 4, Supplemental Table S8-9).

Finally, many previous GWAS on depression (and any other disease) apply other filters in the selection of cases and controls, in addition to case criteria. One such example is the use of “clean controls” which requires individuals who are designated as “controls” do not endorse any of the case criteria. We do not use “clean controls” in this study, as that violates key assumptions in our analyses of genetic architecture (Supplemental Methods), although we show this strategy can increase power in GWAS as compared to using all controls (Supplemental Tables S10-11, Supplemental Figure S5).

### Minimal phenotyping definitions of depression are epidemiologically different from strictly defined MDD

We began our examination of the different definitions of MDD by looking at how known risk factors for depression impacted on each. We assessed whether risk factors were similar between definitions of depression^26^. Figure 2a-g shows the mean effect (odds ratio, OR) with confidence intervals of each of the following risk factors: sex^27,28^, age^29^, educational attainment^30–32^, socio-economic status^33^, neuroticism^28,34^, experience of stressful life events in the two years leading up to the baseline assessment, and cumulative traumatic life events preceding assessment^35,36^ (Supplemental Methods, Supplemental Table S12). Estimates of the risk factor effect sizes differed substantially, and often highly significantly, as shown by the confidence intervals in Figure 2. These may reflect differences in methods of ascertainment, or underlying pathology and genetic architecture, between definitions of depression. We asked if differences in risk factors can be used to classify definitions of depression. We applied a clustering algorithm and found that all minimal phenotyping definitions of depression cluster separately from strictly defined MDD (Figure 2h).

**Figure2:**
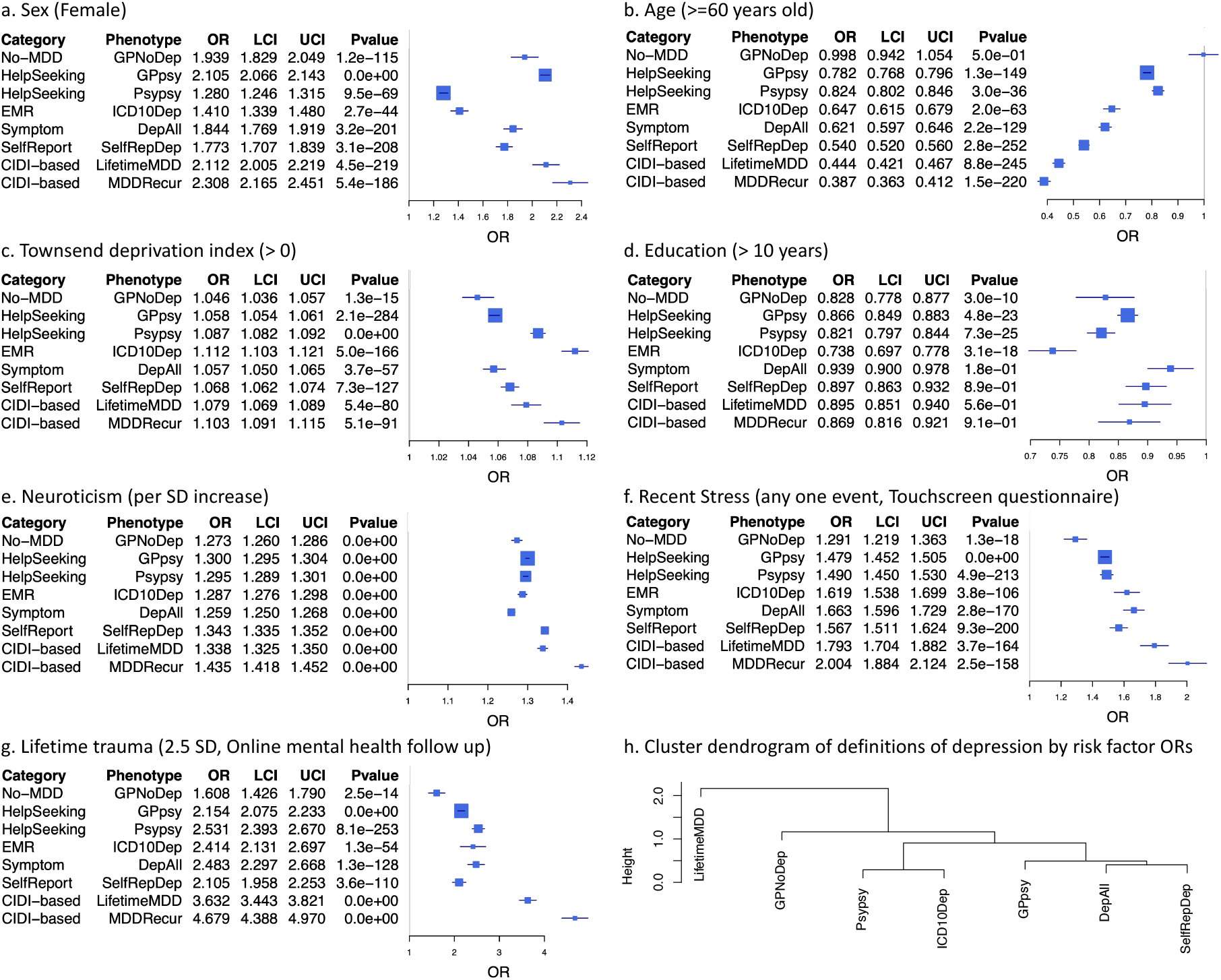
Relationship between definitions of depression and environmental risk factors. a-g) These figures show forest plots of odds ratios (OR) and −log10 P values (LogP) between known environmental risk factors and different types (Category) of definitions of depression in UKBiobank (Definition) from logistic regression, using UKBiobank assessment centre, age, sex and years of education as covariates to control for potential geographical and demographic differences between environmental risk factors, except when they are being tested. Lifetime trauma measure was derived from Online Mental Health Followup (Supplemental Methods, Supplemental Table S7); Townsend deprivation index, years of education, sex, age, recent stress and neuroticism were derived from Touchscreen Questionnaire (Supplemental Methods). f) This figure shows a hierarchical clustering of definitions of depression in UKBiobank using ORs with environmental risk factors performed using the hclust function in R, “Height” refers to the Euclidean distance between MDD definitions at the ORs of all six risk factors. MDDRecur is not included in this clustering analysis as it is a subset of the LifetimeMDD definition.

### Minimal definitions of depression are not just milder or noisier version of strictly defined MDD

We next addressed the question of the genetic relationship between the different MDD definitions. We found that depression defined by minimal phenotyping strategies have lower SNP-based heritabilities (h^2^_SNP_) than more strictly defined definitions (Figure 3a). Self-report (SelfRepDep h^2^_SNP_ = 11%, se = 0.85%) and help-seeking based definitions (Psypsy h^2^_SNP_ = 13%, se = 1.18%; GPpsy h^2^_SNP_ = 14%, se = 0.81%) have heritabilities of 15% or less. By contrast, strictly defined MDD (LifetimeMDD) has a much higher h^2^_SNP_ of 26% (se = 2.15%); imposing the further criterion of recurrence brings the h^2^_SNP_ up to 32% (se = 2.56%). Other definitions have intermediate h^2^_SNP_.

**Figure3:**
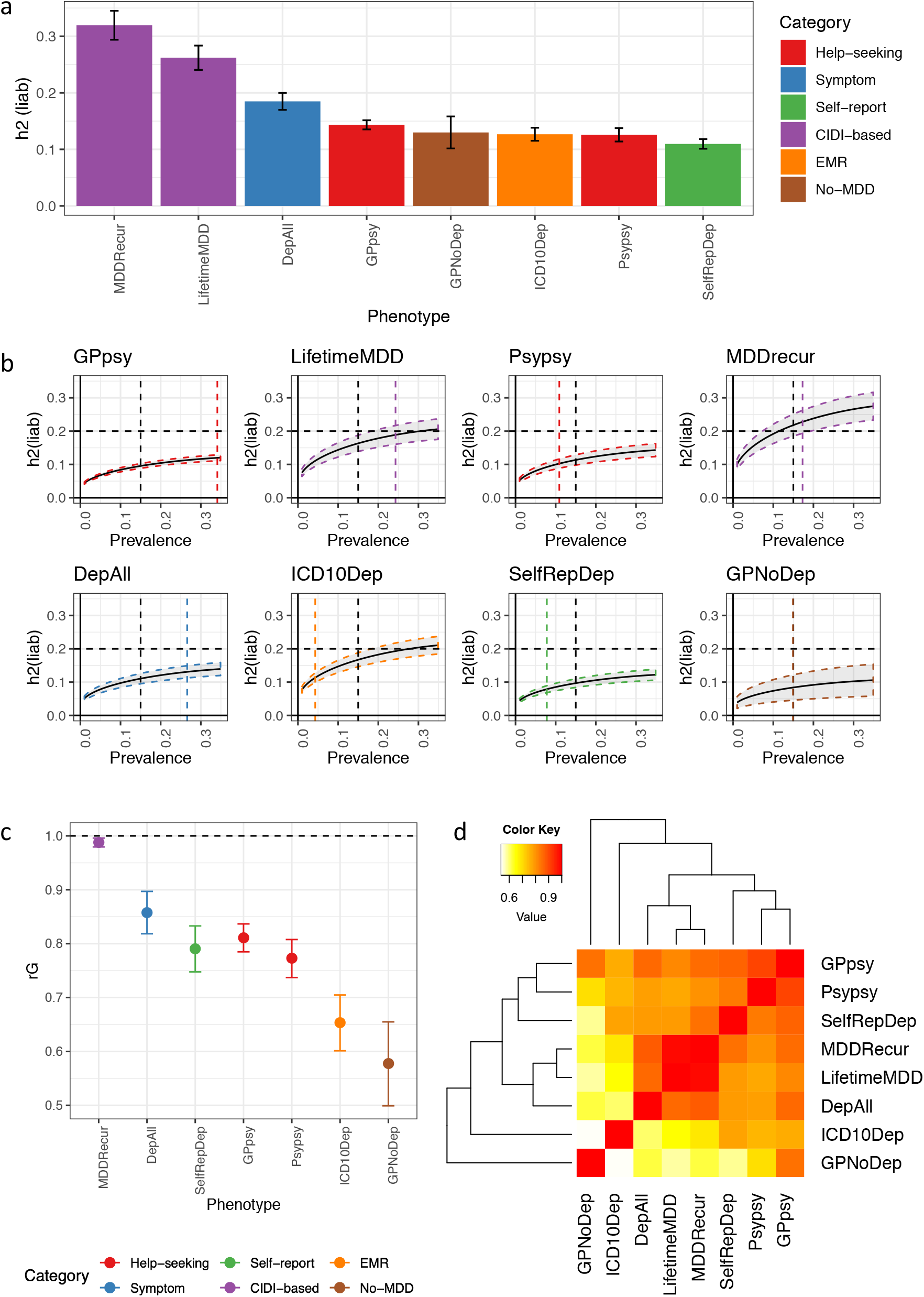
SNP-heritability and genetic correlation estimates among definitions of MDD in UKBiobank. a) This figure shows the h^2^_SNP_ estimates from PCGCs^19^ on each of the definitions of MDD in UKBiobank (Methods). h^2^_SNP_ “h2(liab)” as shown on the figure has been converted to liability scale^44,73^ using the observed prevalence of each definition of depression in UKBiobank as both population and sample prevalences (Supplemental Table S4). b) This figure shows the h^2^_SNP_ estimates of definitions of MDD in UKBiobank from LDSC using logistic regression summary statistics on all SNPs > 5% MAF (Methods), transformed to the liability scale assuming a range of population case prevalence, from 0 to 0.5. We do not show results for case prevalence from 0.5 to 1, as they will be mirroring those from 0 to 0.5. In the figure we indicate with a black vertical dotted line the population prevalence of 0.15, used in PGC1-MDD, and a coloured vertical dotted line for the population prevalence of each definition of depression in UKBiobank. We also indicate with a black horizontal dotted line the arbitrary liability scale h^2^_SNP_ of 0.2, previously estimated for MDD in PGC1-MDD. Using this, we show that at no prevalence would minimal phenotyping defined depression like GPpsy (Help-seeking definition) reach this estimate. c) This figure shows the genetic correlation “rG” between CIDI-based LifetimeMDD and all other definitions of MDD in UKBiobank, estimated using PCGCs. d) This figure shows pairwise rG between all definitions of depression in UKBiobank, also detailed in Supplemental Table SX.

All h^2^_SNP_ estimates were estimated on the liability scale using PCGCs^23^, a method specifically suited for analyses of case-control data with ascertainment bias, where covariates (especially those strongly correlated with the disease, induced by ascertainment biases) are appropriately handled(Supplemental Methods). We ensured that the trend we observe holds regardless of the method used^23,37–39^ (Supplemental Methods, Supplemental Table S13), and was not affected by regions of high linkage-disequilibrium or complexity^40^ (Supplemental Methods, Supplemental Figure S3). For comparison, we provide h^2^_SNP_ estimates from previous studies of MDD^4,41,42^ (Supplemental Figure S6) and find that they fit squarely into the trend we observe: the less strict the criteria used to diagnose MDD, the lower the h^2^_SNP_.

We explored the impact of case prevalence on estimates of h^2^_SNP_, since self-ascertainment biases prevalence rates derived from UK Biobank data. We used case prevalence corrected for age, sex and regional participation in UKBiobank^43^ (Supplemental Methods, Supplemental Tables S5), as well as a constant case prevalence of 0.15, as used in previous studies on depression^5,42^. Neither alters our results (Supplemental Figure S3, Supplemental Tables S13). We further asked if self-ascertainment biases that could not be accounted for by age, sex and regional participation could explain the different h^2^_SNP_ estimates. We estimated h^2^_SNP_ for each definition of depression in UKBiobank using a range of prevalences from 0 to 0.35, and asked if any combination of prevalences would allow all definitions to have similar h^2^_SNP_ that is also consistent with previous estimates of 20%^42^. We found no realistic combination of prevalence across the definitions that would allow this (Figure 3b, See Supplemental Table S1 for all prevalence used in previous GWAS).

We examined the role of a number of additional factors for the lower h^2^_SNP_ of minimal phenotyping definitions of MDD. First, we established that minimal phenotyping definitions do not simply have a higher environmental contribution to MDD than the stricter definitions. When we assessed h^2^_SNP_ in MDD cases with high and low exposure to environmental risk factors^44^ we found that minimal phenotyping definitions of depression (GPpsy, SelfRepDep) show no significant difference between exposures, similar to or lower than strictly defined MDD (LifetimeMDD and MDDRecur) (Supplemental Methods, Supplemental Table S14).

Second, the minimal phenotyping definitions do not merely include milder cases of MDD as previously hypothesized^43^. Inclusion of milder cases is equivalent to lowering the threshold for disease liability in the population above which “cases” for MDD are defined. Under the liability threshold model^45^, this does not reduce the h^2^_SNP_, inconsistent with the lower h^2^_SNP_ in minimal phenotyping definitions of depression (Supplemental Methods, Supplemental Figure S8). Instead, we show, through simulations, that the lower h^2^_SNP_ of minimal phenotyping definitions of depression may be due to misdiagnosis of controls as cases of MDD, and misclassifications of those with other conditions as cases of MDD (Supplemental Figure S8, Supplemental Figure S9). The latter is more consistent with the high genetic correlations between minimal phenotyping definitions of MDD and GPNoDep, the no-MDD definition where cases do not have cardinal symptoms for MDD. However it should be noted that no-MDD definitions likely include some proportion of true cases of MDD.

### Genetic correlations between definitions of depression and other diseases

We found that the genetic correlation (rG) between minimal and strictly defined MDD includes a large proportion of non-specific liability to mental ill-health. The rG between GPpsy (minimal defined MDD) and LifetimeMDD (strictly defined MDD) is 0.81 (se = 0.03), significantly different than unity (Figure 3c-d, Supplemental Table S15, Supplemental Figure S6, Supplemental Methods). One interpretation of this finding is that the correlation represents shared genetic liability to MDD^4,5^. However, the majority of the genetic liability of LifetimeMDD due to GPpsy (approximately rG^2^ = 0.81^2^=66%), is shared with the No-MDD definition, GPNoDep, as the genetic liability of GPNoDep explains approximately 70% of the genetic liability of GPpsy (rG = 0.84, se = 0.05), and 34% of that of LifetimeMDD (rG = 0.58, se = 0.08).

We next examined rG between different definitions of MDD and comorbid diseases. We used cross-trait LDSC^46^ to estimate rG with neuroticism and smoking (Supplemental Figure S10, Supplemental Tables S16-17) in UKBiobank, as well as with all psychiatric conditions in the Psychiatric Genomics Consortium (PGC)^47^ including PGC1-MDD^42^ and depression defined in 23andMe^4^ (Supplemental Table S1). Figure 4a and Supplemental Table S18 shows few differences in rG estimates between other psychiatric disorders and the different definitions of MDD in UKBiobank, consistent with previous reports^48^.

However, similar rG estimates can result from different genetic architectures, indexed by the extent to which genetic liability is spread across the genome: it is spread more widely in a highly polygenic trait than a less polygenic trait. We estimated local rG_L_ and percentage contribution to total rG_T_ using rho-HESS^49^ (Methods, Figure 4b). 65.8% (SE = 0.6%), 37.1% (SE = 4.5%) and 42.7% (SE = 2.3%) of the genome explains 90% of the rG_T_ between strictly defined MDD (LifetimeMDD) and neuroticism, BIP and SCZ respectively. In comparison, 80.2% (SE = 0.6%), 47.3% (SE = 2.4%) and 46.8% (SE = 0.2%) of the genome is needed to explain the same percentage of rG_T_ between help-seeking based GPpsy and the same conditions (Figure 4c). In other words, minimal phenotyping definitions of depression share more genetic loci with other psychiatric conditions than strictly defined MDD does.

**Figure4:**
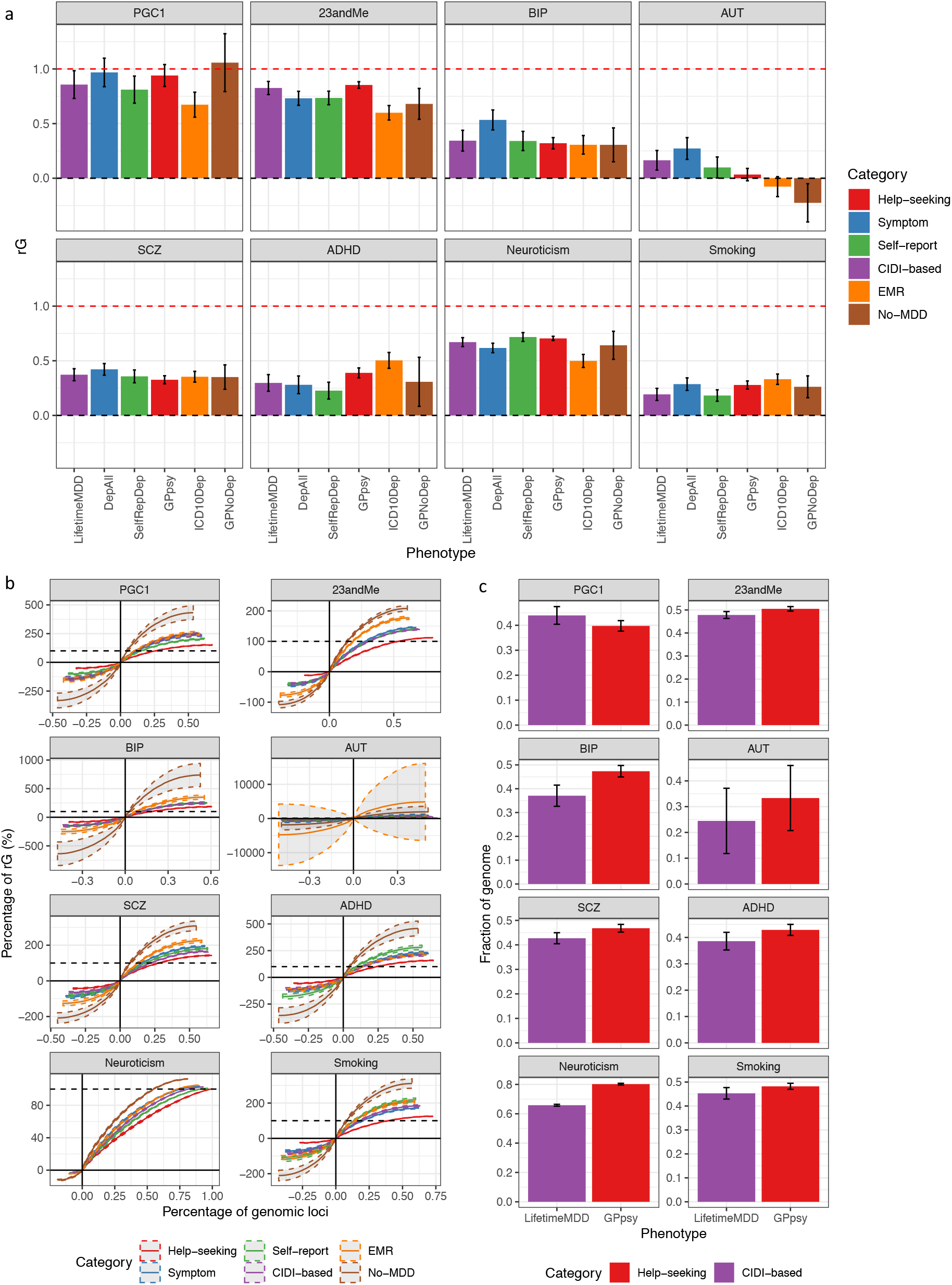
Genetic correlation between definitions of MDD and other psychiatric conditions. a) This figure shows the genetic correlation “rG” estimated by cross-trait LDSC^46^ on the liability scale between definitions of MDD in UKbiobank with other psychiatric conditions in both UKBiobank (smoking and neuroticism) and PGC^47^ (Supplemental Table S1), including schizophrenia^53^ (SCZ) and bipolar disorder^54^ (BIP) (Supplemental Table S1). b) This figure shows the cumulative fraction of regional genetic correlation “rG” (out of sum of regional genetic correlation across all loci) between definitions of MDD in UKBiobank with SCZ in 1703 indepedent loci in the genome^79^ estimated using rho-HESS^49^, plotted against percentage of independent loci. CIDI-based LifetimeMDD is shown in purple while help-seeking based GPpsy is shown in red. The steeper the curve, the smaller the number of loci explain the total genetic correlation. The dotted coloured curves around each solid line represent the standard errors of the estimate computed using a jackknife approach as described in Shi et al 2016^39^. The dotted black line represents 100% of the sum of genetic correlation between each definition of MDD in UKBiobank with SCZ. The cumulative sums of positive regional genetic correlations (right of y axis) go beyond 100% – this is mirrored by the negative regional genetic correlation (left of y axis) that go below 0%. c) We rank all 1703 loci by their magnitude of genetic correlation, and ask what fraction of loci sums up to 90% of total genetic correlation. This figure shows the percentage of loci summing up to 90% of total genetic correlation “rG” between either LifetimeMDD (in purple) or GPpsy (in red) with all psychiatric conditions tested, with standard errors estimated using the same jackknife approach. The higher the percentage, the higher the number of genetic loci contributing to 90% of total genetic correlation.

Previous work^4^ reported that depression defined through minimal phenotyping shows enrichment of h^2^_SNP_ in regions of the genome encoding genes specifically and highly expressed in central nervous system (CNS) tissues represented in GTEx^50^. We assessed this in the definitions of depression in UKBiobank using LDSC-SEG^51^. As shown in Figure 5, neither strictly defined MDD (LifetimeMDD) nor MDD defined based on structured clinical assessments in PGC1-MDD show significant CNS enrichments. This is inconsistent with CNS enrichment in a larger MDD cohort from PGC, PGC29-MDD^5^ (Methods, Supplemental Table S1, Supplemental Figure S11), and a previously reported meta-analysis between PGC29-MDD and cohorts collected in other strategies^5^, including a dominant contribution from the minimal phenotyping defined depression in 23andMe^4^. As the latter is a heterogeneous cohort collected with both structured interviews and electronic health records with varying degree of adherence to DSM-5 criteria^52^ (Supplemental Figure S12), this discrepancy reflects factors influencing results from enrichment analyses, including statistical power, diagnoses strategy, strictness of DSM-5 diagnostic criteria. We provide a full discussion in Supplemental Methods.

**Figure5:**
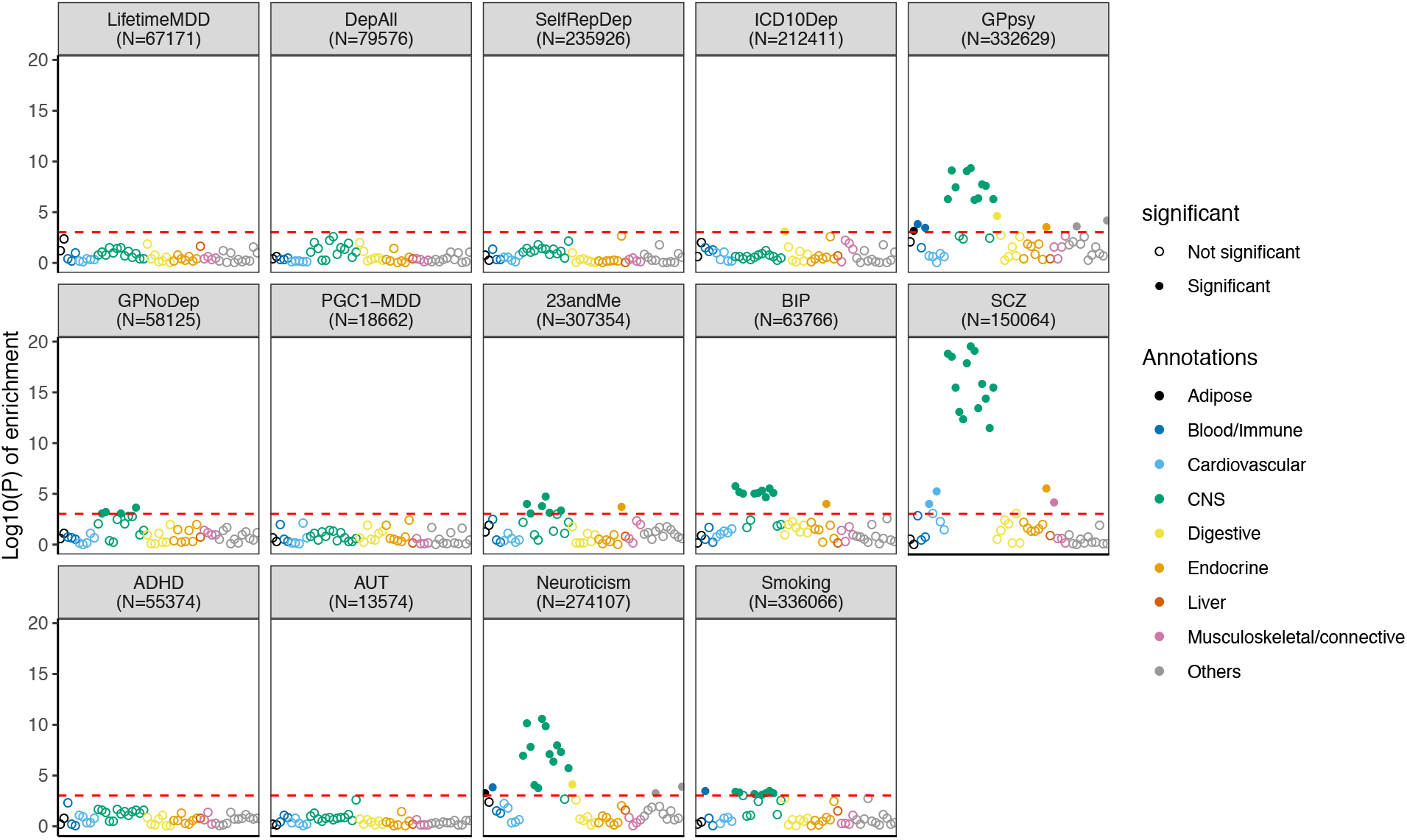
Tissue-specific gene expression enrichment in definitions of MDD. This figure shows the - log10(P) of enrichment in h^2^_SNP_ in genes specifically expressed in 44 GTEx tissues, estimated using partitioned h^2^_SNP_ in LDSC; help-seeking based definitions of MDD GPpsy, as well as its constituent no-MDD phenotype GPNoDep, show enrichment of h^2^_SNP_ in genes specifically expressed in CNS tissues, similar to an independent cohort of help-seeking based MDD (23andMe^4^) and other psychiatric conditions such as bipolar disorder (BIP)^54^, schizophrenia (SCZ)^53^, autism (AUT), personality dimension neuroticism, and behavioural trait smoking. We indicate the sample size (N) for each definition of depression and psychiatric condition.

Also shown in Figure 5, minimal phenotyping definition GPpsy showed a significant CNS enrichment (also see Supplemental Methods, Supplemental Figure S10). Notably, the non-MDD help-seeking definition GPNoDep also showed enrichment of h^2^_SNP_ in genes specifically expressed in CNS, as did neuroticism, smoking, and other disorders in the PGC^47^ such as schizophrenia^53^ (SCZ) and bipolar disorder^54^ (BIP). CNS enrichment therefore indexes genetic effects for many conditions rather than specifically MDD, and cannot alone be used to validate minimal phenotyping definitions of depression as being biologically relevant or equivalent to strictly-defined MDD.

### GWAS hits from minimal phenotyping are not specific to MDD

We next examined the specificity of action of individual genetic loci found in GWAS of each definition of MDD. We found that the help-seeking definitions gave the greatest number of genome-wide significant hits (27 from GPpsy and Psypsy, Supplemental Table S10) in GWAS, consistent with their larger sample sizes and statistical power for finding associations. Are these loci relevant and specific to MDD, or are they nonspecific and shared with other conditions such as SCZ, neuroticism and smoking?

Of the 27 loci from minimal phenotyping definitions, 10 showed significant effects (at P<0·05 after multiple testing correction for 27 loci) on strictly defined LifetimeMDD, despite the latter’s much smaller sample size. This is consistent with the hypothesis that using minimal phenotyping for GWAS of MDD can be useful for identifying loci relevant to MDD. However, all 10 loci also showed significant effects in neuroticism, smoking, SCZ, or the no-MDD help-seeking condition (GPNoDep, Supplemental Table S19). Furthermore, regardless of the absolute P value of association, the effects of the SNPs always go in the same direction for strictly-defined MDD and the non-MDD phenotypes, as shown in Figure 6. Hence, many of the loci relevant for MDD found using GWAS on minimal phenotyping definitions are not specific to MDD.

**Figure6:**
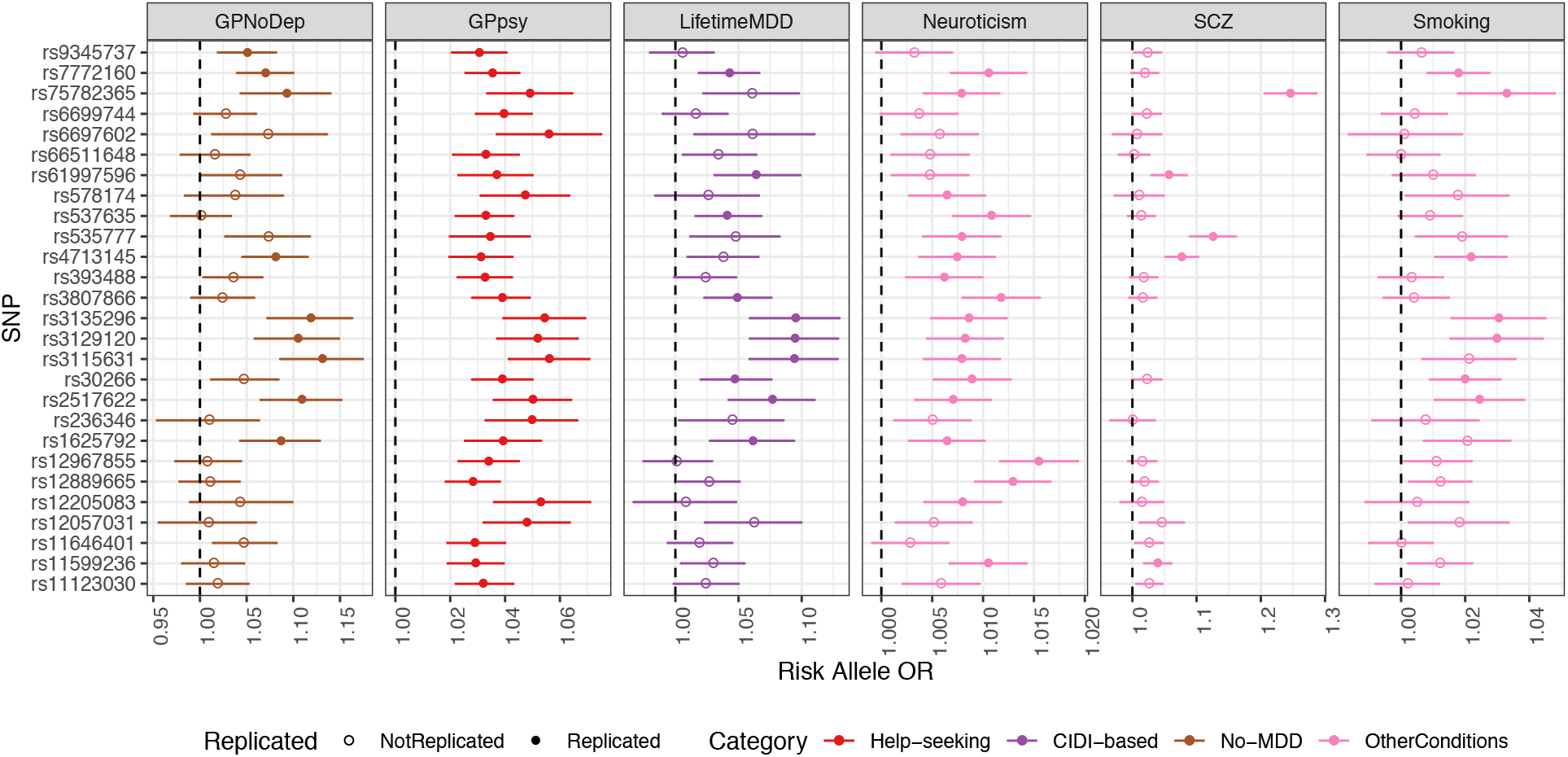
GWAS hits from minimal phenotyping definition of MDD in UKBiobank are not specific to MDD. This figure shows the odds ratios (ORs) for the risk alleles at 27 loci significantly associated with help-seeking based definitions of MDD in UKBiobank (GPpsy and Psypsy), in GWAS conducted on CIDI (LifetimeMDD, in purple), help-seeking (GPpsy in red) and no-MDD (GPNoDep, in brown) based definitions of MDD. For comparison we show the same in conditions other than MDD: neuroticism, smoking and SCZ (all in pink). SNPs missing in each panel are not tested in the respective GWAS. For clarity of display, scales on different panels vary to accommodate the different magnitudes of ORs of SNPs in different conditions. ORs at all 27 loci are highly consistent across phenotypes, being completely aligned in direction of effect, regardless of whether it is a definition or MDD or a risk factor or condition other than MDD. All results shown in Supplemental Table S14).

We find the same pattern of results when we use loci identified from a minimal phenotyping strategy in an independent study. The consumer genetics company 23andMe mapped loci that contribute to the risk of MDD defined by self-reports of receiving a doctor’s diagnosis and treatment of depression^4^. Of the 17 loci, ten replicated in GPpsy (at P<0·05, after multiple testing correction for 17 loci), only 3 replicated in LifetimeMDD. Effects at all of the loci go in the same direction in neuroticism, smoking or SCZ (Supplemental Figure S11, Supplemental Table S20) and are therefore not specific to MDD. These results are consistent with what we observe of minimal phenotyping definitions in UKBiobank, and show that they primarily enable the discovery of pathways associated with depression that are shared with other conditions. It is not currently possible to assess the specificity of GWAS loci from strictly defined MDD in the same way, given the sample size of strictly defined MDD remains relatively small, and GWAS hits relatively few.

### Out-of-sample prediction of MDD

Finally, we explored whether the definitions of depression in UKBiobank are genetically different by testing the extent to which they predict strictly defined, CIDI-based MDD in independent cohorts. We carried out an out-of-sample prediction analysis using data from the MDD Working Group of the Psychiatric Genomics Consortium (PGC). We used 23 MDD cohorts from PGC29-MDD^5,52^, 20 of which recorded endorsement of DSM-5 criteria A for MDD from structured interviews (Supplemental Methods, Supplemental Table S21, Supplemental Figure S12). We constructed polygenic risk scores (PRS) on each definition of depression in UKBiobank (Methods) and examined their prediction in each of the PGC29-MDD cohorts. Of note, PRS from all definitions of depression in UKBiobank, whether minimally or strictly phenotyped, accounted for a small proportion of variation in disease status in PGC29-MDD (Supplemental Table S22). We observed the following features.

First, we found that using the full sample of GPpsy, its PRS performs best at predicting MDD status in independent cohorts from PGC29-MDD (Figure 7a, Nargelkerke’s r2 = 0.017, AUC = 0.56 at P value threshold of 0.05, Supplemental Figure S13). When equal sample sizes are used (randomly down-sampled to 50,000 and case prevalence of 0.15, Methods), GPpsy no longer performs best at predicting MDD status in PGC29-MDD cohorts (Figure 7b). Rather, PRS from the strictly defined CIDI-based MDD (LifetimeMDD) best predicts MDD disease status (Nargelkerke’s r2 = 0.0026, AUC = 0.52 at P value threshold of 0.05, Supplemental Figure S13).

**Figure7:**
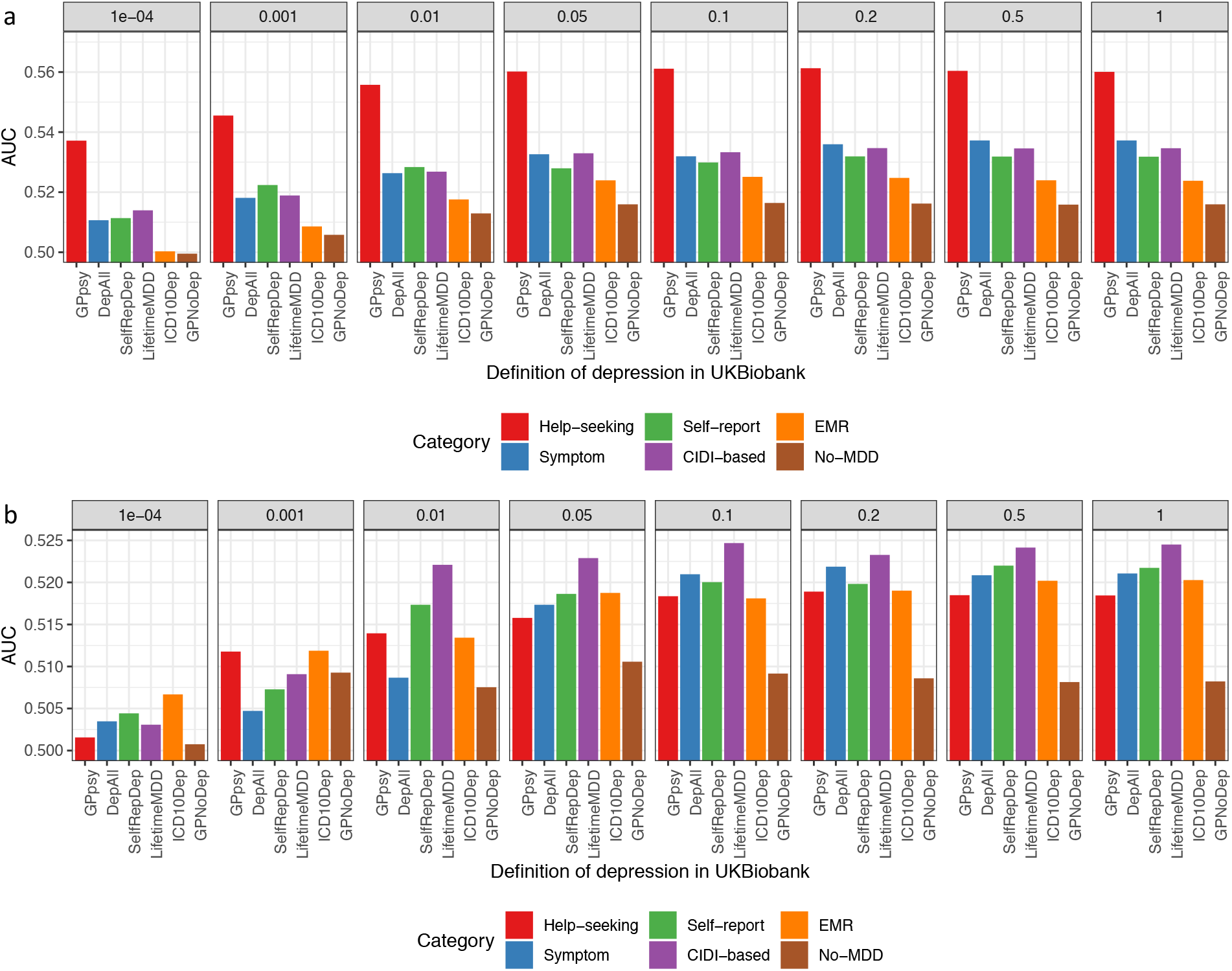
Out-of-sample prediction of MDD in PGC cohorts. a) This figure shows the area under the curve (AUC) of polygenic risk scores (PRS) calculated for each definition of depression in UKBiobank and MDD status indicated in 19 PGC29-MDD cohorts^5^, while controlling for cohort specific effects. PRS were calculated using effect sizes at independent (LD r^2^ < 0.1) SNPs passing P value thresholds 10^-4^, 0.001, 0.01, 0.05, 0.01, 0.2, 0.5 and 1 respectively, in GWAS performed on all definitions of depression in UKBiobank. b) This figure shows the same analysis performed on down-sampled data (7,500 cases, 42,500 controls) for each definition of depression.

Second, the higher prediction accuracy can be entirely explained by the larger sample size^55^ (Supplemental Methods, Supplemental Figure S14). The same was true for all other definitions of depression. We calculated the effective sample size needed for other definitions to have the same predictive power as GPpsy at its full sample size (113,260 cases, 219,362 controls, effective sample size = 298,677). For strictly defined LifetimeMDD, we would need an effective sample size of 129,106 (Supplemental Methods, Supplemental Figure S14).

Third, PRS from strictly defined LifetimeMDD predicts MDD disease status better in PGC29-MDD cohorts with higher percentage of cases fulfilling full DSM-5 symptom criteria for MDD (Supplemental Table S21, Supplemental Figure S15, Pearson r^2^ between AUC and percentage cases in PGC29-MDD cohorts fulfilling DSM-5 symptom criteria = 0.20, P = 0.05, at PRS P value threshold = 0.05). This is consistent with the interpretation that LifetimeMDD captures signal specific to MDD. We did not observe such a trend for GPpsy (Pearson r = 0.08, P = 0.23 at PRS P value = 0.05), or any other definition of depression (Supplemental Table S23), suggesting their lower specificity to MDD.

## Discussion

Though it is believed that the gain in statistical power from using larger sample sizes compensates for inaccuracies in minimal phenotyping^4,5,56–58^, our study demonstrates that the genetic architecture of minimal phenotyping definitions of depression is enriched for non-specific effects on MDD. Using a range of definitions of MDD in UKBiobank, from self-reported help-seeking to a full assessment of the DSM-5 criteria for MDD through self-reported symptoms from the MHQ, we made five key observations.

First, the heritabilities of depression defined by minimal phenotyping strategies are lower than MDD defined by full DSM-5 criteria using the CIDI questionnaire. Second, although there is substantial genetic correlation between definitions, there remain significant differences, indicating the presence of genetic effects unique to each definition. Of the genetic liability shared between depression defined by minimal phenotyping and CIDI-based MDD, a large proportion is not specific to MDD: for example, 66% of the genetic liability of LifetimeMDD due to GPpsy is shared with a non-MDD definition that excludes core depressive symptoms. Third, a larger proportion of genetic loci contributing to minimal phenotyping definitions of depression contribute to rG with other psychiatric conditions than those contributing to CIDI-based MDD, likely driving its enrichment of h^2^_SNP_ in CNS-specific genes and shared effects at GWAS loci with other psychiatric conditions. Fourth, all GWAS hits from minimal definition of depression GPpsy are shared with genetically correlated conditions such as neuroticism and smoking. There were too few GWAS hits for LifetimeMDD to allow us to test for their specificity. Finally, results from our analysis of the PRS derived from different definitions of depression support the view that those from minimal phenotyping definitions enable greater predictive power. However, consistent with findings from GWAS, this increase is due to large sample size instead of MDD-specific effects: LifetimeMDD predictions are correlated with the percentage of DSM-MDD cases in the PGC29-MDD cohorts, while GPpsy predictions are not. These results point to the non-specific nature of genetic factors identified in minimal phenotyping definitions of depression.

A number of factors need to be borne in mind when interpreting the above observations. Importantly, none of the definitions of depression in the UKBiobank were obtained from structured clinical interviews with an experienced rater (the gold standard for diagnosing MDD). The closest to that standard in UKBiobank is the online MHQ^18^, based on the Composite International Diagnostic Interview Short Form (CIDI-SF)^19^. There is evidence that self-reported diagnoses often over-estimate the prevalence of MDD based on interviews (for example in one population survey self-report questionnaire gave an estimate of 22.6%, compared to 8% based on interviews^59^). Agreements between self-report measures and clinician ratings are low^60^, as are test-retest reliabilities from clinicians in general practice (kappa coefficients of around 0.3 in DSM-5 field trials^61^. This is in contrast to high inter-rater reliability between trained clinicians following diagnostic manuals (kappa coefficients of around 0.7 in both DSM-3^62^ and DSM-4^63^ field trials).

The lack of CNS h^2^_SNP_ enrichment in strictly-defined MDD, although present in published studies and minimal phenotyping definitions of depression, may seem counterintuitive. Many factors influence the results of this analysis, including statistical power. Nevertheless, the non-specificity of CNS enrichment to MDD precludes its use for validation of any definition of depression. Our assumptions of a CNS involvement in MDD, however widely-accepted or reasonable, is neither sufficient nor valid as evidence that any particular definition of depression better represents MDD, or captures the biological mechanisms behind MDD.

Our results suggest that self-reported diagnoses using a CIDI-SF or other diagnostic questionnaires with full DSM-5 criteria lie on the same genetic liability continuum as MDD. This would argue that MDD cases identified through self-report means, using a full diagnostic questionnaire will be enriched for more strictly defined forms, with the consequence that results from genetic analysis will include loci that contribute to strictly defined MDD disease risk^64,65^.

In contrast, we show that minimal definitions of MDD do not simply include cases with lower genetic liability to MDD, and that PRS derived from minimal definitions are poor predictors of strictly defined MDD in independent samples. While it is true there is a high degree of genetic correlation between MDD and depressive symptoms (rG = 0.7, implying roughly rG^2^ = 49% of genetic factors contributing to liability, or genetic variance, of the former is attributable to that of the latter)^26^, it is also true that there is an even higher degree of sharing between depressive symptoms and other traits such as neuroticism (rG = 0.79-0.94, implying roughly rG^2^ = 62%-88% of genetic variance of the former is attributable to that of the latter, especially if both were assayed at a single time point^66^). In other words, the correlation is driven by nonspecific genetic risk factors. This is consistent with a recent study of three large twin cohorts, which asked if a combination of MDD, depressive symptoms and neuroticism is able to capture all genetic liability of MDD^67^. That study showed that 65% of the genetic effects contributing to MDD are specific, and minimally defined depression (inclusive of MDD, depressive symptoms and neuroticism) can index only around one-third of the genetic liability to MDD.

As expected from our analysis of SNP-heritabilities, genetic correlations, PRS predictions and GWAS loci, genetic findings from minimal definitions of MDD are not specific to MDD. This has important implications for downstream investigations. The biological information from a minimal definition of MDD in terms of genes and pathways discovered may be informative about mental ill health in general, but not MDD in particular. One interpretation is that the characterization of genetic loci with such non-specific effects will still advance understanding of the biology of psychiatric disorders and their treatment^5,56^. A recent report on genetic analyses of subjective well-being, depressive symptoms and neuroticism identified a high degree of sharing between genetic liabilities to the three measures, and use the “quasi-replication” of GWAS loci between depressive symptoms and neuroticism as validation of their functional significance^66^. An alternative view is that these loci reflect the ways in which depressive symptoms can develop as secondary effects, including through susceptibility to adverse life events^68^, personality types^28^, and use or exposure to psychoactive agents like cigarette smoking^69,70^. In which case, while useful for understanding the basis of mental ill health, they are not informative about the genetic etiology of MDD, and are not useful for developing disease specific treatment. Indeed one implication of our findings is that in conjunction with GWAS of strictly defined MDD, minimal phenotyping will be most useful in determining which loci to not prioritize for follow-up functional analyses.

Our findings indicate the need for ways to assess symptoms for diagnosing MDD with specificity and at scale. Fast and accurate diagnostic methods that use a limited number of questionnaire items are becoming available: for example, computerized adaptive diagnostic screening may be as effective for the diagnosis of MDD as an hour-long face-to-face clinician diagnostic interview^71^. Furthermore, there are ongoing attempts to convert behavioural health tracking data from phones or wearable devices into diagnostic information^72^. If successful, these attempts may lead to a dramatic expansion in our ability to collect data appropriate for psychiatric genetics. The need for the development and implementation of such novel approaches, rather than reliance on minimal phenotyping, is apparent from the genetic analyses we have carried out. We expect that combining genetic and novel phenotyping approaches could be a powerful solution to the problem of acquiring high quality psychiatric phenotypes at scale and a low cost.

## Methods

### Control for population structure

We performed principal component analysis (PCA) on directly genotyped SNPs from samples in UKBiobank and used PCs as covariates in all our analyses to control for population structure. From the array genotype data, we first removed all samples who did not pass QC, leaving 337,198 White-British, unrelated samples. We then removed SNPs not included in the phasing and imputation and retained those with minor allele frequencies (MAF) >= 0·1%, and P value for violation of Hardy-Weinberg equilibrium > 10^-6^, leaving 593,300 SNPs. We then removed 20,567 SNPs that are in known structural variants (SVs) and the major histocompatibility complex (MHC)^40^ as recommended by UKBiobank^73^, leaving 572,733 SNPs that we consistently use for all analyses. Of these, 334,702 are common (MAF > 5%), and from these common SNPs we further filtered based on missingness <0·02 and pairwise LD r^2^ < 0·1 with SNPs in a sliding window of 1000 SNPs to obtain 68,619 LD-pruned SNPs for computing PCs using flashPCA^74^. We obtained 20 PCs, their eigenvalues, loadings and variance explained, and consistently use these PCs as covariates for all our genetic analyses. We note that control over population structure over the SVs and MHC is minimal, and we explore the impact of this in Supplemental Methods, Supplemental Table S4-6 and Supplemental Figure S2.

### Imputed genotype filtering

We performed stringent filtering on imputed variants (version 2) used for GWAS in this study, removing variants not among the 33,619,058 variant sites in the Haplotype Reference Consortium^75^ (HRC) panel, then removing all insertions and deletions (INDELs) and multi-allelic SNPs. We hard-called genotypes from imputed dosages at 8,968,715 biallelic SNPs with imputation INFO score greater than 0·9, MAF greater than 0·1%, and P value for violation of Hardy-Weinberg equilibrium > 10^-6^, with a genotype probability threshold of 0·9 (anything below would be considered missing). Of these, 5,276,842 SNPs are common (MAF > 5%). We consistently use these SNPs for all analyses in this study.

### Genome-wide associations

To obtain and access the difference between odds ratios of associations in different definitions of depression in UKBiobank, as well as smoking (data field 20160) and neuroticism (data field 20127), we perform logistic regression (or linear regression with --standard-beta for neuroticism) on all 5,276,842 common SNPs (MAF > 5% in all 337,198 White-British, unrelated samples) in PLINK^76^ (version 1·9) with 20 PCs and genotyping array as covariates. We report all associations with P values smaller than 5 x 10^-8^ as genome-wide significant. We indicated the SNPs in SVs and the MHC in tables of top hits as well as all manhattan plots as hollow points instead of solid points due to lack of control for population structure in these regions, and do not interpret potential causal effects in these regions (Supplemental Tables S10-11,16-17, Supplemental Figure S5,9).

### Estimation of SNP-heritability and genetic correlation among definitions of MDD

All estimates of h^2^_SNP_ we refer to in the main text of this paper are computed with the phenotype-correlation-genotype-correlation (PCGC)^77^ approach implemented with PCGCs^23^, using 5,276,842 common SNPs (MAF > 5% in all 337,198 White-British, unrelated samples). We used LD scores at SNPs computed with LDSC^37^ in 10,000 random samples drawn from the White-British samples in UKBiobank as LD reference, and MAF at all 5,276,842 common SNPs in all 337,198 White-British samples as MAF reference. As covariates we used genotyping array and 20 PCs computed using samples in each definition of MDD with flashPCA^74^, and supplied the eigenvalues of the 20 PCs to PCGCs as deflation factors to correct for deflation in sum of squared kinship coefficients when regressing PCs out of genotype vectors. Where we stratified each definition of MDD in UKBiobank into two strata by risk factors such as sex (Supplemental Methods), we computed specific PCs for each definition and strata. We use summary statistics generated by PCGCs for each definition of MDD in UKBiobank to estimate its h^2^_SNP_, and summary statistics of pairs of definitions of MDD to estimate their genetic correlation. We detail the methods and results from other approaches we used to compare our estimates from PCGCs with in Supplemental Methods and Supplemental Table S13.

### Estimation of genetic correlation between definitions of MDD and other conditions

We obtained summary statistics for other psychiatric conditions from previous GWAS studies as described in Supplemental Table S1. We also performed GWAS on smoking and neuroticism in UKBiobank (Supplemental Table S15-16, Supplemental Figure S10) to generate their association summary statistics. We estimated the genetic correlation between definitions of MDD in UKBiobank with each of these conditions with LDSC^46^, with a LD reference panel generated with EUR individuals from 1000 Genomes^78^, using SNPs with association statistics available in both conditions in each instance. To obtain regional genetic correlation, we partitioned the genome into 1703 independent loci^79^ and estimated regional genetic correlation at each locus with rho-HESS^49^, using a LD reference panel generated with EUR individuals from 1000 Genomes^78^. We estimated standard errors for each regional genetic correlation and the cumulative genetic correlations across the genome using a jackknife approach implemented in HESS^39^. To estimate the percentage of loci needed to explain 90% of the total genetic correlation for both LifetimeMDD and GPpsy with all conditions, we ranked all independent loci by their absolute genetic correlation (such that a locus with large negative genetic correlation counts just as much as that with a large positive genetic correlation), and calculated the percentage of all genomic loci that would contribute 90% of the total genetic covariance.

### Enrichment of SNP-heritability in genes specifically expressed in tissues

We estimate the enrichment of h^2^_SNP_ in genes specifically expressed in 44 tissues in the Genotype–Tissue Expression (GTEx)^50^ project using the partitioned h^2^_SNP_ framework in LDSC-SEG^49^, and a LD reference panel generated with EUR individuals from 1000 Genomes^78^. We first obtained tissue specific gene expression annotations in GTEx tissues from LDSC-SEG, then estimated the enrichment of h^2^_SNP_ in annotations that corresponded to each of the tissues together with 52 annotations in the baseline model^80^ which we also obtain from LDSC-SEG. We report the P value of the one-sided test of enrichment of h^2^_SNP_ (positive regression coefficient for the tissue-specific annotation conditioning on other baseline annotations) in genes specifically expressed in each tissue against the baseline.

### Meta-analysis between PGC29 and LifetimeMDD

We obtained summary statistics of meta-analysis of GWAS on 29 MDD cohorts in the Psychiatric Genomics Consortium (PGC29, N = 42,455) through the MDD Working Group of the Psychiatric Genomics Consortium (PGC-MDD), reported in Wray et al 2018^5^. Prior to meta-analysis with LifetimeMDD (N = 67,171), we removed all INDELs, as well as SNPs with MAF < 5% and imputation INFO score < 0.9, leaving 5,828,030 SNPs. We use this set of summary statistics for estimation of h^2^_SNP_ with LDSC, as well as for enrichment analyses in LDSC-SEG^51^. We then performed a meta-analysis between the filtered summary statistics of PGC29 with those of LifetimeMDD using METAL^81^ using the SCHEME STDERR. We removed those SNPs at which no meta-analysis was performed due to their absence in either dataset. The final meta-analysed data contained summary statistics at 4,693,521 SNPs for a total sample size N = 109,626, which we used for estimation of h^2^_SNP_ with LDSC, as well as for enrichment analyses in LDSC-SEG.

### Out of sample predictions of MDD

We carried out an out-of-sample prediction using individual level genotype and phenotype data from MDD cohorts among PGC29 in Wray et al 2018^5^. We obtained permissions from PGC-MDD for 20 cohorts with sample sizes (both cases and controls) greater than 500, among which 17 recorded endorsement of DSM-5 criteria A for MDD, using the same criteria we did for CIDI-based definition LifetimeMDD (Supplemental Methods, Supplemental Table S20). To obtain polygenic risk scores (PRS) for each definintion of depression in UKBiobank without confounding from power differences due to sample sizes, we downsampled all definitions to a constant sample size of 50,000 and case prevalence of 0.15 through randomly sampling 7,500 cases from all cases, and 42,500 controls from all controls in each definition. We then obtained PRS using GWAS on the down-sampled data: for each definition of depression in UKBiobank, we obtained SNPs with P values of associations below 8 thresholds (P < 10^-4^, 0.001, 0.01, 0.05, 0.1, 0.2, 0.5 and 1), and LD clumped them (LD r^2^ < 0.1) to obtain independent SNPs. We then used independent SNPs for each threshold and each definition of depression in UKBiobank to construct polygenic risk scores (PRS) and predict MDD status in the 20 PGC cohorts using the Ricopili pipeline^82^. We obtained Nagelkerke’s r^2^ between the PRS and MDD status, AUC of the prediction, and variance of MDD status explained by the PRS for each cohort. We also obtained the same measures for MDD status pulling data from all cohorts, controlling for cohort differences by including it as a covariate.

## Author Contributions

NC and JF designed the study. NC and JR performed the analyses. NC and JF obtained the data from the UKBiobank Resource. MJA, TFMA, GB, EMB, TKC, AJF, HJG, SPH, DFL, CML, GL, NGM, YM, OM, BMM, BWJHP, RHP, GP, JBP, MP, JS, JWS, FS, HT, RU, SVA, AV, MMW and all investigators from the MDD Working Group of the PGC contributed data from the PGC. NC, KSK and JF interpreted the results and wrote the manuscript.

## Ethical approval

This research was conducted under the ethical approval from the UKBiobank Resource under application no. 28709.

## Acknowledgements

We would like to thank Omer Weissbrod, Andy Dahl, Huwenbo Shi and Verena Zuber for insightful discussions. Na Cai is supported by the ESPOD Fellowship from the European Bioinformatics (EMBL-EBI) and Wellcome Sanger Institute. Alexander Viktorin is supported by the Swedish Brain Foundation. In the last three years Myrna Weissman has received research funds from NIMH, Templeton Foundation and the Sackler Foundation and has received royalties for publications of books on interpersonal psychotherapy from Perseus Press, Oxford University Press, on other topics from the American Psychiatric Association Press and royalties on the social adjustment scale from Multihealth Systems. The CoLaus|PsyCoLaus study was and is supported by research grants from GlaxoSmithKline, the Faculty of Biology and Medicine of Lausanne, and the Swiss National Science Foundation (grants 3200B0–105993, 3200B0-118308, 33CSCO-122661, 33CS30-139468 and 33CS30-148401). The PGC has received major funding from the US National Institute of Mental Health and the US National Institute of Drug Abuse (U01 MH109528 and U01 MH1095320). This research was conducted using the UKBiobank Resource under application no. 28709, and with the support and collaboration from all investigators who comprise the MDD Working Group of the PGC (full list in Supplemental Materials). We are greatly indebted to the hundreds of thousands of individuals who have shared their life experiences with the UKBiobank and PGC investigators.

## Declaration of interests

Cathryn M Lewis is on the scientific advisory board of Myriad Neuroscience. Hans J Grabe has received travel grants and speakers honoraria from Fresenius Medical Care, Neuraxpharm and Janssen Cilag as well as research funding from Fresenius Medical Care.

## Data availability

Genotype and phenotype data used in this study are from the full release (imputation version 2) of the UKBiobank Resource obtained under application no. 28709. We used publicly available summary statistics from other studies downloadable from the website of Psychiatric Genomics Consortium (https://www.med.unc.edu/pgc/results-and-downloads), and the references for which can be found in Supplemental Table S1. We also referenced the 2011 Census aggregate data from the UK Data Service (http://dx.doi.org/10.5257/census/aggregate-2011-2).

