## Supplemental Material for "Minimal phenotyping yields GWAS hits of reduced specificity for major depression"

**Members of the MDD Working Group of the Psychiatric Genomics Consortium 3**

**Supplemental Methods 8**

Other studies referenced in this paper 8

Sample filtering 8

Minimal phenotyping definitions of depression in UKBiobank 8

Electronic health record-based definition of depression in UKBiobank 9

Strictly defined, CIDI-based definition of MDD in UKBiobank 9

The non-MDD definition of depression 10

Comparison of definitions of depression in UKBiobank with those from previous GWAS 11

Prevalence of definitions of depression in UKBiobank 13

Control of population structure in SNP-heritability estimates 14

Measure of lifetime trauma 14

Epidemiological analysis of risk factors for MDD 15

Potential confounding from self-ascertainment bias 16

Case enrichment and use of "clean" controls in GWAS 17

Comparison of methods of SNP-heritability and genetic correlation estimation 18

Stratification of definitions of depression by environmental risk factors 20

Simulations of effects of misdiagnosis and misclassification on SNP-heritability estimates 20

LDSC-SEG analysis of tissue-specific enrichment of $h^2_{SNP}$ 22

Out-of-sample prediction of MDD in PGC cohorts 23

Effect of Sample size and specificity on prediction power 24

**Supplemental Figures 26**

Supplemental Figure S1: Symptoms of CIDI-based MDD LifetimeMDD in UKBiobank 26

Supplemental Figure S2: Overlap between definitions of MDD in UKBiobank 27

Supplemental Figure S3: Effects of population structure and MHC/SVs 28

Supplemental Figure S4: Effect of MHQ participation on definitions of depression 30

Supplemental Figure S5: GWAS on definitions of depression using "clean" controls 31

Supplemental Figure S6: LDSC estimates of heritability and genetic correlation 32

Supplemental Figure S7: Simulations of misdiagnosis and misclassification 34

Supplemental Figure S8: Simulations of misclassification at different heritabilities 35

Supplemental Figure S9: GWAS on neuroticism and smoking in UKBiobank 36

Supplemental Figure S10: LDSC-SEG analysis of tissue-specific enrichment of $h^2_{SNP}$ 37

Supplemental Figure S11: GWAS hits from 23andMe are not specific to MDD 38
Supplemental Figure S12: Symptoms of MDD cases in PGC cohorts 39
Supplemental Figure S13: Out-of-sample prediction in PGC cohorts 40
Supplemental Figure S14: Relationship between effective sample size and prediction accuracy 41
Supplemental Figure S15: Prediction accuracy in cohorts with different percentage DSM MDD cases 42

**Supplemental Tables 43**

Supplemental Table S1: Other studies referenced in this paper 44
Supplemental Table S2: CIDI-based criteria for lifetime and current MDD 46
Supplemental Table S3: Characterizing no-MDD definition GPNoDep 47
Supplemental Table S4: Number of samples by assessment centre 49
Supplemental Table S6: Correction for population in assessment centres 51
Supplemental Table S7: Correction for age and sex 52
Supplemental Table S8: GWAS hits for participation in the MHQ (adapted from Adams et al 2019) 54
Supplemental Table S9: Assessing casusal relationship between MHQ participation and definitions of depression 56
Supplemental Table S10: Genome-wide significant loci in GWAS for definitions of MDD 58
Supplemental Table S11: Genome-wide significant loci in GWAS for definitions of MDD using “clean” controls 60
Supplemental Table S12: Derivation of lifetime trauma score 62
Supplemental Table S13: Heritability estimates of definitions of MDD from different methods 64
Supplemental Table S14: Heritability of definitions of MDD stratified by risk factors 69
Supplemental Table S15: Genetic correlation between definitions of depression in UKBiobank 71
Supplemental Table S16: Genome-wide significant loci in GWAS for smoking 73
Supplemental Table S17: Genome-wide significant loci in GWAS for neuroticism 76
Supplemental Table S18: Genetic correlation between definitions of MDD and other conditions 79
Supplemental Table S19: Effects of GWAS loci from help-seeking definitions on other definitions of MDD in UKBiobank and psychiatric conditions 85
Supplemental Table S20: Effects of GWAS loci from minimal phenotyping definition of MDD in 23andMe on definitions of MDD in UKBiobank and psychiatric conditions 89
Supplemental Table S21: PGC cohorts for out-of-sample predictions of MDD 91
Supplemental Table S22: Out-of-sample prediction of MDD in PGC cohorts 97
Supplemental Table S23: Correlation between prediction Naglekerke’s r2 and percentage DSM-MDD cases in PGC cohorts 98

**Members of the MDD Working Group of the Psychiatric Genomics Consortium**

Naomi R Wray 1,2, Stephan Ripke 3,4,5, Manuel Mattheisen 6,7,8, Maciej Trzaskowski 1, Enda M Byrne 1, Abdel Abdellaoui 9, Mark J Adams 10, Esben Agerbo 11,12,13, Tracy M Air 14, Till F M Andlauer 15,16, Silviu-Alin Bacanu 17, Marie Bækvad-Hansen 13,18, Aartjan T F Beekman 19, Tim B Bigdeli 17,20, Elisabeth B Binder 15, 21, Julien Bryois 22, Henriette N Buttenschøn 13,23,24, Jonas Bybjerg-Grauholm 13,18, Na Cai 25,26, Enrique Castelao 27, Jane Hvarregaard Christensen 8,13,24, Toni-Kim Clarke 10, Jonathan R I Coleman 28, Lucía Colodro-Conde 29, Baptiste Couvy-Duchesne 2, 30, Nick Craddock 31, Gregory E Crawford 32, 33, Gail Davies 34, Ian J Deary 34, Franziska Degenhardt 35, Eske M Derks 29, Nese Direk 36,37, Conor V Dolan 9, Erin C Dunn 38,39,40, Thalia C Eley 28, Valentina Escott-Price 41, Farnush Farhadi Hassan Kiadeh 42, Hilary K Finucane 43,44, Jerome C Foo 45, Andreas J Forstner 35,46,47,48, Josef Frank 45, Héléna A Gaspar 28, Michael Gill 49, Fernando S Goes 50, Scott D Gordon 29, Jakob Grove 8,13,24,51, Lynsey S Hall 10,52, Christine Søholm Hansen 13,18, Thomas F Hansen 53,54,55, Stefan Herms 35,47, Ian B Hickie 56, Per Hoffmann 35,47, Georg Homuth 57, Carsten Horn 58, Jouke-Jan Hottenga 9, David M Hougaard 13,18, David M Howard 10,28, Marcus Ising 59, Rick Jansen 19, Ian Jones 60, Lisa A Jones 61, Eric Jorgenson 62, James A Knowles 63, Isaac S Kohane 64,65,66, Julia Kraft 4, Warren W. Kretzschmar 67, Zoltán Kutalik 68,69, Yihan Li 67, Penelope A Lind 29, Donald J MacIntyre 70,71, Dean F MacKinnon 50, Robert M Maier 2, Wolfgang Maier 72, Jonathan Marchini 73, Hamdi Mbarek 9, Patrick McGrath 74, Peter McGuffin 28, Sarah E Medland 29, Divya Mehta 2,75, Christel M Middeldorp 9,76,77, Evelin Mihailov 78, Yuri Milaneschi 19, Lili Milani 78, Francis M Mondimore 50, Grant W Montgomery 1, Sara Mostafavi 79,80, Niamh Mullins 28, Matthias Nauck 81,82, Bernard Ng 80, Michel G Nivard 9, Dale R Nyholt 83, Paul F O'Reilly 28, Hogni Oskarsson 84, Michael J Owen 60, Jodie N Painter 29, Carsten Bøcker Pedersen 11,12,13, Marianne Giørtz Pedersen 11,12,13, Roseann E Peterson 17, 85, Erik Pettersson 22, Wouter J Peyrot 19, Giorgio Pistis 27, Danielle Posthuma 86,87, Jorge A Quiroz 88, Per Qvist 8,13,24, John P Rice 89, Brien P. Riley 17, Margarita Rivera 28,90, Saira Saeed Mirza 36, Robert Schoevers 91, Eva C Schulte 92,93, Ling Shen 62, Jianxin Shi 94, Stanley I Shyn 95, Engilbert Sigurdsson 96, Grant C B Sinnamon 97, Johannes H Smit 19, Daniel J Smith 98, Hreinn Stefansson 99, Stacy Steinberg 99, Fabian Streit 45, Jana Strohmaier 45, Katherine E Tansey 100, Henning Teismann 101, Alexander Teumer 102, Wesley Thompson 13,54,103,104, Pippa A Thomson 105, Thorgeir E Thorgeirsson 99, Matthew Traylor 106, Jens Treutlein 45, Vassily Trubetskoy 4, Andrés G Uitterlinden 107, Daniel Umbricht 108, Sandra Van der Auwera 109, Albert M van Hemert 110, Alexander Viktorin 22, Peter M Visscher 1,2, Yunpeng Wang 13,54,104, Bradley T. Webb 111, Shantel Marie Weinsheimer 13,54, Jürgen Wellmann 101, Gonneke Willemsen 9, Stephanie H Witt 45, Yang Wu 1, Hualin S Xi 112, Jian Yang 2,113, Futao Zhang 1, Volker Arolt 114, Bernhard T Baune 115,116,117, Klaus Berger 101, Dorret I Boomsma 9, Sven Cichon 35,47,118,119, Udo Dannlowski 114, EJC de Geus 9,120, J Raymond DePaulo 50, Enrico Domenici 121, Katharina Domschke 122,123, Tõnu Esko 5,78, Hans J Grabe 109, Steven P Hamilton 124, Caroline Hayward 125, Andrew C Heath 89, Kenneth S Kendler 17, Stefan Kloiber 59,126,127, Glyn Lewis 128, Qingqin S Li 129, Susanne Lucae 59, Pamela AF Madden 89, Patrik K Magnusson 22, Nicholas G Martin 29, Andrew M McIntosh 10,34, Andres Metspalu 78,130, Ole Mors 13,131, Preben Bo Mortensen 11,12,13,24, Bertram Müller-Myhsok 15,132,133, Merete Nordentoft 13,134, Markus M Nöthen 35, Michael C O'Donovan 60, Sara A Paciga 135, Nancy L Pedersen 22, Brenda WJH Penninx 19, Roy H Perlis 38,136, David J Porteous 105, James B Potash 137, Martin Preisig 27, Marcella Rietschel 45, Catherine Schaefer 62, Thomas G Schulze 45,93,138,139,140, Jordan W Smoller 38,39,40, Kari Stefansson 99,141, Henning Tiemeier 36,142,143, Rudolf Uher 144, Henry Völzke 102, Myrna M Weissman 74,145, Thomas Werge 13,54,146, Cathryn M Lewis 28,147, Douglas F Levinson 148, Gerome Breen 28,149, Anders D Børglum 8,13,24

1. Institute for Molecular Bioscience, The University of Queensland, Brisbane, QLD, AU
2. Queensland Brain Institute, The University of Queensland, Brisbane, QLD, AU
3. Analytic and Translational Genetics Unit, Massachusetts General Hospital, Boston, MA, US
4. Department of Psychiatry and Psychotherapy, Universitätsmedizin Berlin Campus Charité Mitte, Berlin, DE
5. Medical and Population Genetics, Broad Institute, Cambridge, MA, US
6. Department of Psychiatry, Psychosomatics and Psychotherapy, University of Wurzburg, Wurzburg, DE
7. Centre for Psychiatry Research, Department of Clinical Neuroscience, Karolinska Institutet, Stockholm, SE

8. Department of Biomedicine, Aarhus University, Aarhus, DK
9. Dept of Biological Psychology & EMGO+ Institute for Health and Care Research, Vrije Universiteit Amsterdam, Amsterdam, NL
10. Division of Psychiatry, University of Edinburgh, Edinburgh, GB
11. Centre for Integrated Register-based Research, Aarhus University, Aarhus, DK
12. National Centre for Register-Based Research, Aarhus University, Aarhus, DK
13. iPSYCH, The Lundbeck Foundation Initiative for Integrative Psychiatric Research,, DK
14. Discipline of Psychiatry, University of Adelaide, Adelaide, SA, AU
15. Department of Translational Research in Psychiatry, Max Planck Institute of Psychiatry, Munich, DE
16. Department of Neurology, Klinikum rechts der Isar, Technical University of Munich, Munich, DE
17. Department of Psychiatry, Virginia Commonwealth University, Richmond, VA, US
18. Center for Neonatal Screening, Department for Congenital Disorders, Statens Serum Institut, Copenhagen, DK
19. Department of Psychiatry, Vrije Universiteit Medical Center and GGZ inGeest, Amsterdam, NL
20. Virginia Institute for Psychiatric and Behavior Genetics, Richmond, VA, US
21. Department of Psychiatry and Behavioral Sciences, Emory University School of Medicine, Atlanta, GA, US
22. Department of Medical Epidemiology and Biostatistics, Karolinska Institutet, Stockholm, SE
23. Department of Clinical Medicine, Translational Neuropsychiatry Unit, Aarhus University, Aarhus, DK
24. iSEQ, Centre for Integrative Sequencing, Aarhus University, Aarhus, DK
25. Human Genetics, Wellcome Trust Sanger Institute, Cambridge, GB
26. Statistical genomics and systems genetics, European Bioinformatics Institute (EMBL-EBI), Cambridge, GB
27. Department of Psychiatry, Lausanne University Hospital and University of Lausanne, Lausanne, CH
28. Social Genetic and Developmental Psychiatry Centre, King's College London, London, GB
29. Genetics and Computational Biology, QIMR Berghofer Medical Research Institute, Brisbane, QLD, AU
30. Centre for Advanced Imaging, The University of Queensland, Brisbane, QLD, AU
31. Psychological Medicine, Cardiff University, Cardiff, GB
32. Center for Genomic and Computational Biology, Duke University, Durham, NC, US
33. Department of Pediatrics, Division of Medical Genetics, Duke University, Durham, NC, US
34. Centre for Cognitive Ageing and Cognitive Epidemiology, University of Edinburgh, Edinburgh, GB
35. Institute of Human Genetics, School of Medicine & University Hospital Bonn, University of Bonn, Bonn, DE
36. Epidemiology, Erasmus MC, Rotterdam, Zuid-Holland, NL
37. Psychiatry, Dokuz Eylul University School Of Medicine, Izmir, TR
38. Department of Psychiatry, Massachusetts General Hospital, Boston, MA, US
39. Psychiatric and Neurodevelopmental Genetics Unit (PNGU), Massachusetts General Hospital, Boston, MA, US
40. Stanley Center for Psychiatric Research, Broad Institute, Cambridge, MA, US
41. Neuroscience and Mental Health, Cardiff University, Cardiff, GB
42. Bioinformatics, University of British Columbia, Vancouver, BC, CA
43. Department of Epidemiology, Harvard T.H. Chan School of Public Health, Boston, MA, US
44. Department of Mathematics, Massachusetts Institute of Technology, Cambridge, MA, US
45. Department of Genetic Epidemiology in Psychiatry, Central Institute of Mental Health, Medical Faculty Mannheim, Heidelberg University, Mannheim, Baden-Württemberg, DE
46. Department of Psychiatry (UPK), University of Basel, Basel, CH
47. Department of Biomedicine, University of Basel, Basel, CH
48. Centre for Human Genetics, University of Marburg, Marburg, DE
49. Department of Psychiatry, Trinity College Dublin, Dublin, IE
50. Psychiatry & Behavioral Sciences, Johns Hopkins University, Baltimore, MD, US
51. Bioinformatics Research Centre, Aarhus University, Aarhus, DK
52. Institute of Genetic Medicine, Newcastle University, Newcastle upon Tyne, GB
53. Danish Headache Centre, Department of Neurology, Rigshospitalet, Glostrup, DK

54. Institute of Biological Psychiatry, Mental Health Center Sct. Hans, Mental Health Services Capital Region of Denmark, Copenhagen, DK
55. iPSYCH, The Lundbeck Foundation Initiative for Psychiatric Research, Copenhagen, DK
56. Brain and Mind Centre, University of Sydney, Sydney, NSW, AU
57. Interfaculty Institute for Genetics and Functional Genomics, Department of Functional Genomics, University Medicine and Ernst Moritz Arndt University Greifswald, Greifswald, Mecklenburg-Vorpommern, DE
58. Roche Pharmaceutical Research and Early Development, Pharmaceutical Sciences, Roche Innovation Center Basel, F. Hoffmann-La Roche Ltd, Basel, CH
59. Max Planck Institute of Psychiatry, Munich, DE
60. MRC Centre for Neuropsychiatric Genetics and Genomics, Cardiff University, Cardiff, GB
61. Department of Psychological Medicine, University of Worcester, Worcester, GB
62. Division of Research, Kaiser Permanente Northern California, Oakland, CA, US
63. Psychiatry & The Behavioral Sciences, University of Southern California, Los Angeles, CA, US
64. Department of Biomedical Informatics, Harvard Medical School, Boston, MA, US
65. Department of Medicine, Brigham and Women's Hospital, Boston, MA, US
66. Informatics Program, Boston Children's Hospital, Boston, MA, US
67. Wellcome Trust Centre for Human Genetics, University of Oxford, Oxford, GB
68. Institute of Social and Preventive Medicine (IUMSP), University Hospital of Lausanne, Lausanne, VD, CH
69. Swiss Institute of Bioinformatics, Lausanne, VD, CH
70. Division of Psychiatry, Centre for Clinical Brain Sciences, University of Edinburgh, Edinburgh, GB
71. Mental Health, NHS 24. Glasgow, GB
72. Department of Psychiatry and Psychotherapy, University of Bonn, Bonn, DE
73. Statistics, University of Oxford, Oxford, GB
74. Department of Psychiatry, Columbia University, Vagelos College of Physicians and Surgeons, New York, NY, US
75. School of Psychology and Counseling, Queensland University of Technology, Brisbane, QLD, AU
76. Child and Youth Mental Health Service, Children's Health Queensland Hospital and Health Service, South Brisbane, QLD, AU
77. Child Health Research Centre, University of Queensland, Brisbane, QLD, AU
78. Estonian Genome Center, University of Tartu, Tartu, EE
79. Medical Genetics, University of British Columbia, Vancouver, BC, CA
80. Statistics, University of British Columbia, Vancouver, BC, CA
81. DZHK (German Centre for Cardiovascular Research), Partner Site Greifswald, University Medicine, University Medicine Greifswald, Greifswald, Mecklenburg-Vorpommern, DE
82. Institute of Clinical Chemistry and Laboratory Medicine, University Medicine Greifswald, Greifswald, Mecklenburg-Vorpommern, DE
83. Institute of Health and Biomedical Innovation, Queensland University of Technology, Brisbane, QLD, AU
84. Humus, Reykjavik, IS
85. Virginia Institute for Psychiatric & Behavioral Genetics, Virginia Commonwealth University, Richmond, VA, US
86. Clinical Genetics, Vrije Universiteit Medical Center, Amsterdam, NL
87. Complex Trait Genetics, Vrije Universiteit Amsterdam, Amsterdam, NL
88. Solid Biosciences, Boston, MA, US
89. Department of Psychiatry, Washington University in Saint Louis School of Medicine, Saint Louis, MO, US
90. Department of Biochemistry and Molecular Biology II, Institute of Neurosciences, Center for Biomedical Research, University of Granada, Granada, ES
91. Department of Psychiatry, University of Groningen, University Medical Center Groningen, Groningen, NL
92. Department of Psychiatry and Psychotherapy, University Hospital, Ludwig Maximilian University Munich, Munich, DE
93. Institute of Psychiatric Phenomics and Genomics (IPPG), University Hospital, Ludwig Maximilian University Munich, Munich, DE
94. Division of Cancer Epidemiology and Genetics, National Cancer Institute, Bethesda, MD, US

95. Behavioral Health Services, Kaiser Permanente Washington, Seattle, WA, US
96. Faculty of Medicine, Department of Psychiatry, University of Iceland, Reykjavik, IS
97. School of Medicine and Dentistry, James Cook University, Townsville, QLD, AU
98. Institute of Health and Wellbeing, University of Glasgow, Glasgow, GB
99. deCODE Genetics / Amgen, Reykjavik, IS
100. College of Biomedical and Life Sciences, Cardiff University, Cardiff, GB
101. Institute of Epidemiology and Social Medicine, University of Münster, Münster, Nordrhein-Westfalen, DE
102. Institute for Community Medicine, University Medicine Greifswald, Greifswald, Mecklenburg-Vorpommern, DE
103. Department of Psychiatry, University of California, San Diego, San Diego, CA, US
104. KG Jebsen Centre for Psychosis Research, Norway Division of Mental Health and Addiction, Oslo University Hospital, Oslo, NO
105. Medical Genetics Section, CGEM, IGMM, University of Edinburgh, Edinburgh, GB
106. Clinical Neurosciences, University of Cambridge, Cambridge, GB
107. Internal Medicine, Erasmus MC, Rotterdam, Zuid-Holland, NL
108. Roche Pharmaceutical Research and Early Development, Neuroscience, Ophthalmology and Rare Diseases Discovery & Translational Medicine Area, Roche Innovation Center Basel, F. Hoffmann-La Roche Ltd, Basel, CH
109. Department of Psychiatry and Psychotherapy, University Medicine Greifswald, Greifswald, Mecklenburg-Vorpommern, DE
110. Department of Psychiatry, Leiden University Medical Center, Leiden, NL
111. Virginia Institute for Psychiatric & Behavioral Genetics, Virginia Commonwealth University, Richmond, VA, US
112. Computational Sciences Center of Emphasis, Pfizer Global Research and Development, Cambridge, MA, US
113. Institute for Molecular Bioscience; Queensland Brain Institute, The University of Queensland, Brisbane, QLD, AU
114. Department of Psychiatry, University of Münster, Münster, Nordrhein-Westfalen, DE
115. Department of Psychiatry, University of Münster, Münster, DE
116. Department of Psychiatry, Melbourne Medical School, University of Melbourne, Melbourne, AU
117. Florey Institute for Neuroscience and Mental Health, University of Melbourne, Melbourne, AU
118. Institute of Medical Genetics and Pathology, University Hospital Basel, University of Basel, Basel, CH
119. Institute of Neuroscience and Medicine (INM-1), Research Center Juelich, Juelich, DE
120. Amsterdam Public Health Institute, Vrije Universiteit Medical Center, Amsterdam, NL
121. Centre for Integrative Biology, Università degli Studi di Trento, Trento, Trentino-Alto Adige, IT
122. Department of Psychiatry and Psychotherapy, Medical Center - University of Freiburg, Faculty of Medicine, University of Freiburg, Freiburg, DE
123. Center for NeuroModulation, Faculty of Medicine, University of Freiburg, Freiburg, DE
124. Psychiatry, Kaiser Permanente Northern California, San Francisco, CA, US
125. Medical Research Council Human Genetics Unit, Institute of Genetics and Molecular Medicine, University of Edinburgh, Edinburgh, GB
126. Department of Psychiatry, University of Toronto, Toronto, ON, CA
127. Centre for Addiction and Mental Health, Toronto, ON, CA
128. Division of Psychiatry, University College London, London, GB
129. Neuroscience Therapeutic Area, Janssen Research and Development, LLC, Titusville, NJ, US
130. Institute of Molecular and Cell Biology, University of Tartu, Tartu, EE
131. Psychosis Research Unit, Aarhus University Hospital, Risskov, Aarhus, DK
132. Munich Cluster for Systems Neurology (SyNergy), Munich, DE
133. University of Liverpool, Liverpool, GB
134. Mental Health Center Copenhagen, Copenhagen Universtity Hospital, Copenhagen, DK
135. Human Genetics and Computational Biomedicine, Pfizer Global Research and Development, Groton, CT, US

136. Psychiatry, Harvard Medical School, Boston, MA, US
137. Psychiatry, University of Iowa, Iowa City, IA, US
138. Department of Psychiatry and Behavioral Sciences, Johns Hopkins University, Baltimore, MD, US
139. Department of Psychiatry and Psychotherapy, University Medical Center Göttingen, Goettingen, Niedersachsen, DE
140. Human Genetics Branch, NIMH Division of Intramural Research Programs, Bethesda, MD, US
141. Faculty of Medicine, University of Iceland, Reykjavik, IS
142. Child and Adolescent Psychiatry, Erasmus MC, Rotterdam, Zuid-Holland, NL
143. Psychiatry, Erasmus MC, Rotterdam, Zuid-Holland, NL
144. Psychiatry, Dalhousie University, Halifax, NS, CA
145. Division of Translational Epidemiology, New York State Psychiatric Institute, New York, NY, US
146. Department of Clinical Medicine, University of Copenhagen, Copenhagen, DK
147. Department of Medical & Molecular Genetics, King's College London, London, GB
148. Psychiatry & Behavioral Sciences, Stanford University, Stanford, CA, US
149. NIHR Maudsley Biomedical Research Centre, King's College London, London, GB
150. Genetics, University of North Carolina at Chapel Hill, Chapel Hill, NC, US
151. Psychiatry, University of North Carolina at Chapel Hill, Chapel Hill, NC, US

**Supplemental Methods**

**Other studies referenced in this paper**

We have summarised the other studies we reference in our paper, summary statistics of many of which are available on the Psychiatric Genomics Consortium website, in Supplemental Table S1. For each of the studies, we use only those SNPs with imputation INFO score above 0.9 and MAF > 5% in estimation of heritability, partitioned heritability, and genetic correlation.

**Sample filtering**

Of all 502,637 samples in UKBiobank full release, we performed the following QC steps to select the samples for use in our analyses. We first removed samples that were not included in the UKBiobank full release PCA analysis, which includes samples that were indicated as "het.missing.outliers" ("Indicates samples identified as outliers in heterozygosity and missing rates, which indicates poor-quality genotypes for these samples"), "excess.relatives" ("Indicates samples which have more than 10 putative third-degree relatives in the kinship table"), and whose "Submitted.Gender" were different from "Inferred.Gender". Applying these filters brought the sample size down to 407,219. We checked that the remaining sample contains only one out of any pair or group of related individuals with relatedness > 0.05.

We then selected samples indicated to be "in.white.British.ancestry.subset" ("Indicates samples who self-reported 'White British' and have very similar genetic ancestry based on a principal components analysis of the genotypes"), resulting in a sample size of 337,545. We then removed 337 samples indicated as having "putative.sex.chromosome.aneuploidy" ("Indicates samples identified as putatively carrying sex chromosome configurations that are not either XX or XY"). Finally, we removed 7 samples who have withdrawn their consent for use of their genetic data in analyses, arriving at our final set of 337,198 samples passing QC[1].

Of these samples, 37,041 were part of UK Biobank Lung Exome Variant Evaluation (UKBiLEVE), a study for chronic obstructive pulmonary disease (COPD). We retain all samples in UKBiLEVE, but as they are genotyped using a custom array optimized for coverage over regions implicated in lung health and disease[2], we consistently use genotyping array as a covariate in all our analyses.

For our analyses on different definitions of depression in UKBiobank, we further excluded 2,499 samples that indicated as having a history of substance abuse ("alcohol dependency": 1408, "opioid dependency": 1409 or "other substance abuse/dependency": 1410), manic or psychotic conditions (in the "Non-cancer illness code, self-reported" (data field 20002) section of the verbal interview, or who are classified under "Bipolar I Disorder" or "Bipolar II Disorder" in the "Bipolar and major depression status" (data field 20126) derived data field, giving a total of 334,699 samples for our study of the different definitions of MDD in UKBiobank.

**Minimal phenotyping definitions of depression in UKBiobank**

We define three minimal phenotyping-based definitions of depression in UKBiobank, each based on one or two questions instead of the full DSM-V diagnostic criteria, all of which are shown in Figure 1.

First, we identified 115,360 individuals who had sought help for "nerves, anxiety, tension or depression" from either their general practitioner (GPpsy, 113,260 out of 332,633, data field 2090) or a psychiatrist (Psypsy, 36,286 out of 333,412, data field 2100). 34,186 (29.6%) of those who sought help have seen both a GP and a psychiatrist. These make up most of the cases if the "broad depression" phenotype in Howard et al 2018[3], which we refer to and discuss in detail in the Supplemental Methods section "Comparison of definitions of depression in UKBiobank with those from previous GWAS".

Second, participants from 10 out of 21 assessment centers were asked whether they had ever experienced apathy and/or low mood (data fields 4631 and 4598), and how long they experienced it for (data fields 5375 and 4609). Having one or more of the two symptoms for 2 weeks or more is necessary, but not sufficient, for a DSM-V diagnosis of MDD. We applied this criterion together with help-seeking on the cases in GPpsy and Psypsy to identify 21,177 symptomatically defined cases (DepAll, data field 20126, returned by Smith et al 2013[4] which applied the same criteria), making up 70.7% of GPpsy and 78.0% of Psypsy who answered the questions on the two cardinal symptoms. Smith et al 2013[4] further divide these individuals into the following sub-categories: those with a single episode (DepSingle, 5,440 cases), and those with recurrent episodes of moderate depression (so defined because they were only examined by their primary care physician - DepRecurMod, 10,142) or severe depression (so defined because they were examined by psychiatrists - DepRecurSev, 5,595 cases). In this study, we do not consider the distinctions between these sub-categories.

Third, we identified 19,805 cases who self-reported having depression (SelfRepDep) during a verbal interview with a trained nurse (data field 20002, code 1286).

**Electronic health record-based definition of depression in UKBiobank**

We derive an electronic health record (EMR)-based definition of depression using the ICD10 codes in UKBiobank for both primary (data field 41202) and secondary diagnoses (data field 41204).

To qualify as a case of EMR-based definition of depression in UKBiobank, we require individuals to have a ICD10 code for any of the mood or affective disorders for either a primary or a secondary diagnosis, including: depressive episodes (F32), recurrent depressive disorder (F33), persistent mood (affective) disorders (F34), other mood (affective) disorders (F38), or unspecific mood (affective) disorders (F39). This is consistent with the definition of "ICD10-coded depression" in Howard et al 2018. Controls are those individuals who have not had either a primary or a secondary diagnosis of any of the above.

We then exclude from both cases and controls in this definition individuals that have the following primary or secondary diagnosis: delirium, not induced by alcohol and other psychoactive substances (F05), other mental disorders due to brain damage and dysfunction and to physical disease (F06), personality and behavioural disorders due to brain disease, damage and dysfunction (F07), unspecified organic or symptomatic mental disorder (F09), mental and behavioural disorders due to psychoactive substance use (F10-F19), schizophrenia, schizotypal and delusional disorders (F20-29), manic episodes (F30), and bipolar affective disorder (F31). Some of these other conditions are also excluded in "ICD10-coded depression" in Howard et al 2018, as we detail in the Supplemental Methods section "Comparison of definitions of depression in UKBiobank with those from previous GWAS".

**Strictly defined, CIDI-based definition of MDD in UKBiobank**

We derive a CIDI-based based *in silico* diagnosis of lifetime MDD using the CIDI questionnaire from the online mental health follow-up assaying DSM-V[5] symptom criteria for MDD (data category 138).

To qualify as a case of lifetime MDD according to DSM-V criteria, we first require individuals participating in UKBiobank to answer "Yes" to either the questions "Have you ever had a time in your life when you felt sad, blue, or depressed for two weeks or more in a row?" (data field 20446, DSM criterion A1 in Supplemental Table S2) or "Have you ever had a time in your life lasting two weeks or more when you lost interest in most things like hobbies, work, or activities that usually give you pleasure?" (data field 20441, DSM criterion A2 in Supplemental Table S2). Both are cardinal symptoms for DSM-V defined MDD. We then require them to have 3 or 4 among the criteria A3-A9 shown in Supplemental Table S2. We note that this questionnaire does not contain the DSM criterion A5, which requires "Psychomotor agitation or retardation nearly every day (observable by others, not merely subjective feelings of restlessness or being slowed down)", and hence assays only 8 out of the 9 symptoms for DSM-V MDD. Nonetheless, we require individuals to have a total of at least 5 symptoms, including at least one out of the two cardinal symptoms (A1 and A2) such that they have at least 5 out of 9 symptoms for DSM-V defined MDD, to fulfil the DSM-V A criterion: *Five (or more) of the following symptoms have been present during the same 2-week period and represent a change from previous functioning; at least one of the symptoms is either (1) depressed mood or (2) loss of interest or pleasure*.

To qualify as a case of current MDD according to DSM-V criteria, we first require individuals participating in UKBiobank to answer "Nearly every day" to cardinal symptoms "Feeling down, depressed, or hopeless" (data field 20510, DSM criterion A1 in Supplemental Table S2)" or "Little interest or pleasure in doing things" (data field 20514, DSM criterion A2 in Supplemental Table S2)", for the question "Over the last 2 weeks, how often have you been bothered by any of the following problems?". We then require them to have 3 or 4 among the criteria A3-A9 shown in Supplemental Table S2. This questionnaire, unlike the one for lifetime MDD, does contain the DSM criterion A5, which requires "Moving or speaking so slowly that other people could have noticed? Or the opposite - being so fidgety or restless that you have been moving around a lot more than usual", and hence assays all 9 symptoms for DSM-V MDD. We require individuals to have a total of at least 5 symptoms, including at least one out of the two cardinal symptoms (A1 and A2) such that they have at least 5 out of 9 symptoms for DSM-V defined MDD, to fulfil the DSM-V A criterion as described above.

Finally, we require those who fulfil the symptomatic criteria for both lifetime and current MDD to answer “Yes” to the question "Think about your roles at the time of this episode, including study / employment, childcare and housework, leisure pursuits. How much did these problems interfere with your life or activities?" (data field 20440), to fulfil the DSM-V B criterion: *The symptoms cause clinically significant distress or impairment in social, occupational, or other important areas of functioning*. Those who fulfil both DSM-V A and B criteria for either lifetime or current MDD are considered a “case” in our CIDI-based definition of MDD, LifetimeMDD. Supplemental Figure S1 shows the percentage endorsement of each symptom, as well as the total number of symptoms endorsed by cases of LifetimeMDD.

We excluded individuals who report a history of substance abuse, as well as manic or psychotic conditions so that our MDD cases all further fulfil criteria C, D and E of DSM-V:

*C. The (MDD) episode is not attributable to the physiological effects of a substance or to another medical condition.*

*D. The occurrence of the major depressive episode is not better explained by schizoaffective disorder, schizophrenia, schizophreniform disorder, delusional disorder, or other specified and unspecified schizophrenia spectrum and other psychotic disorders.*

*E. There has never been a manic episode or a hypomanic episode” respectively.*

Further, using the question on lifetime number of depressed episodes in the online mental health follow-up (data field 20442), we designated those cases of LIfetimeMDD who indicated they had two or more episodes as cases for recurrent CIDI-based MDD (MDDRecur) and those who didn’t as controls. We show the overlaps between definitions of MDD in UKBiobank in Supplemental Figure S2.

**The non-MDD definition of depression**

In order to understand the difference between the help-seeking definition of depression GPpsy and the cardinal symptoms-based definition of depression DepAll, we derived another phenotype GPNoDep, which contained individuals who were cases in GPpsy but not DepAll.

DepAll is defined based by answering a) “Yes” to “Ever unenthusiastic/disinterested for a whole week” (data field 4631) with “Longest period of unenthusiasm / disinterest” being greater than or equal to 2 weeks, or “Ever depressed for a whole week” (data field 4598) with “Longest period of depression” (data field 4609) being greater than or equals to 2 weeks, and b) having answered “Yes” to “Seen doctor (GP) for nerves, anxiety, tension or depression” (data field 2090) or “Seen a psychiatrist for nerves, anxiety, tension or depression”. These questions are answered by roughly half the participants in the UKBiobank (10 out of 22 collection centres, shown in Supplemental Table S4).

Controls in DepAll are those who have a) answered "No" to either symptoms, b) answered 0 or 1 week for durations of either symptoms, or c) answered "No" to seeing either a GP or Psychiatrist.

Cases in GPNoDep consist of those cases of GPpsy who are controls in DepAll, are those with a) or b). As such, they actively reported that they do *not* have one or other of the two cardinal symptoms of MDD (low mood or lack of interest). This forms 30% of cases in GPpsy who answered questions on the cardinal symptoms and their duration. As those questions on the cardinal symptoms and their duration are asked in individuals in 10 out of 22 collection centres, rather than those selected based on depression related traits, we can assume that collection of the same information in the other 12 collection centres will yield similar results. As such, we assume that those fulfilling case criteira of GPNoDep should constitute around 30% of the cases in GPpsy in UKBiobank.

We were only able to derive GPNoDep because data on cardinal symptoms of MDD was collected using the touchscreen questionnaire at baseline interview in UKBiobank, at the same time as information on whether one has sought medical help for depression was obtained. As such, GPNoDep is not meant to be a "gold standard" non-MDD phenotype, but one that is enabled by collection of information on both symptoms and self-reported seeking of medical attention at the same time point. It is likely that a proportion of cases in GPNoDep may meet criteria for lifetime MDD.

To further validate if cases in GPNoDep have MDD, we examined data collected at a different time point from the touchscreen interview at baseline, through verbal interviews (from which we derived SelfRepDep) or online mental health questionnaire (from which we derived LifetimeMDD).

Of the 58,125 individuals who had answered the questions necessary to have non-missing entries for GPNoDep, 43,613 (75.0%) answered the questions necessary to have a non-missing entry for self-reported depression SelfRepDep through verbal interview. Among this, 7139 are cases in GPNoDep, of which 610 (8.5%) are cases in SelfRepDep. If we take these data at face value, only 8.5% of the cases of GPNoDep who have gone to the GP for medical attention self-identify as having "depression" – a small proportion. Of the same 58,125 individuals in GPNoDep, 15,055 (25.9%) answered the questions in MHQ necessary to have a non-missing entry for self-reported DSM symptoms based LifetimeMDD. Among the 1,541 cases of GPNoDep that answered the MHQ, 307 (19.9%) are cases in LifetimeMDD. Again, if we take these data at face value, about a fifth of the cases in GPNoDep who answered the MHQ self-identify with symptoms that would constitute CIDI-based MDD. Both these findings are only enabled by the availability of self-report of depression or the symptom endorsement data from the online mental health questionnaire (MHQ).

Notably, the proportion of GPNoDep cases who answered the self-reported non-cancer illnesses during the verbal interview (16.4%) is significantly different from the proportion who did not answer (10.0%, Chisq test $P < 10^{-16}$, df = 1). Similarly, the proportion of GPNoDep cases who answered the MHQ (10.2%) is significantly different from the proportion who did not (16.5%, Chisq test $P < 10^{-16}$, df = 1). These numbers are shown in Supplemental Table S3. This shows that the percentages of GPNoDep cases who may qualify for SelfRepDep (8.5%) or LifetimeMDD (19.9%) are likely not an accurate estimate of the proportion of cases in GPNoDep that would self-report depression or qualify for CIDI-based MDD if all of them answered the relevant questions. We have little information to infer further due to the lack of self-report or symptom data on the rest of the cases and controls in GPNoDep. This points to the value of obtaining information on MDD symptoms in data collection.

**Comparison of definitions of depression in UKBiobank with those from previous GWAS**

In this section, we draw direct comparisons between definitions of depression included in this paper with that from previous studies that included UKBiobank data. We found that the minimal phenotyping definitions used in previous cohorts (such as "broad depression" in Howard et al 2018 and 23andMe in Hyde et al 2016, which contribute greatly to the sample size in Wray et al 2018) gave similar findings to help-seeking definitions of depression GPpsy in UKBiobank, while DSM-based cohorts like CONVERGE and PGC1-MDD show similar findings as CIDI-based LifetimeMDD in UKBiobank (see Supplemental Figure S6, cohorts are summarised in Supplemental Table S1). We explore the extent to which this finding reflects the use of the same definitions of MDD.

Cases in Wray et al 2018 are from a meta-analysis of multiple cohorts, including the first release data from UKBiobank. These cohorts are collected with vastly different approaches, and as such it is difficult to draw direct comparisons between the whole meta-analysis and definitions of depression included in this study. We consider individual cohorts from the PGC2-MDD included in Wray et al 2018 in our Supplemental Methods section "Cohorts in the MDD Working Group of the Psychiatric Genomics Consortium (PGC2-MDD)" and our Results section "Out-of-sample prediction of MDD".

Howard et al 2018 used only individuals from the UKBiobank and described in detail how each of the three definitions of depression that were included in their paper were derived, as reiterated below:

1. Broad depression, for which "Case and control status was determined by the touchscreen response to either of two questions: … UK Biobank field: 2090… or …UK Biobank field 2010. Caseness for broad depression was determined by answering "Yes" to either question at either the initial assessment visit, at any repeat assessment visit, or if there was a primary or secondary diagnosis of a depressive mood disorder from linked hospital admission records… The remaining respondents were classed as controls if they provided "No" responses to both questions during all assessments that they participated in."

In this phenotype, cases are the union of our definitions GPpsy, Psypsy, and ICD10Dep. Controls are all those who do not meet criteria for any of these definitions. There is no additional filter of either cases or controls.

2. ICD10-coded depression, for which "Participants were classified as cases if they had either an ICD-9/10 primary or secondary diagnosis for a depressive unipolar mood disorder … controls were participants who had linked hospital records, but who did not have any diagnosis of a mood disorder and were not probable MDD cases".

There is no mention of which ICD9 code was used to derive the ICD9 based cases or controls, so it is assumed that only the ICD10 codes were used in derivation of this phenotype. One additional filter was applied: controls for this definition that are cases for "probable depression" (as defined below) were removed.

3. Probable depression, for which cases and controls were defined "following the definitions from Smith et al… Cases for the probable MDD definition were supplemented by diagnoses of depressive mood disorder from linked hospital admission records (UK Biobank fields: 41202 and 41204) as per the broad depression phenotype."

This phenotype is the same as our definition DepAll, which is also derived from definitions from Smith et al 2013[4], with the addition of cases from ICD10Dep.

As such, all three definitions are combinations of definitions used in our study, but came with more inclusions and exclusions than we used. Here, we explain how these additional inclusions and exclusions are likely to impact their genetic architectures, in comparison to definitions we present in this paper.

First, Howard et al 2018 used information collected at two different time points (Touchscreen questionnaire and electronic medical record) for defining each of their depression phenotypes. Instead of assessing inter-rater liability between the two diagnoses and taking the intersection of cases obtained through two diagnoses to increase the confidence in their "caseness", they took the union to increase the total number of cases, which will result in either the same (at best) or higher heterogeneity between cases.

Second, Howard et al 2018 removed controls from "ICD10-coded depression" who are cases in "probable depression". As the questions that qualify one for a case or control of "probable depression" were only asked in half the collection centres in UKBiobank, Howard et al 2018 applied more stringent filtering of controls to one half of the collection centres in the UKBiobank (see our Result section "Definitions of depression in UKBiobank" and Supplemental Methods section "Control of population structure in heritability estimates" for our explanation of this). This creates different ascertainment biases between different collection centres, and creates confounding between population structure and case status.

Third, Howard et al 2018 excluded from both cases and controls those with ICD10 codes for some other mental health conditions: Bipolar (ICD codes F30, F31 or non-cancer illness code 1291), Multiple personality disorder (ICD code F44.8), Schizophrenia / psychosis (ICD codes F2*, or non-cancer illness code 1289). This removed samples from both cases and controls using the same criteria, and hence does not create artificial discontinuities in the liability distribution. However they have not excluded dementia (F00-F09) which may have affected the answering of the questionnaire, or substance abuse (F10-19) which is standard exclusion for MDD studies. In our definition ICD10Dep, we exclude those with codes F00-F31 (dementia, substance abuse, schizophrenia and psychosis, manic episodes and bipolar disorder) from both cases and controls.

Finally, Howard et al 2019 excluded different medication codes from cases and controls. This introduces artificial discontinuities in the liability distribution, making assessment of genetic architecture more difficult (as we explain in Supplemental section "Case enrichment and "cleaned" controls in GWAS").

**Prevalence of definitions of depression in UKBiobank**

In our analyses of definitions of depression in UKBiobank, we assume that the prevalence of each definition of depression in the population UKBiobank is represented by the prevalence in the dataset. This cannot be true, as not all participants in UKBiobank were asked questions from all categories. Questions for the Symptom-based definition DepAll were asked in only 10 out of 22 assessment centers in UKBiobank (Supplemental Table S4), and CIDI-based definitions LifetimeMDD and MDDrecur were derived from voluntary participation in the MHQ, which has been shown to have a genetic component genetically correlated with that of mental health conditions[6]. In addition, there may be self-report biases and uneven sampling of different demographics.

We carried out three analyses to ensure that the use of observed sample prevalences as population prevalence is unlikely to affect our conclusions.

First, using correction factors we derived using 2011 UK census data[7], we calculated a corrected prevalence of each definition of depression in the UKBiobank, and re-ran all our analyses of $h^2_{SNP}$ using that, with the results shown in Supplemental Tables S13. We find that discrepancies between sample and population prevalences for all definitions of depression we examine in UKBiobank are small, when corrected for regional populations, age and sex (Supplemental Tables S5-7).

Second, we obtained estimates assuming there is a "true" prevalence of 15% for MDD in European populations, as PGC1-MDD[8] had used, and show the results in Supplemental Tables S13. Both results are plotted out together with estimates using the observed sample prevalences derived directly from the number of cases and controls in each definition as population prevalence, (shown in Supplemental Figure S3). While there are some differences in the estimates, they do not qualitatively change our results: minimal phenotyping definitions of depression (GPPsy, Psypsy) show significantly lower $h^2_{SNP}$ than MDD defined with more stringent criteria (LifetimeMDD, MDDRecur).

Finally, we asked if there are is any combination of prevalences we can use to arrive at a conclusion that all definitions of depression in UKBiobank have the same liability scale h2SNP, comparable to the 0.20 found in

PGC1-MDD. We show in Supplemental Figure S7 that there is no such combination: liability scale h2SNP of minimal phenotyping definitions of depression cannot reach a liability scale h2SNP of 0.20 regardless of the prevalence we assume they have in the population.

**Control of population structure in SNP-heritability estimates**

Questions forming the criteria for different definitions of MDD are not all answered by the same individuals in UKBiobank, potentially leading to different levels of cryptic relatedness and population structure among individuals making up cases and controls in different definitions of MDD. For example, questions on cardinal symptoms of MDD in the touchscreen questionnaire, necessary for meeting the criteria for being a case in DepAll, are answered by only those who went to 10 out of 22 assessment centres (Supplemental Table S4).

To ensure we control for population structure adequately for all definitions of MDD in UKBiobank regardless assessment centre,, we assessed the per-chromosome heritabilities of all definitions of MDD estimated jointly and separately with BOLT-REML, using a total of 334,681 genotyped SNPs with MAF > 5% and HWE P-value > $10^{-6}$ across all chromosomes as model SNPs. Difference between the slopes between per chromosome $h^2_{SNP}$ and length of chromosome (approximated with number of SNPs used as model SNPs in each chromosome) in the two models (joint and separate) reveals population structure that induces long-range LD between chromosomes. This is because in presence of population structure $h^2_{SNP}$ attributable to SNPs on one chromosome "leaks" into the estimates of a different chromosome, due to long-range LD, when $h^2_{SNP}$ are estimated separately.

We find that the difference between the two slopes are minimal when using PCs calculated using all White-British samples in UKBiobank, which we used as covariates in $h^2_{SNP}$ estimates and GWAS on the definitions of MDD in UKBiobank (Methods). We show the results in Supplemental Figure S3; we therefore use all-samples PCs in all the $h^2_{SNP}$ estimates and GWAS we report, except for stratified analyses where numbers of samples in each stratum are substantially smaller than the whole UKBiobank White-British cohort where we used definition and stratum specific PCs.

As recommended by the UKBiobank[1], we removed the major histocompatibility complex (MHC) region as well as known structural variants (SVs)[9] before selecting SNPs for the computation of PCs (Methods) which we use for all association and $h^2_{SNP}$ analyses. As a result, there is little control over population structure over these regions in both association testing and $h^2_{SNP}$ estimation. This may lead to false positive associations as well as inflated $h^2_{SNP}$ estimates.

We therefore indicated whether each significant association is in MHC and SV regions in Supplemental Table S10, S11, S15 and S16, and use hollow points to represent SNPs in Manhattan plots (Supplemental Figure S5 and S10) - the validity of these associations can be followed up with sequencing of regions involved, and does not fall in the scope of this paper.

To check if lack of population structure control over SVs and MHC regions affect $h^2_{SNP}$ estimates, we estimate $h^2_{SNP}$ using LDSC using all SNPs (LDSC-AllSNPs), excluding SNPs in the MHC region on chromosome 6:25-35MB (LDSC-noMHC), and excluding SNPs in the both MHC region and SVs [9] (LDSC-noMHCSVs). In Supplemental Figure S3 we show that while $h^2_{SNP}$ decreases from LDSC-AllSNPs estimates when we remove MHC or/and SVs, the decreases are not significant in any of the definitions of MDD in UKBiobank, and the trend between definitions of MDD remain the same. As such, we conclude that excluding MHC and SVs in the calculation of PCs is unlikely to cause significant biases in estimation of heritabilities of particular or all of the definitions of MDD.

**Measure of lifetime trauma**

A binary measure of lifetime trauma was derived from self-reported experience of traumatic events from the online mental health follow up questionnaire (data category 145), in order to identify individuals exposed to severe environmental adversities. The online mental health questionnaire for traumatic events consist of 16 questions on childhood, adulthood and lifetime trauma. We have scored individuals as having experienced each traumatic event as "1" and those who have not experienced each traumatic event as "0", as shown in Supplemental Table S12. Of the 16 questions, we have included 12 for derivation of an aggregate "lifetime trauma" measure, the remaining 4 capture the same traumatic events the other questions we have included do, and are hence redundant.

We asked how much each traumatic experience contributes to the risk of developing LifetimeMDD, the CIDI-based definition of MDD in UKBiobank, by jointly modeling their contribution in a logistic regression, along with age, sex, social deprivation (using the Townsend deprivation index), years of education, experience of any traumatic life event in the past 2 years (data field 6145), neuroticism, and total number of traumatic experiences. The odds ratios (ORs) of each traumatic experience on LifetimeMDD is shown in Supplemental Table S12.

Since traumatic events vary in severity, a lifetime trauma score was constructed by weighting each item by their effect size on LifetimeMDD, and summing across all 12 items. As not all questions about traumatic experience are answered by all individuals who took part in the online mental health follow-up, we removed individuals who did not answer more than 3 questions on traumatic experience. We obtained lifetime trauma score for 109,699 individuals, with mean score 1.90 (standard deviation = 2.11). We categorize 33,619 individuals with scores above 2.5 as having experienced significant lifetime trauma (who report a mean of 3.17 traumatic experiences) and those with scores below 2.5 as not having experienced significant lifetime trauma (who report a mean of 0.66 traumatic experiences).

We note the following caveats in our use of self-reported answers to questionnaires to infer level of lifetime trauma. First, in weighting traumatic events based on their effects on LifetimeMDD in creating the measure of lifetime trauma, one can potentially incur biases in genetic associations to LifetimeMDD at genetics variants that contribute to both lifetime trauma and LifetimeMDD when one stratifies individuals in LifetimeMDD by lifetime trauma. However, this bias will likely be small, as the weighted lifetime trauma score is highly correlated with the number of traumatic experiences an individual self-reports ($r2=0.92$, $p < 10^{-16}$), and this self-report is assumed to be independent of LifetimeMDD (see second caveat). Moreover, even if present, this bias will not extend to other definitions of MDD in UKBiobank acquired through independent questionnaires.

Second, as both questionnaires on traumatic events and CIDI-based MDD symptoms (used for specifying LifetimeMDD) are on the same online mental health follow up assessment, how one answers one questionnaire may affect how one answers the other, potentially incurring errors in retrospective recall. This is an unavoidable problem in UKBiobank given the structure of the questionnaire, and as such here we assume that the errors incurred are negligible. In addition, as the online mental health follow-up is conducted a year or two after one takes part in the initial assessment through the touchscreen questionnaire, there should be little to no effects of retrospective recall errors on all other definitions of MDD in UKBiobank, which we derive from answers to questions in the touchscreen questionnaire.

**Epidemiological analysis of risk factors for MDD**

We assess the effects of known risk factors for MDD on the different definitions of MDD in UKBiobank; for each binary or quantitative risk factor, we estimatd its odds ratio (OR) for each definition of MDD using logistic regression, correcting for UKBiobank collection centers (data field 54, as factors), as well as years of education (data field 845, quantitative), age (data field 21022, quantitative) and sex (data field 31, binary) unless one of three factors is being tested. The binary risk factors we tested are:

1. Age: data field 21022; dividing individuals into those < 60 years old and those >= 60
2. Sex: data field 31; dividing individuals into males and females
3. lifetime trauma: data category 145 for "Traumatic events reported within the on-line mental health questionnaire"; we calculated a weighted measure of traumatic events, dividing individuals into those who have weighted measure = 1 and those with weighted measure = 0.

The quantitative risk factors we tested are:

1. neuroticism: data field 20127 for "12 neurotic behaviour domains as reported from fields 1920, 1930, 1940, 1950, 1960, 1970, 1980, 1990, 2000, 2010, 2020 and 2030 from the touchscreen questionnaire at baseline"
2. social deprivation: data field 189 for townsend deprivation index (TDI) calculated for per individual just before baseline assessment)
3. years of education: data field 845; for age one stopped continuous full-time education, from which we infer years of continuous education received by each individual assuming starting age of 6.

For quantitative risk factors we report ORs for per SD increase in the measure. The UK Biobank cohort contains more women than men (the female to male ratio is 1.16 to 1), with more younger women than men (there are 6% more women than men younger than 60, compare to 2% older than 60) and is more educated among the young than the old (those with more than 10 years of education form 42.5% of those younger than 60, compare to 28.4% among those older than 60). Thus, in each analysis we included age, sex and number of years of education as covariates unless they are the risk factors tested (together with collection center).

**Potential confounding from self-ascertainment bias**

A recent preprint has shown that voluntary participation in the MHQ has a genetic component genetically correlated with that of mental health conditions[6]. This raises the concern that difference between definitions of depression in their genetic architecture are in fact driven by genetic differences between those who answered this questionnaire and those who did not. In fact, this is a more general problem that affects all phenotypes in the UKBiobank, all of which are dependent on voluntary self-reporting. Using the derivation of CIDI-based definitions of MDD LifetimeMDD from the MHQ as an example, we explain how we may understand self-ascertainment bias.

First, if the self-ascertainment bias to participate in the MHQ is directly driven by MDD (the underlying condition rather than definitions of MDD from the self-reports), where those with MDD are more likely to participate, then the prevalence of cases of MDD will be higher among those who have taken the MHQ. This is consistent with what we observe with our CIDI-based definitions of MDD in UKBiobank LifetimeMDD, both of which have much higher case prevalence than expected (0.243 and 0.173 respectively). In this scenario, the ascertainment bias for MDD from using data from MHQ is akin to the ascertainment in any GWAS where cases are oversampled (the vast majority of disease GWAS to date).

This does not violate assumptions used in estimation laid out in section "Case enrichment and use of "clean" controls in GWAS" – in particular it does not violate assumption 4 "The probability for a case or a control to be selected for the study, from all cases and all controls in the population, depends only on their phenotype and not any other factors". As explained in the section "Comparison of methods of heritability and genetic correlation estimation", PCGCs can account for the over-sampling of cases better than other methods. For this reason we use it for estimation of $h^2_{SNP}$ of all definitions of depression, as well as to estimate rG between them.

Second, the self-ascertainment bias to participate in the MHQ may not be driven by MDD itself, but by conditions genetically correlated with MDD, either alone or in combination with MDD. In this scenario, those with MDD are more likely to participate, and those with conditions correlated with MDD (but who are not cases of MDD) are also more likely to participate. What biases result from this scenario depends on whether participation in the MHQ is independent of one's ability to answer questioning relating to MDD symptoms accurately.

If we assume participation in the MHQ *does not* affect one's ability to answer questions relating to MDD symptoms, then our bias will lie in the controls – these controls contain more individuals with conditionally genetically correlated with MDD (A). If we believe participation in the MHQ *does* affect one's ability to assess whether one has MDD symptoms, then a bias may be present in both cases and controls. This bias may be consistent between cases and controls (B), or different between cases and controls (C). B does not violate any assumptions outlined in "Case enrichment and use of "clean" controls in GWAS", and hence does not cause problems in $h^2_{SNP}$ estimation – this scenario can be seen as $h^2_{SNP}$ of LifetimeMDD is estimated in a different population as that of other definitions of depression in UKBiobank. A and C are problematic in the same way – they violate assumption 4 "The probability for a case or a control to be selected for the study, from all cases and all controls in the population, depends only on their phenotype and not any other factors", and introduce a correlation between defined depression phenotype and the underlying condition that causes the self-ascertainment bias to participate in the MHQ. If this underlying cause has a genetic component, then this genetic component can appear to have effects on LifetimeMDD too. No methods for assessing $h^2_{SNP}$ to date can account for this adequately.

However, we can ask if genetic effects we observe in LifetimeMDD are driven by this underlying cause using the Mendelian Randomization (MR) framework. If MR shows a significant correlation between genetic effects on MHQ participation and that on LifetimeMDD, then we can conclude this is a significant problem, and we cannot trust results from LifetimeMDD. As such, we obtained 25 independent variants (LD $r2 < 1$) from Adams et al 2019, of which 16 are SNPs with INFO scores above 0.9 in the UKBiobank that we include in GWAS on our definitions of depression (Supplemental Table S8).

We performed MR using the MendelianRandomization R package[10], with the following estimators: simple median, weighted median, inverse-variance weighted (IVW)[11] and MR-Egger[12], as shown in Supplemental Table S9. None of the estimators shows significant causal effect of MHQ on any definition of depression (threshold P-value = 0.05/36 = 0.0016). As there are two outlier SNPs with high effects on answering of MHQ (exposure, as shown on Supplemental Figure S4) that may bias our results, we performed the analysis again removing the two SNPs, and still found no significant causal effect of answering the MHQ on any definition of depression (Supplemental Figure S4, Supplemental Table S9). Further, without a significant causal effect from any of the estimators, the MR-Egger intercept cannot be used to infer pleiotropy[13]. We find no support for the view that the genetic factors influencing whether one answers the MHQ has an impact on our findings on the definitions of depression.

**Case enrichment and use of "clean" controls in GWAS**

Many GWAS studies of MDD and of other conditions apply additional filters to the diagnostic criteria to improve power of association. One such additional filter is to remove controls which nearly qualify as cases of the disease by the same set of diagnostic criteria, or qualify as cases of the disease of interest by a different set of diagnostic criteria. For example, in Howard et al 2018[3], controls in "ICD10-coded depression" who qualified as cases of "Probable depression" were excluded from the GWAS (we explore this further in the next Supplemental Methods section). We refer to this approach as using "clean" controls.

Use of "clean" controls does increase power in GWAS. In addition to performing GWAS using all cases and controls for each definition of depression in UKBiobank we define in this paper, we obtained "clean" controls, which are not cases in any other definition of depression, and performed GWAS using these "clean" controls. We obtain more GWAS hits than using all cases and controls (Supplemental Figure S3, Supplemental Table S10-11). However, we do not use "clean" controls for any analysis of genetic architecture and comparison between definitions, for two reasons.

First, our paper focuses on comparing different strategies of phenotyping, and therefore only considered phenotypes that can be obtained with information taken at one time point. Given we are effectively "simulating"

GWAS studies that only collect 1 piece of information at a time, we should not be able to arrive at "clean" controls.

Second, using "clean" controls introduce artificial discontinuities in the liability distribution that are difficult to account for[14]. Under the liability threshold model, liability to a disease in a population is contributed by both genetic and environmental factors. The binary case/control status is assigned by defining a liability threshold corresponding to the observed disease prevalence, where everyone above the threshold would be classified as cases, and everyone below it would be controls. This forms the basis for the following assumptions used in estimating liability scale SNP-heritability ($h^2_{SNP}$) with the conversion from observed scale proposed by Dempster and Lerner[15] and appropriated for $h^2_{SNP}$ estimation using REML by Lee et al [16], which is later adopted for BOLT-REML[17], summary statistics based method LDSC[18], and exact methods for binary traits PCGC[19] and PCGCs[20].

1. The liability is normally distributed. A corollary of this is that unmeasured environmental factors induce a normally-distributed effect on the phenotype that is independent from the normally distributed effect exerted by the genotypic factors.
2. Each causal SNP exerts a linear effect on the phenotype
3. Each covariate exerts a linear effect on the phenotype
4. The probability for a case or a control to be selected for the study, from all cases and all controls in the population, depends **only** on their phenotype and not any other factors

If we had used a "clean" set of controls (those who would come up negative on criteria for all definitions of depression) for GWAS on each definition of depression, we would be violating assumption 4 and introducing an artificial discontinuity in the liability distribution. This bias our estimates of $h^2_{SNP}$ and all other measures of genetic architecture based on it.

**Comparison of methods of SNP-heritability and genetic correlation estimation**

Many methods have been developed to estimate narrow sense SNP heritability ($h^2_{SNP}$) from either whole-genome SNP data[17,19,21,22] or summary statistics of association tests [18,20,23] across the whole genome, and the sensitivity of $h^2_{SNP}$ estimates to assumptions in different models employed by the different methods have been discussed and reviewed extensively in the past few years[24,25].

For all the $h^2_{SNP}$ estimates in our analysis we use the phenotype-correlation-genotype-correlation (PCGC) approach, an adaptation of Haseman-Elston regression. We prefer this method because it produces unbiased estimates of $h^2_{SNP}$ and genetic correlation (rG) for case-control phenotypes[19,20] in the presence of covariates of large effect, such as sex and age. As mentioned in the Results section as well as the Supplemental Methods section "Prevalence of definitions of MDD in UKBiobank", cases and controls for some or all of the definitions of depression we use may not represent the full liability distribution of the population. Instead they may represent a potentially skewed or discontinuous proportion of it. For example, LifetimeMDD is derived from the MHQ, which is voluntarily answered by a non-random proportion of individuals in UKBiobank, and the answering of the MHQ has been shown to be genetically correlated with mental health traits[6]. When this is the case, the second assumption we outlined in the Supplemental methods section "Case enrichment and use of "clean" controls in GWAS" is violated – the effects of the unmeasured environmental factors on the phenotype of interest (definitions of depression) are potentially no longer independent of the effects of the genetic factors. This leads to downward bias in $h^2_{SNP}$ estimates from LDSC and REML-based methods, even when the relevant covariates (which we assumed to be PCs in this case, accounting for population structure in each definition of depression) are used in GWAS[19,20]. Such bias will propagate to other measures of genetic architecture based on $h^2_{SNP}$, such as rG. As Weissbrod et al 2018[20] explained, this is a problem that can be circumvented by "regressing the omitted covariates out of the genotypes and correcting the individual-level affection cutoffs prior to parameter estimation or to computing summary statistics", as is performed in PCGCs.

As such, we use the PCGCs[19,20] framework, generating summary statistics for definitions of MDD in UKBiobank in our study which can be reused for estimation of genetic correlation with other phenotypes from other cohorts processed the same way.

We note, however, that previously published estimates of $h^2_{SNP}$ of MDD[3,8,26-28], are mostly generated using LDSC[18] if they were estimated from GWAS summary statistics, or either GCTA[22], LDAK[21], or BOLT-REML[17] if they were estimated from individual level genotypes. All estimate $h^2_{SNP}$ of case-control traits on the "observed scale" under a quantitative trait framework, and apply a correction factor to results to convert them to a "liability scale" to account for the ascertainment of cases in case-control studies. The same applies when estimating rG.

In Supplemental Table S13, we show the $h^2_{SNP}$ estimates of each definition of MDD in UKBiobank using PCGCs[19,20]., LDSC[18], and BOLT-REML[17]. For both PCGCs and LDSC, we use 8,968,716 imputed SNPs with MAF > 5% in our analysis (Methods), while for BOLT-REML we use 334,681 genotyped SNPs with MAF > 5% and HWE P-value > $10^{-6}$ as model SNPs. For results from LDSC and BOLT-REML, we use LD scores calculated on all imputed SNPs in UKBiobank, using LDSC as reference, and convert the $h^2_{SNP}$ estimates on the observed scale to liability scale specifying the prevalence of each definition of MDD in UKBiobank as shown in Supplemental Table S8.

Results from all three methods show the same trend - the CIDI-based definitions LifetimeMDD and MDDRecur show the highest $h^2_{SNP}$ estimates while the help-seeking based definitions GPpsy and Psypsy, as well as their no-MDD components GPNoDep and PsyNoDep, show the lowest $h^2_{SNP}$ estimates. While LDSC and BOLT-REML estimates are highly similar to each other, PCGCs estimates are higher than both of them, consistent with expectations of downward bias in $h^2_{SNP}$ estimation of case-control traits using LDSC and REML due to handling of covariates.

For comparison of $h^2_{SNP}$ estimates from our study with previously published estimates using LDSC, we show in Supplemental Figure S6 that the CIDI-based definitions LifetimeMDD and MDDRecur have heritabilities closer to LDSC $h^2_{SNP}$ estimates of MDD in CONVERGE[26] and PGC1-MDD[8], while help-seeking based definitions GPpsy and Psypsy have estimates closer to that in similarly defined, minimal phenotyping based MDD in 23andMe[27].

For PCGCs, we show estimates of genetic variance and its standard error for every definition of depression in UKBiobank. "Genetic variance" is variance attributable to genetics over total phenotypic variance (referred to as "marginal heritability" in Weissbrod et al 2018[20]), and is the measure comparable measure to $h^2_{SNP}$ from BOLT-REML and LDSC (Supplemental Table S13). For completeness, we also show estimates of "conditional heritability", which is genetic variance divided by the total liability variance, and which also includes variance introduced by fixed effects (in this case, genotype PCs and genotyping array, Supplemental Table S13).

We also compared the estimation of rG between definitions of MDD using PCGCs and LDSC. In Supplemental Figure S6 we show the genetic correlation between LifetimeMDD, the CIDI-based definition, with all other definitions of MDD in UKBiobank estimated using LDSC (Figure 3b shows the PCGCs estimates). LDSC estimates show a similar trent to those from PCGCs, though both point estimates are higher across the board and standard errors are larger than from PCGCs, making most estimates not significantly different from 1. This is consistent with the expectation that LDSC may over-estimate genetic correlations due to mishandling of covariates and overlap of samples[20]. Overestimates of rG between definitions of MDD (not significantly different from 1) may send the misleading message that the genetic architectures in different definitions of MDD, even those on opposite ends of the spectrum in terms of case criteria, are not significantly different from each other. This obscures important true genetic differences between different definitions of MDD.

**Stratification of definitions of depression by environmental risk factors**

We stratified samples from definitions of MDD in UKBiobank by environmental risk factors in the following ways. We estimate $h^2_{SNP}$ in each strata of samples using PCGCs (Methods). Results are shown in Supplemental Table S14.

1. by age: data field 21022; dividing individuals into those < 60 years old and those >= 60
2. by sex: data field 31; dividing individuals into males and females
3. by lifetime trauma: data category 145 for "Traumatic events reported within the on-line mental health questionnaire"; we calculated a weighted measure of traumatic events, dividing individuals into those who have weighted measure >= 2.5 and those with weighted measure < 2.5
4. by neuroticism: data field 20127 for neuroticism score; dividing individuals into those who score >= 5 and those who score < 5
5. by social deprivation: data field 189 for Townsend deprivation index (TDI); dividing individuals into those with TDI <0 and those with TDI >=0
6. by years of education: data field 845; dividing individuals into those with years of education >= 11 and < 11).

Using the $h^2_{SNP}$ estimates and standard errors from PCGCs, we performed t-tests on the $h^2_{SNP}$ from the strata in each stratification, and ask whether they are significantly different from 0 at $p < 0.05$, using a two-tailed test and correcting for 49 stratifications. None of the difference between strata, from any definition of MDD and stratification, is significant, though the method is underpowered as compared to other methods that require the use of individual level genotypes[29].

**Simulations of effects of misdiagnosis and misclassification on SNP-heritability estimates**

In this section we explore whether minimal phenotyping definitions of MDD represent milder forms of the disease (those with lower liability) that do not qualify for CIDI-based definition of MDD, or whether they contain misdiagnosis of those without the disease as cases.

We perform simulations to show the effects of misdiagnosis and misclassification on $h^2_{SNP}$ estimates, and in turn, whether they may be the cause of lower $h^2_{SNP}$ estimates in the minimal phenotyping definitions of MDD. We adopt the theoretical framework of the liability threshold model, where every individual has a normally distributed liability value for a trait such that case subjects of the trait are individuals whose liability exceeds a given cutoff[15]. The cutoff for the liability was determined as the 1-$K_i$ percentile of the simulated liabilities, where $K_i$ for the i-th simulated trait is set as the prevalence of cases. To simulate a biobank sample (which is assumed to be representative of the population), we do not ascertain for cases.

Using array genotypes at 344,184 SNPs (LD < 0.5, MAF > 5%, HWE P-value > 10e-6) in 25,000 random individuals from UKBiobank who are White-British (Methods), we used LDAK[21,30] to simulate pairs of traits $y_{i,1}$ and $y_{i,2}$ with genetic correlation $rG_i$, where for each $i$ in $i \in \{1..10\}$: 5000 causal SNPs are picked uniformly at random to contribute a total $h^2_{SNP}$ for $y_{i,1}$ and $y_{i,2}$ of $h^2_{i,1} = h^2_{i,2} = h^2_i$ using the model $Y = \sum \beta_j X_j + e$, where $\beta_j$ is the effect size of $X_j$, the $j$th causal SNP, and $e$ is Gaussian-distributed noise. Effect size at each SNP $X_j$ is sampled under the Model $\beta_j \sim \mathrm{N}(0, [f_j(1 - f_j)] - 1)$, where $f_j$ is the MAF of $X_j$.

The LDAK command we use is as follows:

*ldak5.linux --make-phenos traitpair$i.h2$h2i.rg$rGi --bfile $bfile --ignore-weights YES --power -1 --num-causals 5000 --num-phenos 2 --her $h2i --bivar $rGi*
*Where* $i \in \{1..10\}$, $h_i^2 \in \{0.2, 0.4, 0.6, 0.8\}$, $rG_i \in \{0, 0.2, 0.4, 0.6, 0.8, 0.95\}$

In the first set of simulations, we show that including individuals with lower liabilities to a disease (trait) as cases (increasing prevalence of cases through lowering the liability cutoff) would not have an effect on liability scale $h^2_{SNP}$ estimates of the trait. We estimate the liability scale $h^2_{SNP}$ of each trait using the --pcgc option in LDAK appropriate for binary traits, accounting for the prevalence of cases $K_i$ used. As we both do not ascertain for cases, and do not simulate covariates with effects on the traits, $h^2_{SNP}$ estimates from the simulations do not suffer from downward estimates in REML presented previously for case-control traits[20] when converted to the liability scale[15,19]. As such, using both --reml with --prevalence and --pcgc options give the same liability scale $h^2_{SNP}$ estimates on the simulated traits. Supplemental Figure S8 shows the $h^2_{SNP}$ estimates of traits with heritabilities $h_i^2 \in \{0.2, 0.4, 0.6, 0.8\}$ for $i \in \{1..10\}$, where we shift the prevalence $K_i$ from 0.1 to 0.5, and recover the simulated $h^2_{SNP}$ exactly regardless of $K_i$. While this is well-known, it refutes the misconception that minimal phenotyping definitions of MDD have lower $h^2_{SNP}$ than CIDI-based definitions because they contain individuals with lower liabilities to MDD.

In the second set of simulations, we show that misdiagnosis of controls as cases lowers $h^2_{SNP}$. At a constant liability threshold corresponding to $K_i = 0.2$, if we identify all individuals above liability threshold correctly as cases, but identify random controls as cases such that the percentage of misdiagnosis cases ranges from 0% to 50% of all cases (thereby increasing apparent case prevalence), then liability scale $h^2_{SNP}$ (corrected for the apparent prevalence) decrease as a result (Supplemental Figure S8). This decrease is consistent with the high prevalence and low $h^2_{SNP}$ we observe in minimal phenotyping based MDD in UKBiobank and 23andMe[27] (Figure 3a, Supplemental Figure S6). We note that our simulations do not misidentify true cases as controls (in other words, we have a sensitivity of 100% for true cases, while having a false discovery rate of cases ranging from 0% to 50%); in realistic settings, it is possible sensitivity to true cases will decrease as collection criteria becomes less stringent. If we lower the sensitivity, the $h^2_{SNP}$ estimates are likely to decrease further.

However, the above scenario must not be the only explanation for the lower $h^2_{SNP}$ in minimal phenotyping based MDD. Genetic correlation between definitions of MDD should not differ if all genetic factors are shared and effects were only lower in some definitions due to noise (Figure 3b). Hence, we hypothesize minimal phenotyping definitions of MDD may in fact contain cases of other conditions that are misclassified as MDD. We simulate misclassification of cases from a genetically correlated disease to find out if misclassification may result in a decrease in $h^2_{SNP}$ consistent with the lowered $h^2_{SNP}$ estimate in minimal phenotyping definitions of MDD.

We first correctly identify all individuals above liability threshold for prevalence $K_{i,1} = 0.2$ in $y_{i,1}$ ($h_{i,1}^2 = 0.4$) as cases of $y_{i,1}$. Then, we misclassify 10% to 50% of cases of genetically correlated $y_{i,2}$ ($h_{i,2}^2 = 0.4$, prevalence $K_{i,2} = 0.2$) as cases of $y_{i,1}$, hence increasing the apparent case prevalence of $y_{i,1}$. The two traits have genetic correlation $rG_i \in \{0,0.2,0.4,0.6,0.8,0.95\}$. As shown in Supplemental Figure S8, we find that increasing the percentage of misidentification decreases liability scale $h^2_{SNP}$ of $y_{i,1}$ (corrected for the apparent prevalence) if $rG_i$ is small, but can inflate liability scale $h^2_{SNP}$ if $rG_i$ is large (>=0.6). This inflation decreases with increase in $h^2_{SNP}$ of both traits (Supplemental Figure S9). This analysis shows that as long as two traits are completely heritable, and there are environmental contributions, misclassifying cases of one as cases of the other would lead to erroneous estimates of $h^2_{SNP}$, even at very high genetic correlations.

As genetic correlation between definitions of MDD and other psychiatric conditions are low (maximum genetic correlation with neuroticism at a mean of 0.67 among all definitions of MDD, Supplemental Table S17, Figure 4a), and genetic correlation between CIDI-based definition of MDD (LifetimeMDD) and those without MDD (GPNoDep, making up 30% cases in minimal phenotyping definition of MDD GPpsy) is 0.58 (se=0.078), most of the potential misclassification could have come from conditions with genetic correlation with MDD lower than 0.6 and led to downward bias of $h^2_{SNP}$ observed in minimal definitions of MDD in UKBiobank.

In summary, our simulations are consistent with a model where the lower $h^2_{SNP}$ in minimal phenotyping based MDD is not due to lowering liability threshold for case definition in MDD, but a consequence of potentially both misdiagnosis of those who do not have MDD as cases, and misclassification of those with other conditions as cases (and the misidentification of true cases as controls, not simulated).

**LDSC-SEG analysis of tissue-specific enrichment of $h^2_{SNP}$**

The LDSC-SEG analysis tests whether the $h^2_{SNP}$ of a disease is enriched in regions surrounding genes with the highest specific expression in a given tissue[31]. In a meta-analyses of several depression studies Wray et al 2018 demonstrated significant enrichment of $h^2_{SNP}$ in regions surrounding genes whose expression is highest in CNS tissues, including the frontal cortex, anterior cingulate cortex, putamen, nucleus accumbens, hippocampus and substantia nigra[32].

We tested the extent to which the different definitions of MDD in UK Biobank showed similar enrichment.. We performed the same LDSC-SEG analysis on all definitions of depression in UKBiobank, all psychiatric conditions included in the PGC Cross Disorder Working Group, including MDD (PGC1-MDD), schizophrenia (SCZ), bipolar disorder (BIP), attention deficit hyperactive disorder (ADHD), and autism (AUT), as listed in Supplemental Table S1, and both smoking and neuroticism in UKBiobank (Supplemental Figure S10, Supplemental Tables S15-16). For comparison, we also included an analysis of the minimal phenotyping definition of depression 23andMe[27] (Supplemental Table S1).

We found that the strictly defined definitions of MDD in UKBiobank (LifetimeMDD), as well as MDD defined by structured interview in PGC1-MDD [8], do not show CNS enrichment, while minimal phenotyping definitions of depression such as help-seeking based GPpsy and self-report based 23andMe do. This is counter-intuitive, given a CNS enrichment is expected for MDD. We therefore investigated the potential reasons for differences between tissue-specific enrichments of $h^2_{SNP}$ between strictly defined MDD (LifetimeMDD and PGC1-MDD) and minimally defined depression (GPpsy and 23andMe).

Differences in sample size cannot explain the observation. LifetimeMDD consists of 16,301 cases and 50870 controls and shows no enrichment. To examine the effect of sample size we down-sampled all definitions of depression in UKBiobank to N = 50,000, each with a case prevalence of 0.15 (N cases = 7,500) to ensure all definitions have the same statistical power, and performed the same analysis. As shown in Supplemental Figure S11, for most definitions of depression no tissue-specific enrichment is seen, including GPpsy. However, enrichment in CNS tissue is observed for GPNoDep, consisting of 8,632 cases and 49,493 controls. Forming 30% of the cases in GPpsy, this suggests that GPNoDep in fact drives the CNS enrichments seen in GPpsy.

We asked whether CNS enrichment, or any tissue-specific enrichment, can be used to validate a particular phenotype. We found that other psychiatric conditions including SCZ, BIP, neuroticism and smoking all show CNS enrichment of $h^2_{SNP}$. It is clear that CNS enrichment is not specific to MDD.

We also observed enrichment in the 29 cohorts from MDD working group of the PGC (PGC29 [32,33]) (16,823 cases and 25,632 controls) as shown in Supplemental Figure S11. Could it be the case that the failure to detect signal in UKBiobank LifetimeMDD is due to the greater noise in self-reported questionnaires, compared to the interviewer diagnosis used for the PGC 29 cohorts? If that were true, then a meta-analysis of LifetimeMDD and PGC29 should show enrichment. As shown in Supplemental Figure S11, this is not the case.

Differences between LifetimeMDD and PGC29 may reflect genetic architecture differences between MDD derived from MHQ self-reporting and structured interviews. These differences may result in different tissue-specific enrichment of $h^2_{SNP}$. On one hand, MHQ based self-reporting may capture sub-clinical MDD; on the other, there may be differences in how strictly CIDI-based diagnostic criteria were adhered to in PGC29. As shown in Supplemental Table S20, cohorts in PGC29 have varying percentages of cases fulfilling DSM diagnostic criteria. There is also heterogeneity in ascertainment strategies and genetic outcome among cohorts in PGC29[33]. This heterogeneity is reflected in differences in individual cohort $h^2_{SNP}$[33], as well as the lower $h^2_{SNP}$ of PGC29 compared with PGC1-MDD (Supplemental Figure S6). LifetimeMDD has a genetic architecture more similar to PGC29 cohorts with a greater percentage of cases fulfilling DSM criteria, as demonstrated in the out-of-sample prediction analysis (Figure 7, following section in Supplemental Methods).

GPNoDep is the no-MDD definition that most likely contained mis-diagnosed or misclassified cases, consistent with its low $h^2_{SNP}$ and the deflation of $h^2_{SNP}$ by both mis-diagnosis and misclassification (Figure 3, Supplemental Figures S7-9, Supplemental Methods). Sources of such mis-diagnosis and misclassification in GPNoDep, and by extension GPpsy, could potentially be neuroticism and smoking, both of which demonstrate CNS enrichment. Further, GPNoDep, neuroticism and smoking replicate GWAS hits of GPpsy, and mirror effect size distributions of GWAS hits of GPpsy (Figure 6), consistent with this hypothesis. 23andMe (N = 307,354, 75,607 cases and 231,747 controls), like GPpsy, has a CNS enrichment that likely contributes substantially to the CNS enrichment seen in Wray et al 2018[32], due to its large sample size compared to the rest of the cohorts. Its GWAS hits are also replicated by neuroticism and smoking (Supplemental Figure S12).

In summary, whether a disease phenotype shows an enrichment of $h^2_{SNP}$ in a particular expected tissue depends on a variety of factors including sample size, phenotyping strategy, and heterogeneity, in addition to whether the expected tissue truly harbor enrichments of $h^2_{SNP}$ for the disease.

**Out-of-sample prediction of MDD in PGC cohorts**

To assess the relative power of the different definitions in UK Biobank to index MDD, we carried out an out-of-sample prediction using data from the MDD working group of Psychiatric Genomics Consortium. We obtained access to individual level genotype and phenotype data from 25 out of the PGC29. The 4 cohorts we did not apply to permission to are pharmaceutical company cohorts. Of the 25 cohorts we did obtain access to, 2 have fewer than 500 samples and were left out from out analyses due to low sample size. We detail this information in Supplemental Table S20.

Of the 23 MDD cohorts we used, 20 recorded endorsement of DSM-V criteria A for MDD from structured interviews. We assessed the percentage endorsement of each DSM-V criteria A symptom in these 20 cohorts, as well as the total number of symptoms endorsed per individual, per cohort (Supplemental Figure S14). We found that not all individuals indicated with a case status met the DSM-V A criteria for a "case" of MDD. We report the percentage of "DSM-MDD Cases" among indicated cases in the 20 cohorts in Supplemental Table S20.

We then obtained PRS using GWAS on each definition of depression using the Ricopili pipeline[34] (Methods), and compared the predictive power of PRS derived from different definitions of depression at each P-value threshold, where $P \in \{1e^{-4}, 0.001, 0.01, 0.05, 0.1, 0.2, 0.3, 0.4, 0.5, 1\}$. This is done instead of choosing the best P-value threshold per definition of depression, because our aim is not to achieve the best prediction, but to compare the genetic effects and their predictive power between the definitions of depression in the most unbiased way. We assess the prediction accuracy in terms of Nagelkerke's r2 (NKr2), area under the curve (AUC) of true positive against false positive rates, and variance of the MDD disease status explained by the PRS.

If the predictive power of PRS in each definition of depression in UKBiobank was specific to MDD, then greater percentages of DSM-MDD cases in the target PGC cohorts should give a higher prediction accuracy. If instead the PRS of the definition in question was not specific to MDD, then the percentages of DSM-MDD cases in the target PGC cohorts should not be expected to affect its prediction accuracies. We therefore assessed the specificity of PRS from each definition of depression in UKBiobank to MDD, by asking if it gives better predictions in PGC cohorts containing higher percentages of DSM-MDD cases. This is true across all P value thresholds for strictly defined, CIDI-based LifetimeMDD (Figure 7, Supplemental Table S22). As there is little predictive power across all definitions of depression at P-value thresholds $10^{-4}$ and 0.001, we do not show results at these thresholds for this analysis.

The apparent percentage of "DSM-MDD Cases" and PRS prediction accuracy in each cohort may be explained by the way its data was collected. For example, mmi2 and mmo4 used a questionnaire for assessing lifetime MDD in which some questions were not available and for other questions a choice had to be made about dichotomisation of the ordinal responses; a different choice of dichomisation would give a different result. As such, they have lower apparent percentage of "DSM-MDD Cases". gep3 is only cohort with non-screened controls[33] and may therefore give lower prediction accuracy. In contrast, nes1 has super-screened controls and therefore tends to give high out of sample prediction. In general, community-based cohorts are likely to have lower percentage of "DSM-MDD Cases", while they also have lower proportions of cases. For the same liability variance explained, cohorts with lowest case prevalence in the sample have lowest NKr2. We therefore present AUC as the measure to compare between cohorts.

**Effect of Sample size and specificity on prediction power**

The key dilemma facing researchers is whether there is more power from allowing a larger sample size through allowing more minimal phenotyping. We first performed the out-of-sample prediction analysis on the full sample size for each definition of depression in UKBiobank, and as expected, at a much larger sample size, minimal phenotyping, help-seeking based GPpsy predict much better than other definitions (Figure 7a, Supplemental Table S21).

We therefore asked if the high observed predictive power for GPpsy (at full cohort N=332,629) can be completely explained by its great sample size. Following Turley et al., 2018[35], the expected predictive power of the PRS from GPpsy is

$$\mathrm{E}(R_k^2) = \frac{(h^2)^2}{h^2 + V_k}$$

where $h^2$ is the SNP heritability of the trait, and $V_k$ is the variance of the normally distributed estimation error for individual polygenic scores

$$e_{k,i} \sim N(0, V_k)$$

in individual $i$, for each polygenic score $k \in \{GWAS_N\}$, where $N$ is the effective sample size in GWAS. For a binary case-control phenotype, $N = 4/(1/N_{cases} + 1/N_{controls})$. As $V_k$ is inversely proportional to $N$, $\mathrm{E}(R_k^2)$ increases with increasing $N$.

From Figure 7a and 7b we can see that an increase in sample size from $N_{DS} = 4/(1/7500 + 1/42500) = 25500$ (down-sampled, DS) to $N_{FC} = 4/(1/113260 + 1/219362) = 298777$ (full cohort, FC) resulted in a great increase in the predictive power of PRS from GPpsy. To find the relationship between increasing $N$ and power increase in GWAS and prediction, we use the relationship between $N$ and mean Chi-square ($\overline{\chi^2}$) statistic from GWAS. $\chi^2$ statistic of SNP j with LD score $l_j$ is related to $N$ by

$$\mathrm{E}(\chi_j^2 \mid l_j) = \frac{N h^2 l_j}{M} + Na + 1$$

Where $M$ is the total number of SNPs for which $h^2$ is defined, and $a$ is the variance due to biases such as population structure. $\mathrm{E}(\chi_j^2 \mid l_j)$ scales with $N$ if $M$ and $a$ are constant.

$$\frac{N_{FC}}{N_{DS}} \propto \frac{\overline{\chi^2{}_{FC}} - 1}{\overline{\chi^2{}_{DS}} - 1}$$

We found that across all definitions of depression, $\frac{\overline{\chi^2_{FC}}-1}{\overline{\chi^2_{DS}}-1}$ is indeed highly correlated with $\frac{N_{FC}}{N_{DS}}$ (Pearson r2 = 0.999, P = 5.50 x $10^{-7}$). $\frac{N_{FC}}{N_{DS}}$ has an effect of beta = 1.27 (se = 0.02) on $\frac{\overline{\chi^2_{FC}}-1}{\overline{\chi^2_{DS}}-1}$, as shown in Supplemental Figure S15.

We can therefore obtain theoretical $V_{FC}$ for GPpsy from $V_{DS}$ (which we can obtain using known $h^2$ and NKr2 for down-sampled GPpsy)

$$V_{FC} = V_{DS} \times 1.27 \frac{N_{DS}}{N_{FC}}$$

And obtain theoretical expected predictive power of the PRS from GPpsy at full cohort $N_{FC}$

$$\mathrm{E}(R^2_{FC}) = \frac{(h^2)^2}{h^2 + V_{FC}}$$

If the high predictive power of GPpsy can be completely explained by its large sample size, the theoretical expected predictive power $\mathrm{E}(R^2_{FC})$ should be roughly equal to that we obtain using real data. We see that this is true for GPpsy, where $\mathrm{E}(R^2_{FC})$ = 0.0185 and NKr2 for full cohort GPpsy = 0.0172.

We performed this analysis on all definitions of depression in UKBiobank, and found that for each definition, their increase in predictive power can be predicted and accounted for by the increase in N, as shown in Supplemental Figure S15. The Pearson correlation between predicted and actual NKr2 across all definitions was 0.989 (P = 4.46 x $10^{-5}$).

As such, the higher predictive power of GPpsy in Figure 7a (as compared to that in Figure 7b) can be explained entirely by its larger sample size.

We then asked what effective sample sizes other definitions of depression would need to achieve the same predictive power as GPpsy. We calculated their increase in predictive power for $N_x$ where $x > 25{,}500$ and $N_{DS}$ where $DS = 25{,}500$. We obtained the predicted NKr2 for $1.27\frac{N_x}{N_{DS}} \in \{1 \ldots 15\}$, and found that strictly defined LifetimeMDD needs a smaller $1.27\frac{N_x}{N_{DS}}$ to achieve the same NKr2 as GPpsy: while $N_x$= 274677 for a NKr2 of 0.0172 in full-cohort GPpsy (actual $N$ = 298677 for an actual sample size of 332,629), a smaller $N_x$= 129106 needed to achieve the same Nkr2 for LifetimeMDD.

### Supplemental Figures

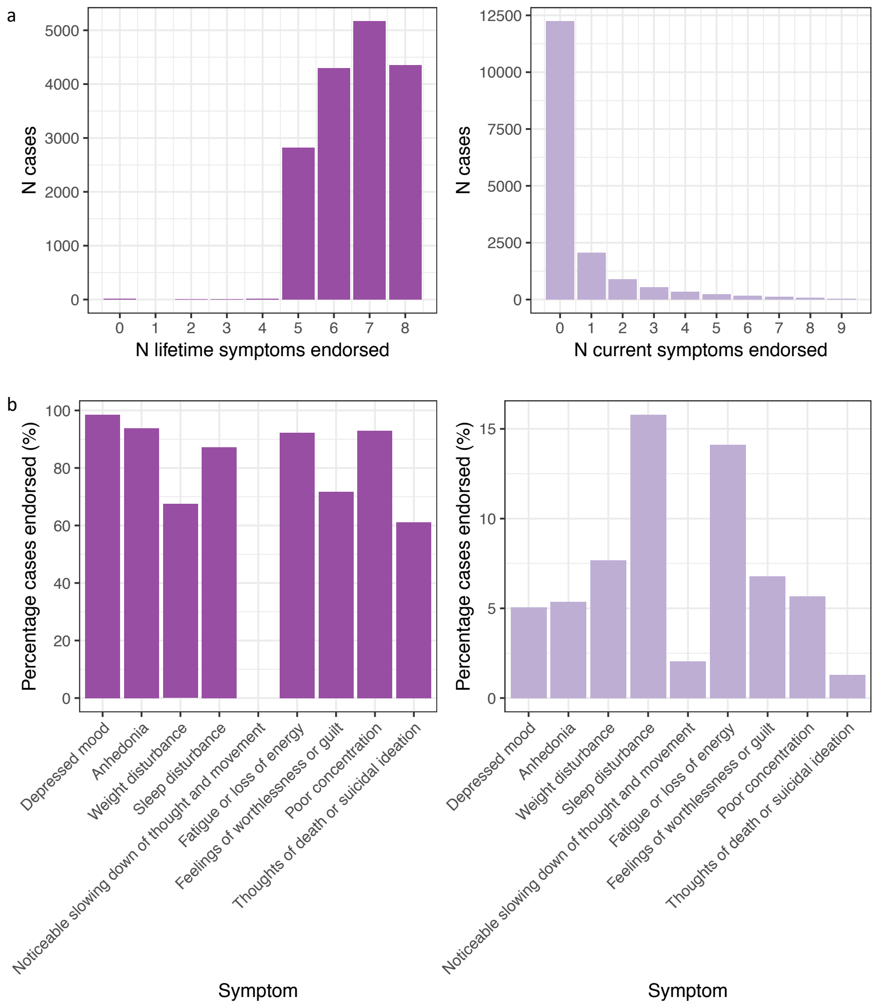

**Supplemental Figure S1: Symptoms of CIDI-based MDD LifetimeMDD in UKBiobank**

a) This figure shows the number of DSM-V criterion A symptoms endorsed by cases of LifetimeMDD defined in Supplemental Methods: the panel on the left (darker purple) shows the number of lifetime symptoms given in answer to the mental health questionnaire (MHQ); the panel on the right (lighter purple) shows the number of symptoms experienced over the two weeks leading up to the Touchscreen questionnaire. Those endorsing 0-4 lifetime symptoms in the left panel would have been classified as cases of LifetimeMDD because they endorsed 5 or more current symptoms in the right panel, including either “Depressed mood” or “Anhedonia”. b) This figure shows the percentage of cases of LifetimeMDD endorsing each of the DSM-V criterion A symptoms: the panel on the left (darker purple) shows the endorsement for each symptom in one’s life up to point of assessment; the panel on the right (lighter purple) shows the endorsement for having each symptom over the two weeks leading up to the Touchscreen questionnaire.

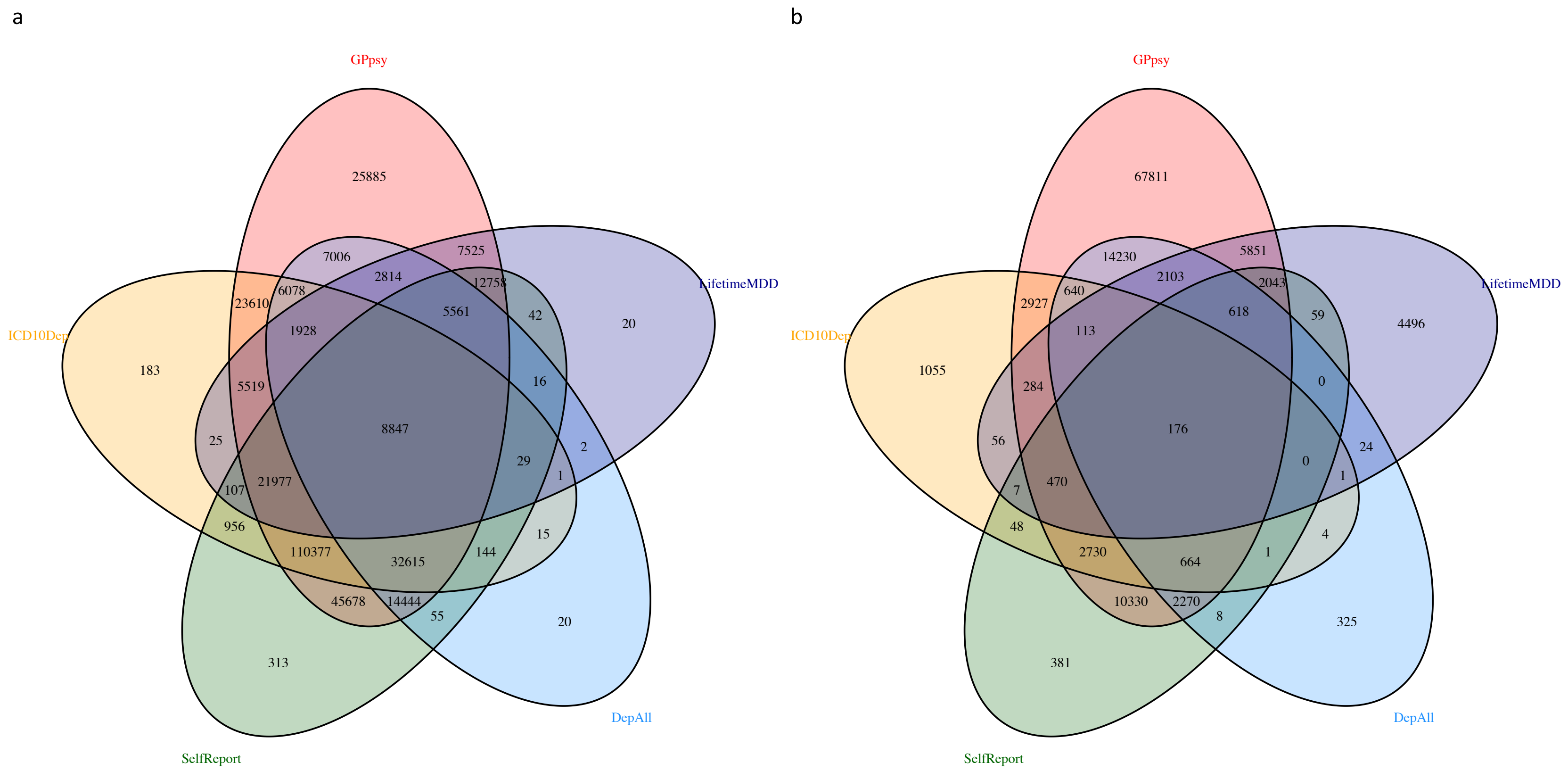

**Supplemental Figure S2: Overlap between definitions of MDD in UKBiobank**

a) This figure shows the number of overlapping cases between help-seeking (GPpsy in red), symptom (DepAll, in blue), self-report (SelfRepDep, in green), CIDI (LifetimeMDD, in purple) based definitions of MDD, as well as the electronic health record (EMR) based, ICD10 code derived depression (ICD10Dep, in orange). As not all individuals answered all questions necessary to assess whether they are a case or control in any of the definitions of MDD, we also show in b) the number of overlapping individuals (both cases and controls) who answered the question necessary to qualify as either cases or controls in each of the definitions of MDD (refer to main text and Figure 1 for details).

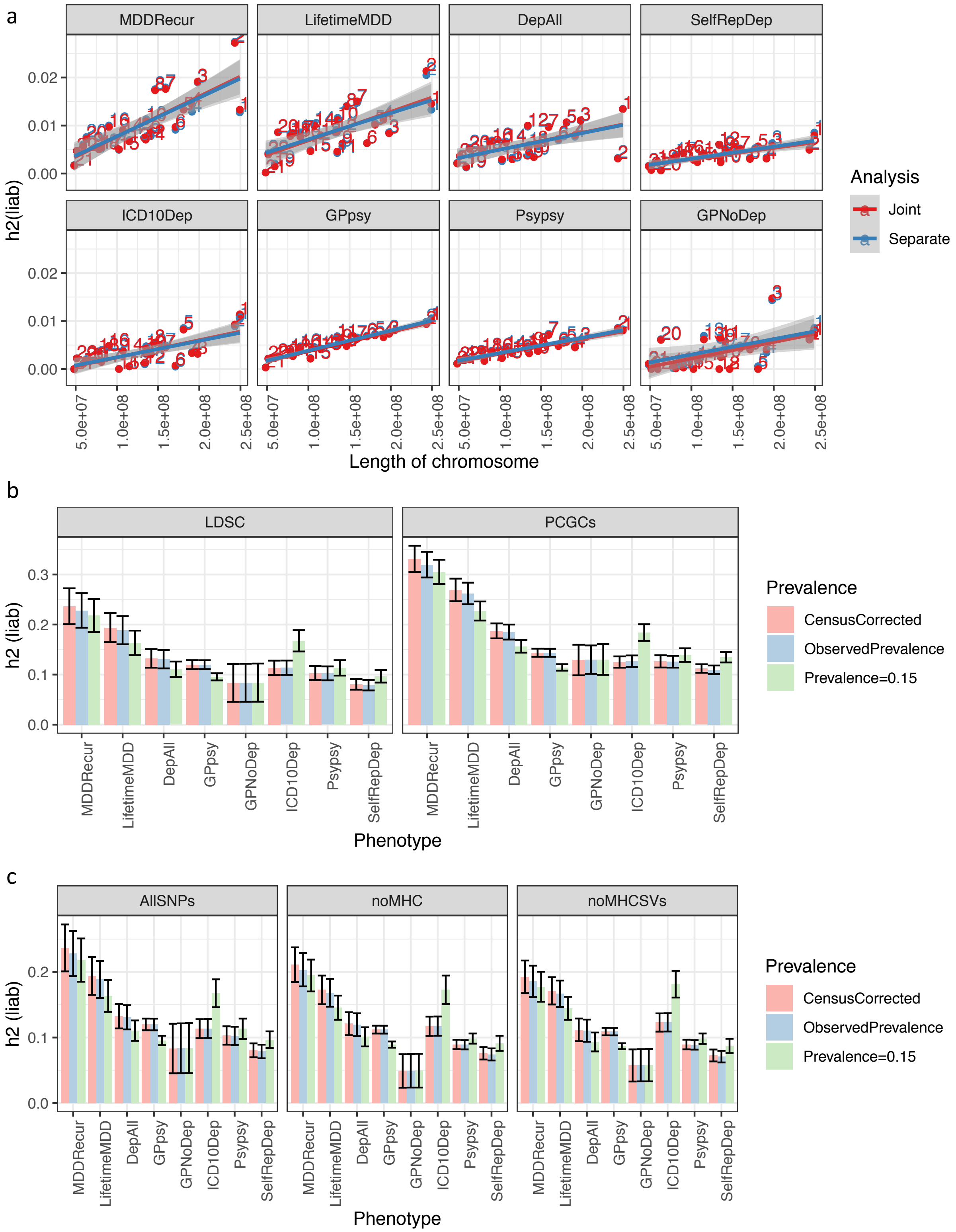

**Supplemental Figure S3: Effects of population structure and MHC/SVs**

a) This figure shows for each definition of depression in UKBiobank the estimates of $h^2_{SNP}$ (on the liability scale) contributed by each chromosome (obtained using BOLT-REML, see Supplemental Methods) estimated both jointly (in red) and separately (in blue), plotted against the lengths of the chromosomes, using both PCs obtained from the whole White-British cohort in UKBiobank. b) This figure shows for each definition of depression in UKBiobank estimtates of $h^2_{SNP}$ (on the liability scale) from PCGCs and LDSC: for each method, we obtained estimates using as population prevalence the observed prevalence (light blue), census corrected prevalence (light red) and assumed prevalence of 0.15 as per PGC1-MDD (light green). c) This figure shows for each definition of depression in UKBiobank the $h^2_{SNP}$ estimates from LDSC using all SNPs > 5% MAF (LDSC-AllSNPs), all SNPs > 5% MAF except those in the MHC region on chromosome 6:25-35MB (LDSC-noMHC), and all SNPs > 5% MAF except those in the MHC region and SVs[9]. The colour scheme reflects the population prevalence used in the analysis as explained in b).

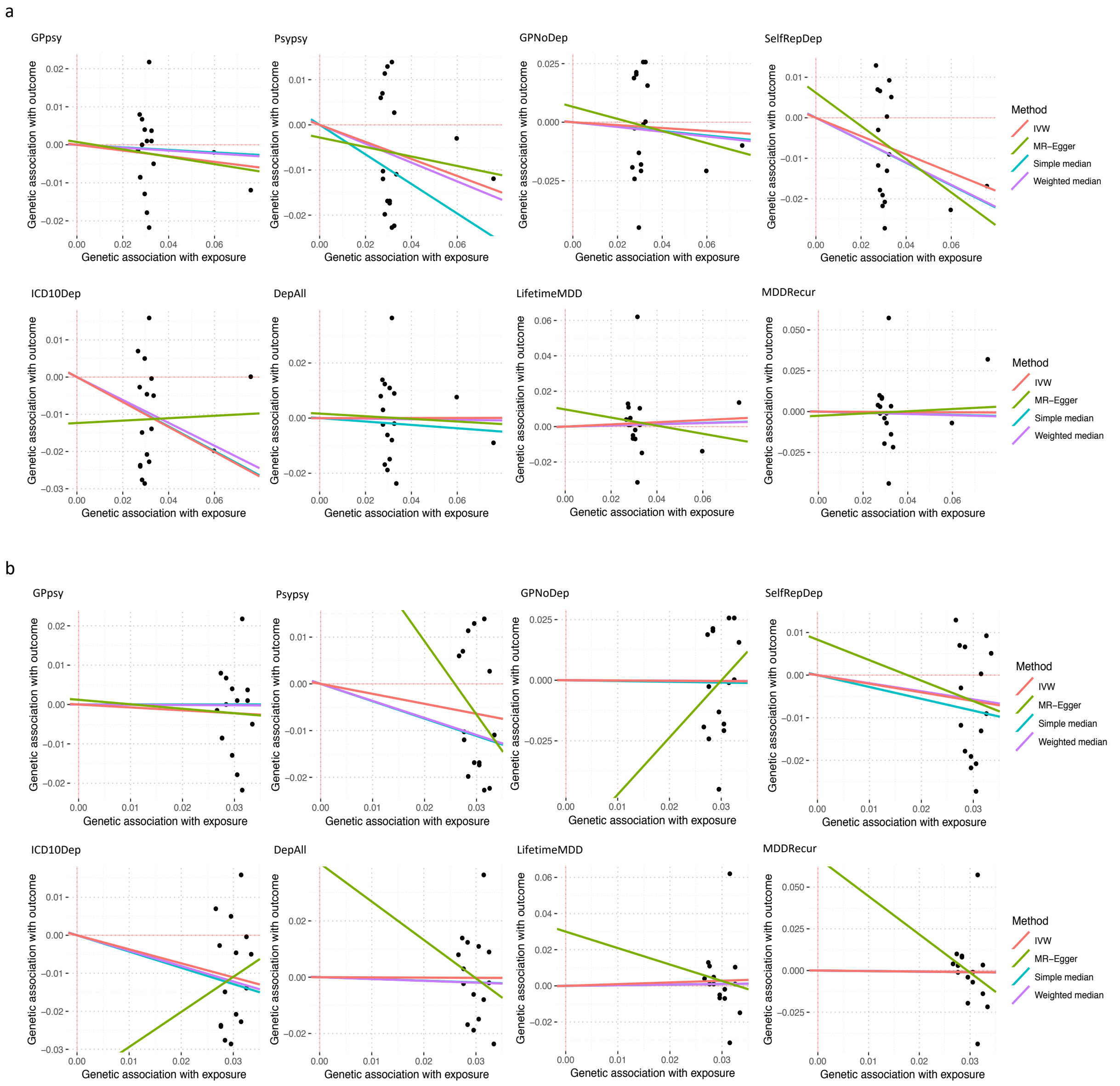

**Supplemental Figure S4: Effect of MHQ participation on definitions of depression**

a) This figure shows the results from Mendelian Randomization (MR) assessment of SNPs significantly associated with MHQ participation for their effects on definitions of depression in UKBiobank, using the following estimators: inverse variance weighted (IVW), MR-Egger, Simple median, and Weighted median. None of the estimators show significant causal effect between MHQ participation on definitions of depression. b) This figure shows the same results, removing two SNPs rs429358 and rs1261078 which have large effects on MHQ participation. Again no estimators show significant causal effects between MHQ participation on definitions of depression.

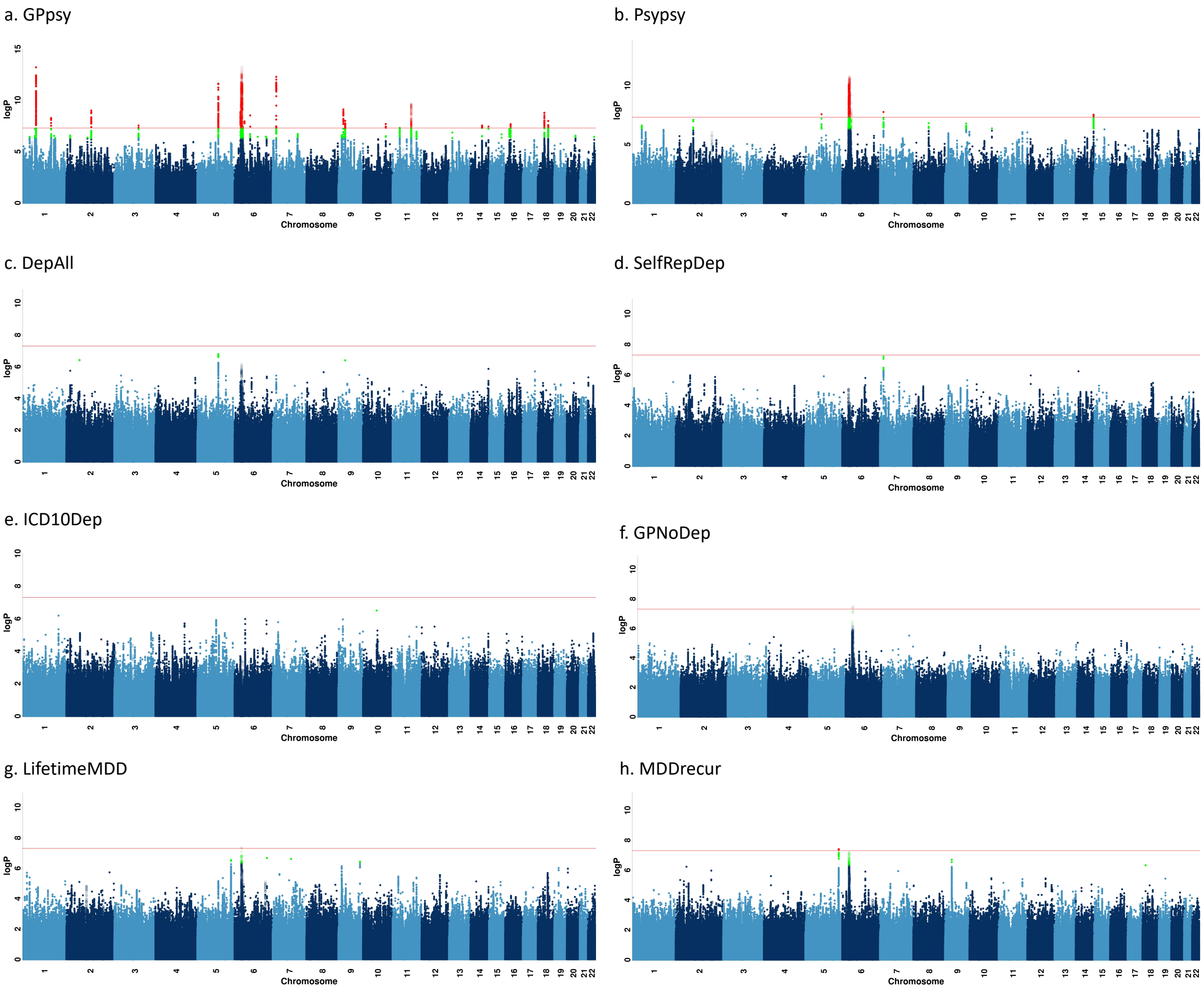

**Supplemental Figure S5: GWAS on definitions of depression using "clean" controls**

a) This figure shows Manhattan plots of GWAS on each definition of depression, performed with "clean" controls rather than all controls. We report all associations with P-values smaller than 5 x $10^{-8}$ as genome-wide significant (red). We indicated the SNPs in SVs and the MHC in all Manhattan plots as hollow points instead of solid points due to lack of control for population structure in these regions, and show all top SNPs within peaks (1MB regions) in Supplemental Table S11.

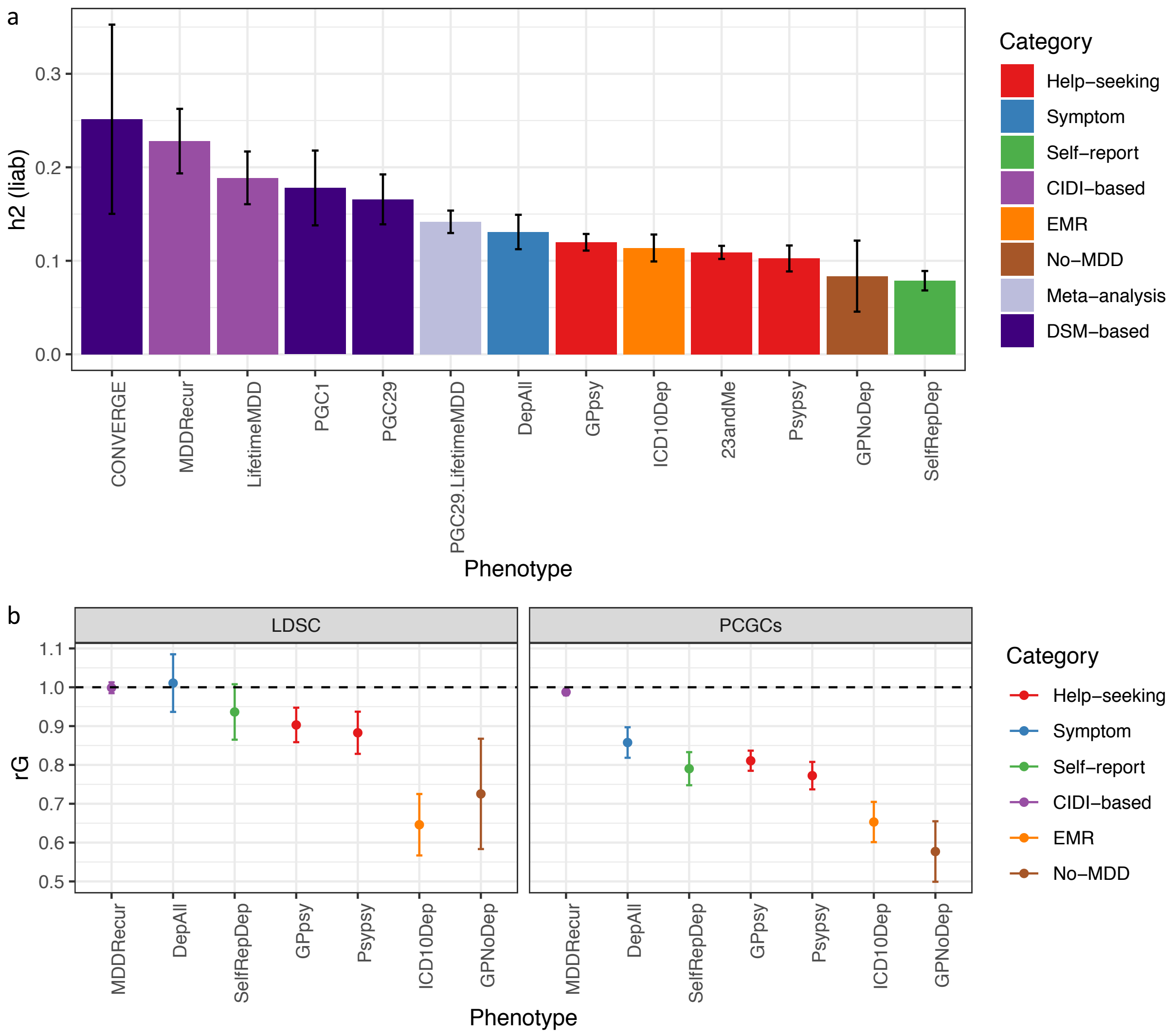

**Supplemental Figure S6: LDSC estimates of heritability and genetic correlation**

a) This figure shows the $h^2_{SNP}$ estimates of definitions of MDD in UKBiobank from LDSC using logistic regression summary statistics on all SNPs > 5% MAF (Methods). In the figure we also show LDSC estimates of heritabilities of previous studies of MDD including CONVERGE[26], PGC1-MDD[8], PGC29-MDD, a meta-analysis between CIDI-based LifetimeMDD and PGC29-MDD excluding individuals in UKBiobank (PGC29.LifetimeMDD), and 23andMe[27]. We show CIDI-based definitions in UKBiobank (in purple) show similar estimates to CONVERGE, symptom-based definitions in UKBiobank (in blue) are similar to PGC1-MDD, while help-seeking based (in red) definitions in UKBiobank are similar to 23andMe. b) This figure shows the genetic correlation between all definitions of MDD in UKBiobank against a CIDI-based definition of MDD (LifetimeMDD) estimated in LDSC and PCGCs (duplicated from Figure 3b for comparison).

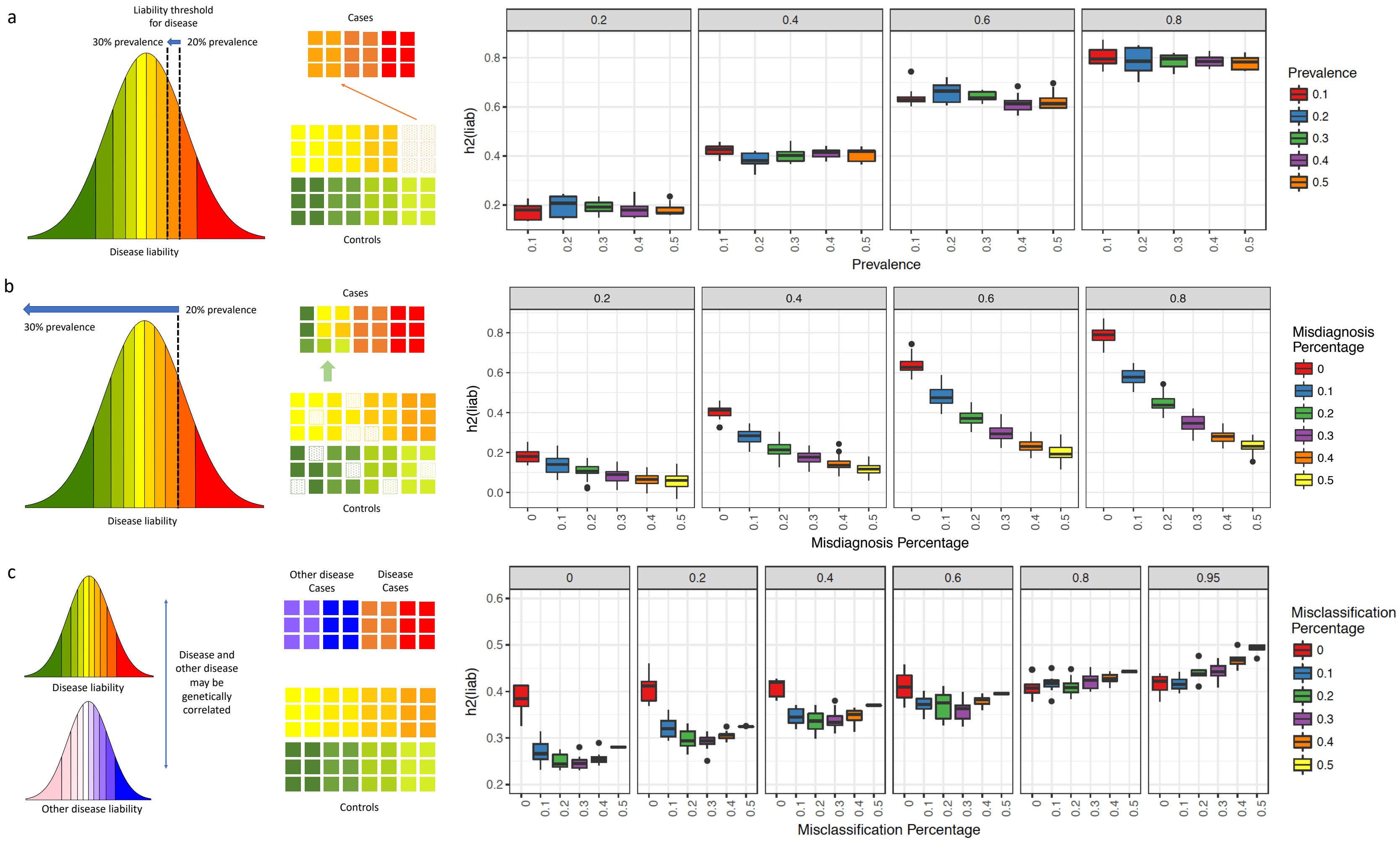
a
Liability threshold for disease
30% prevalence
20% prevalence
Disease liability
Cases
Controls
Prevalence
h2(liab)
b
30% prevalence
20% prevalence
Disease liability
Cases
Controls
Misdiagnosis Percentage
h2(liab)
c
Disease liability
Other disease liability
Disease and other disease may be genetically correlated
Other disease Cases
Disease Cases
Controls
Misclassification Percentage
h2(liab)

**Supplemental Figure S7: Simulations of misdiagnosis and misclassification**

a) $h^2_{SNP}$ changes with shifting of liability threshold. We simulated binary traits $y_i$, where $i \in \{1..10\}$ for simulated heritabilities $h_i^2 \in \{0.2, 0.4, 0.6, 0.8\}$ (in the grey bars above each panel), and plotted $h^2_{SNP}$ (using --pcgc option with --prevalence K in LDAK, plotted on the y axis) against the prevalence $K_i \in \{0.1, 0.2, 0.3, 0.4, 0.5\}$ of cases (defined as those individuals with liabilities greater than 1-Kth percentile). This figure shows liability scale $h^2_{SNP}$ does not change with shifting of liability threshold. b) $h^2_{SNP}$ changes with misdiagnosis of controls (those below the liability threshold) as cases (those above the liability threshold). We estimated simulated binary traits $y_i$, where $i \in \{1..10\}$ for simulated heritabilities $h_i^2 \in \{0.2, 0.4, 0.6, 0.8\}$ (in the grey bars above each panel), for each of which all cases (at prevalence $K_i$ =0.2) are correctly identified as cases. We then varied the numbers of random controls that are misdiagnosed as cases (plotted on the x axis as percentage of all identified cases), thereby increasing apparent prevalence of cases. We estimated $h^2_{SNP}$ (using --pcgc option with --prevalence K = new apparent case prevalence in LDAK), plotted on the y axis. The figure shows that liability scale $h^2_{SNP}$ is deflated with increasing percentage of controls being misdiagnosed as cases. c) $h^2_{SNP}$ changes with misclassification of cases of an “other” disease (with different liability distribution) as cases of a focal disease, when this “other” disease may be genetically correlated with the focal disease. We simulated binary traits $y_{i,1}$, where $i \in \{1..10\}$ for simulated $h_{i,1}^2 = 0.4$, for each of which all cases at prevalence $K_{i,1} = 0.2$ are correctly identified as cases. We then simulated genetically correlated binary traits $y_{i,2}$, where $i \in \{1..10\}$ of equal $h_{i,1}^2$ and prevalence as cases of $y_{i,1}$, for $rG_i \in \{0,0.2,0.4,0.6,0.8,0.95\}$. We varied the number of misclassifications from cases of $y_{i,2}$ in $y_{i,1}$(plotted on the x axis), thereby increasing apparent prevalence of cases. We then estimated $h^2_{SNP}$ (using --pcgc option with --prevalence K = new apparent case prevalence in LDAK), plotted on the y axis. This figure shows liability scale $h^2_{SNP}$ is deflated with increasing percentage of misclassification of cases of “other” disease as cases of focal disease, if rG between the two diseases are moderate to low.

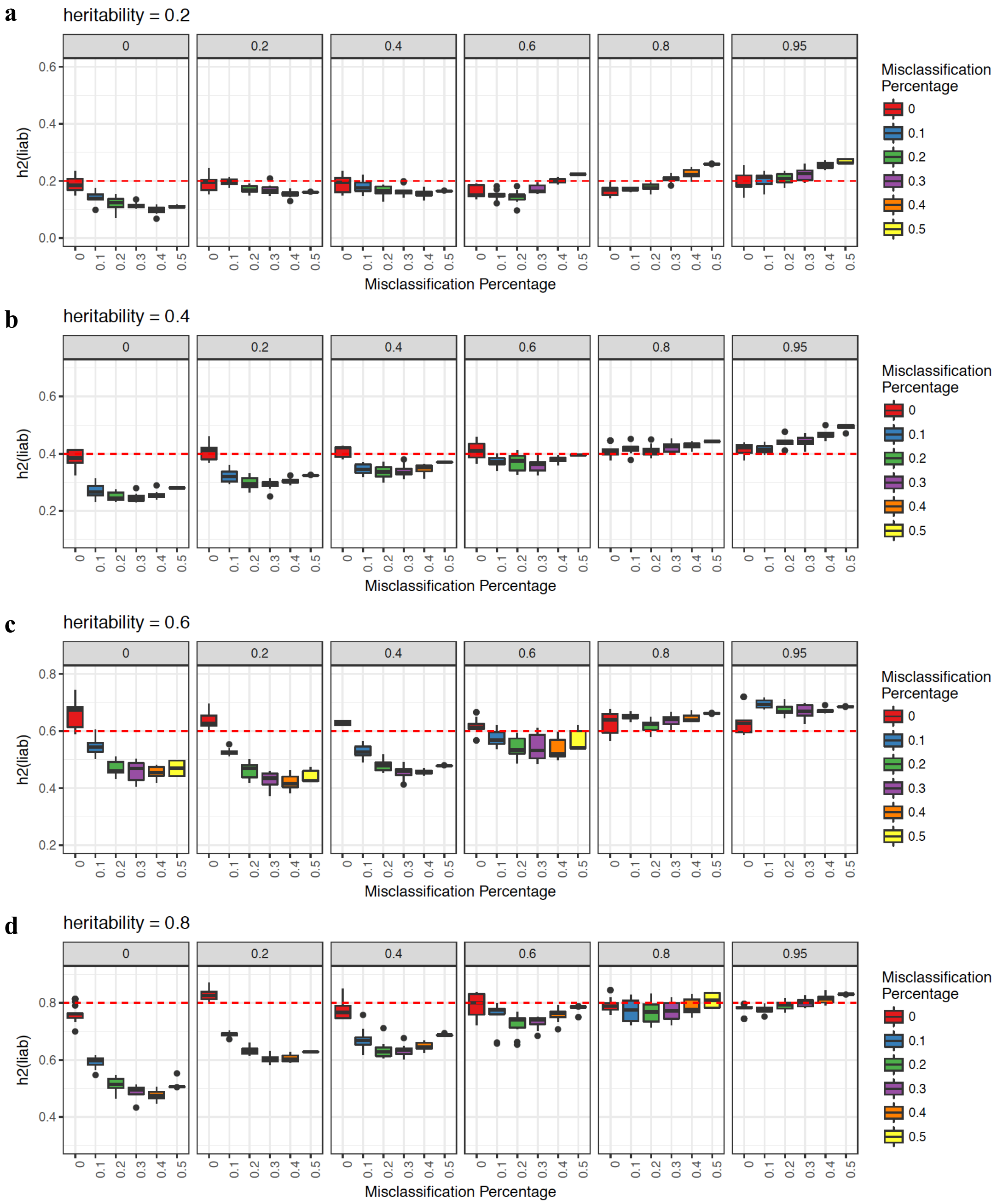

**Supplemental Figure S8: Simulations of misclassification at different heritabilities**

a-d) These figures shows the estimated $h^2_{SNP}$ (using --pcgc option with --prevalence K in LDAK, plotted on the y axis) of binary traits ($y_{i,1}$, where i $\in \{1..10\}$) with simulated $h^2_{i,1}$ 0.2, 0.4, 0.6, and 0.8, for each of which all cases (at prevalence $K_{i,1}$=0.2) are correctly identified as cases, while varying numbers of cases misclassified from a genetically correlated binary trait ($y_{i,2}$, where $i \in \{1..10\}$) of equal $h^2_{i,1}$ and prevalence as cases of $y_{i,1}$. Genetic correlations between $y_{i,1}$and $y_{i,2}$ ($rG_i \in \{0, 0.2, 0.4, 0.6, 0.8, 0.95\}$) are shown in the grey bars above each panel.

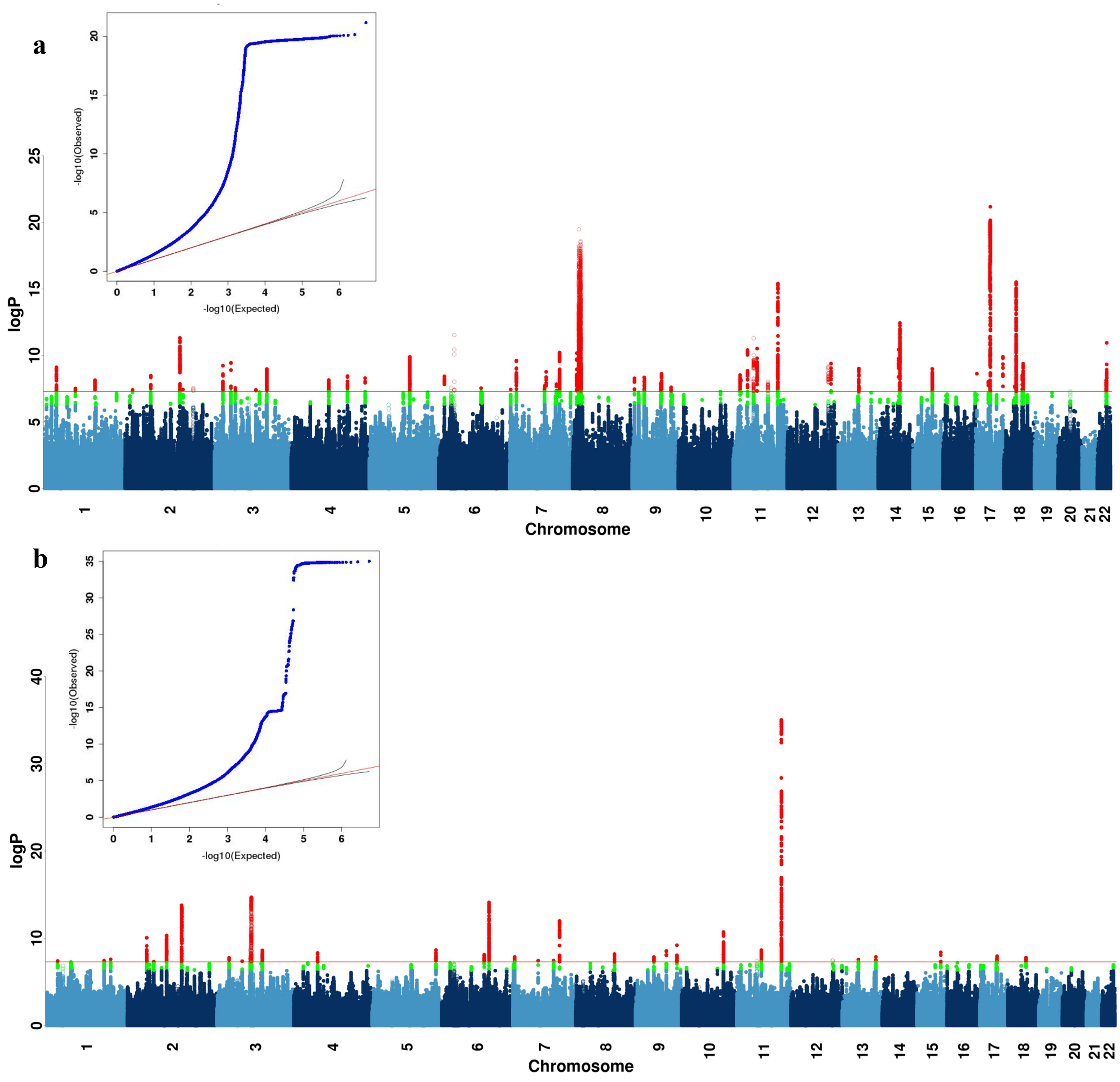

**Supplemental Figure S9: GWAS on neuroticism and smoking in UKBiobank**

a-b) This figure shows the Manhattan plot of neuroticism score (data field 20127, quantitative trait from 0 to 12) in 274,107 individuals and ever smoked status (data field 20160, binary trait of 0 for "No", and 1 for "Yes") in 336,066 individuals in UKBiobank using linear regression on all 8,968,716 common SNPs (MAF > 5% in all 337,198 White-British, unrelated samples) for all the above analyses in PLINK (version 1.9) [36] with 20 PCs and genotyping array as covariates. We report all associations with P-values smaller than $5 \times 10^{-8}$ as genome-wide significant (red). We indicated the SNPs in SVs and the MHC in all Manhattan plots as hollow points instead of solid points due to lack of control for population structure in these regions, and show all top SNPs within peaks (1MB regions) in Supplemental Tables S10-11.

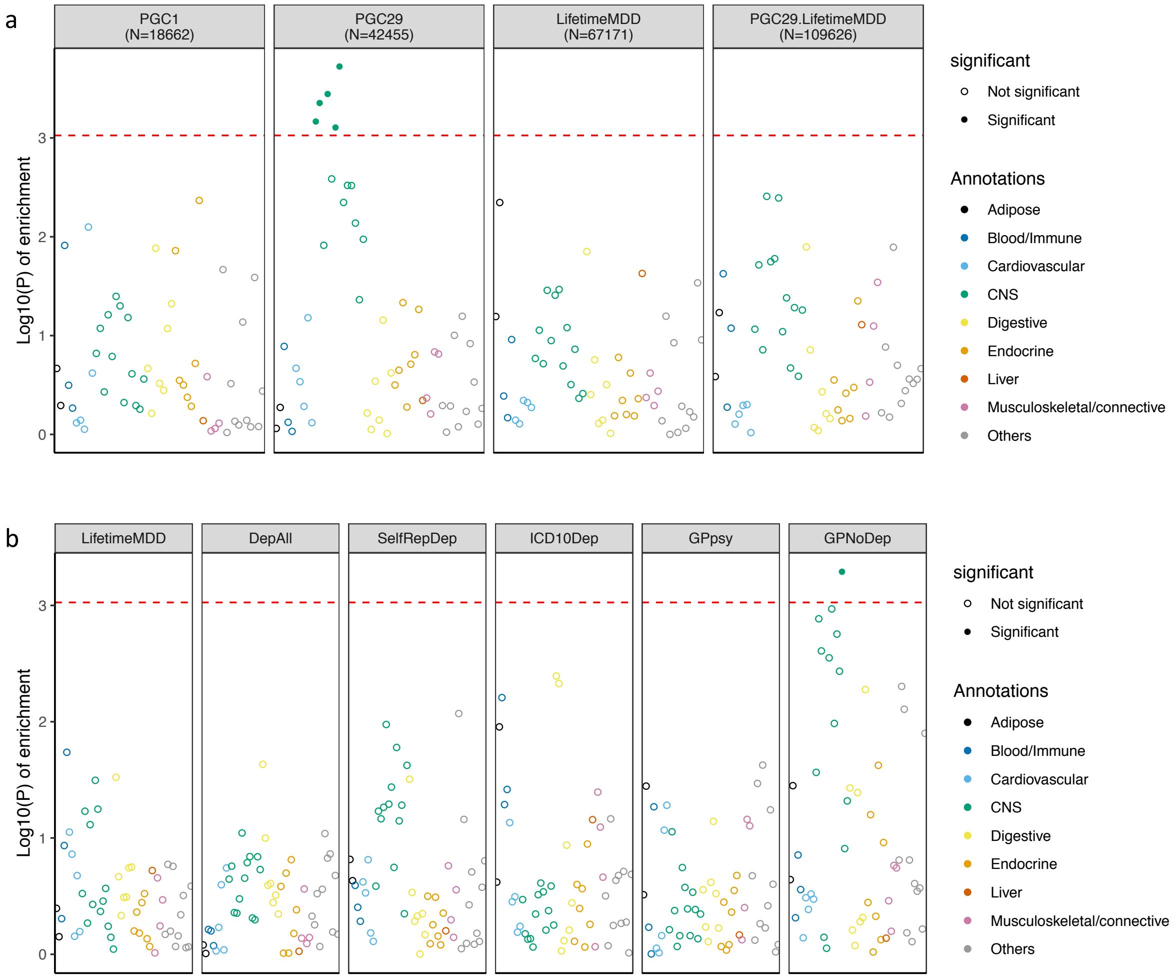

**Supplemental Figure S10: LDSC-SEG analysis of tissue-specific enrichment of $h^2_{SNP}$**

a) This figure shows -log10(P) of enrichment in heritability in genes specifically expressed in 44 GTEx tissues, estimated using partitioned heritability in LDSC-SEG, on LifetimeMDD (N = 67,171), PGC1-MDD (N = 18,759), PGC29 (N = 42,455) and a meta-analysis of LifetimeMDD and PGC29 (N=109,626, PC29.LifetimeMDD, Methods). While PGC29 shows CNS enrichment, neither LifetimeMDD nor the meta-analysis shows the same enrichment. This suggests sample size and differences in genetic architecture and cohort heterogeneity affects results from LDSC-SEG. b) This figure shows the same analysis performed on down-sampled data for each definition of depression. Each definition is randomly down-sampled to 7,500 cases and 42,500 controls, a constant prevalence of 0.15, to remove confounding from sample size and difference in statistical power on the enrichment analysis. This figure shows that at equal sample size and prevalence, GPNoDep (no-MDD Help-seeking phenotype) is the only one showing CNS enrichment, suggesting it may be driving the CNS enrichment signal in GPpsy in Figure 5.

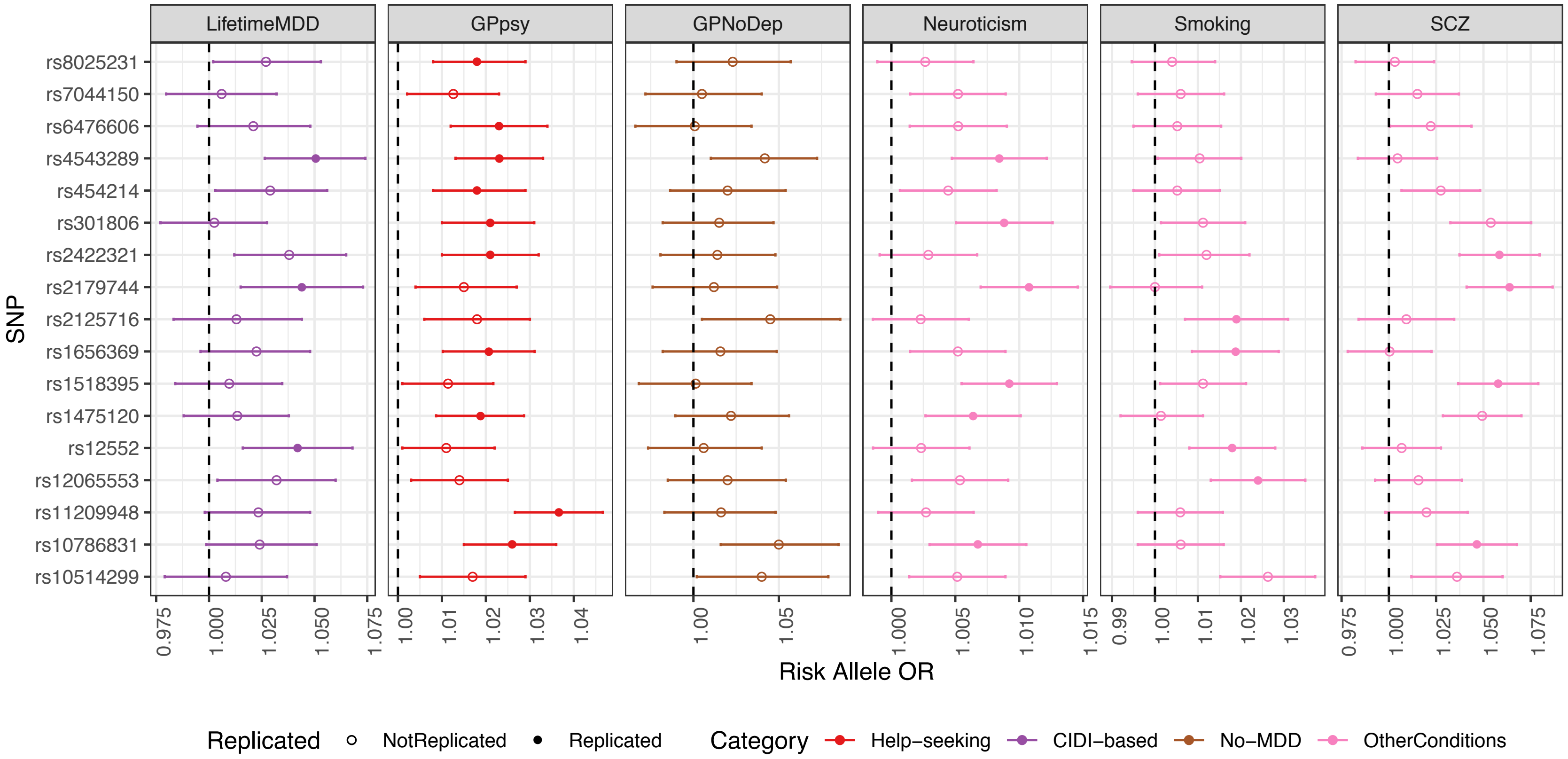

**Supplemental Figure S11: GWAS hits from 23andMe are not specific to MDD**

This figure shows the odds ratios of risk alleles (Risk Allele ORs) at 17 loci significantly associated with help-seeking based definitions of MDD in 23andMe[27], in GWAS conducted on CIDI-based (LifetimeMDD, in purple), help-seeking (GPpsy in red) and no-MDD (GPNoDep, in orange) based definitions of MDD, as well as conditions other than MDD: neuroticism, smoking and SCZ (all in brown). SNPs missing in each panel are not tested in the respective GWAS. For clarity of display, scales on different panels vary to accommodate the different magnitudes of ORs of SNPs in different conditions. ORs at all 17 loci are highly consistent across phenotypes, regardless of whether it is a definition or MDD or a risk factor or condition other than MDD. All results are shown in Supplemental Table S15.

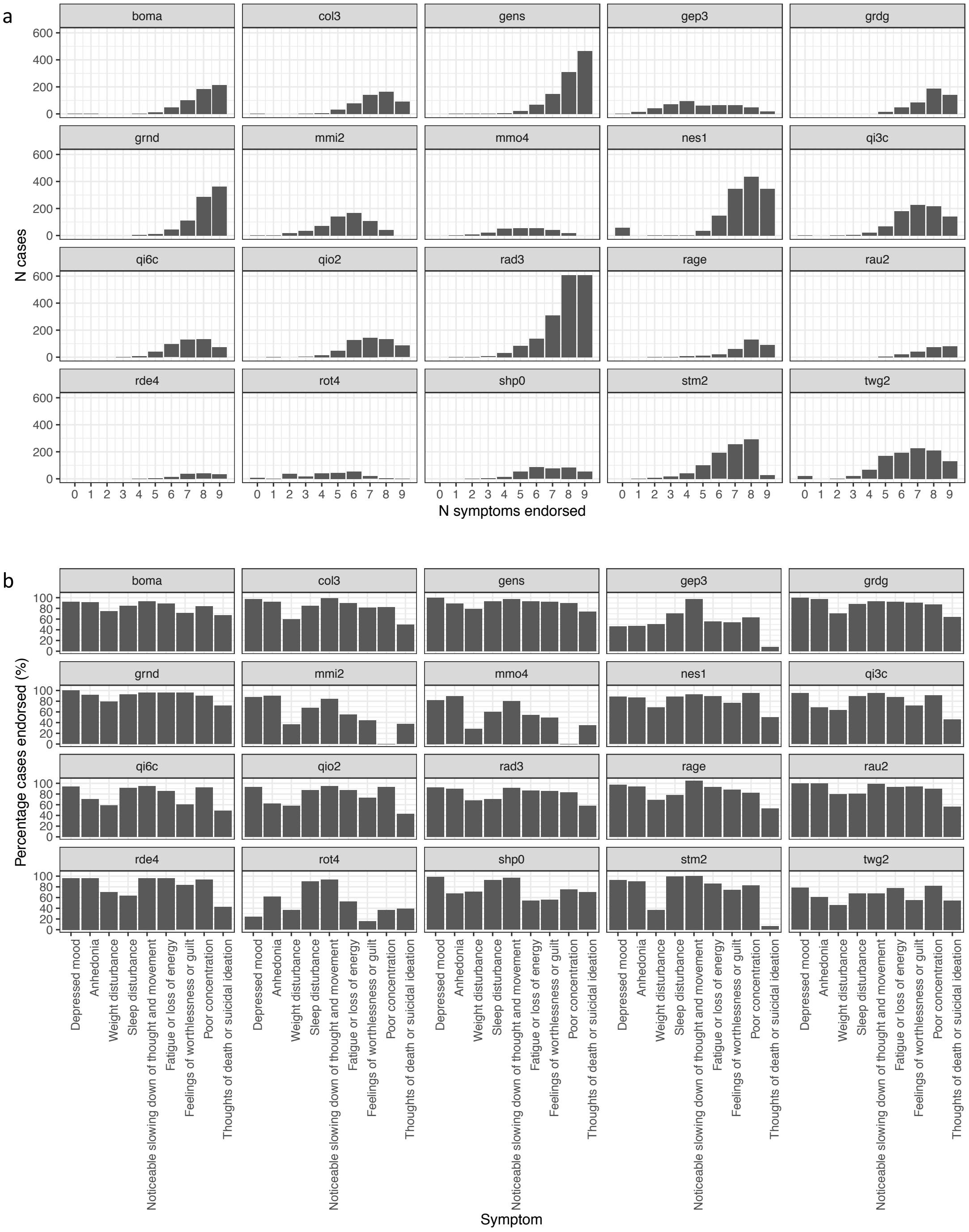

**Supplemental Figure S12: Symptoms of MDD cases in PGC cohorts**

a) The number of DSM-5 criterion A symptoms, defined as described in Supplemental Methods, endorsed by cases of MDD in 23 PGC cohorts. Those endorsing 0-4 lifetime symptoms, or more symptoms without endorsing “Depressed mood” or “Anhedonia”, would not be included in the “DSM-MDD” cases we define in each PGC cohort, as shown in Supplemental Table S20. b) The percentage of cases of MDD in PGC cohorts endorsing each of the DSM-5 criterion A symptoms.

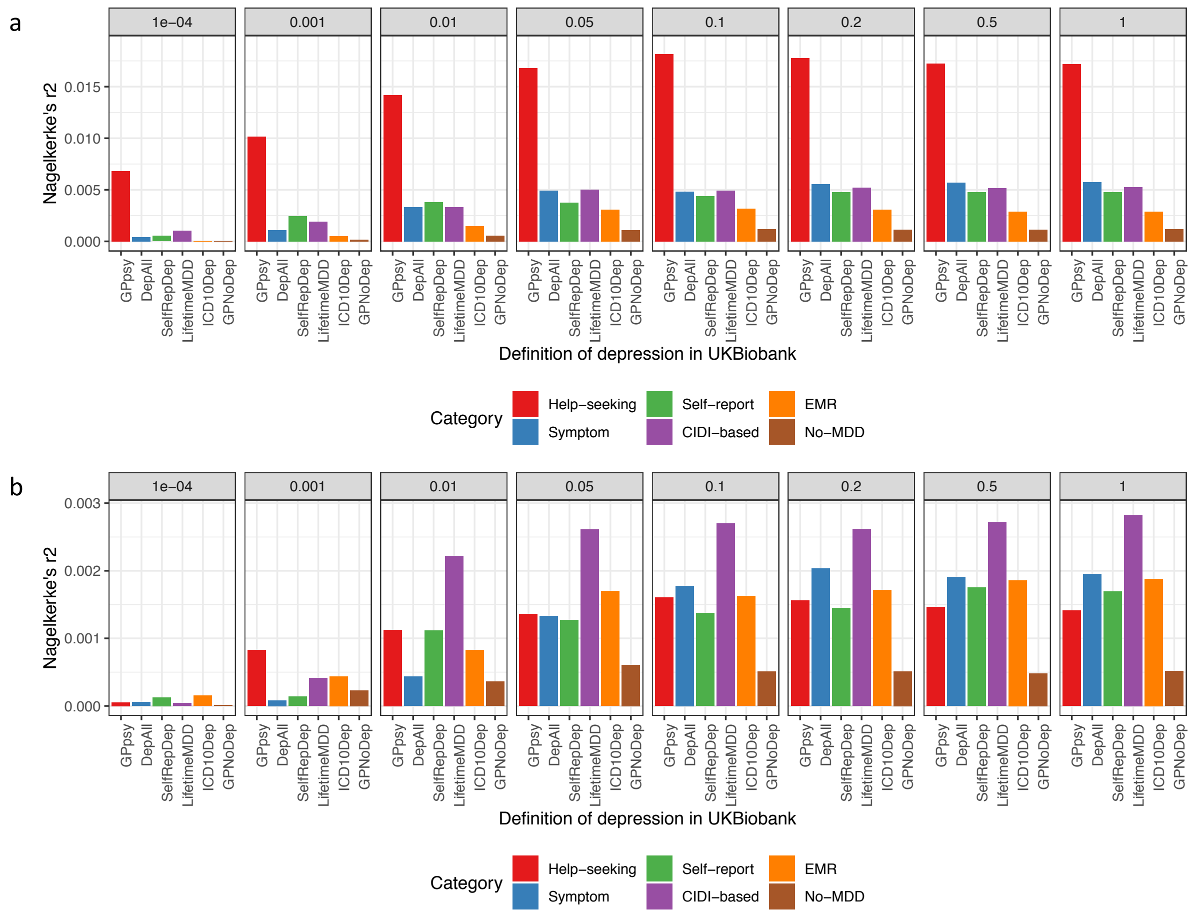

**Supplemental Figure S13: Out-of-sample prediction in PGC cohorts**

a) This figure shows the Nagelkerke's r2 of polygenic risk scores (PRS) calculated for each definition of depression in UKBiobank and MDD status indicated in 19 PGC29-MDD cohorts, while controlling for cohort specific effects. PRS were calculated using effect sizes at independent (LD r2 < 0.1) SNPs passing P value thresholds 10-4, 0.001, 0.01, 0.05, 0.01, 0.2, 0.5 and 1 respectively, in GWAS performed on all definitions of depression in UKBiobank. b) This figure shows the same analysis performed on down-sampled data (7,500 cases, 42,500 controls) for each definition of depression.

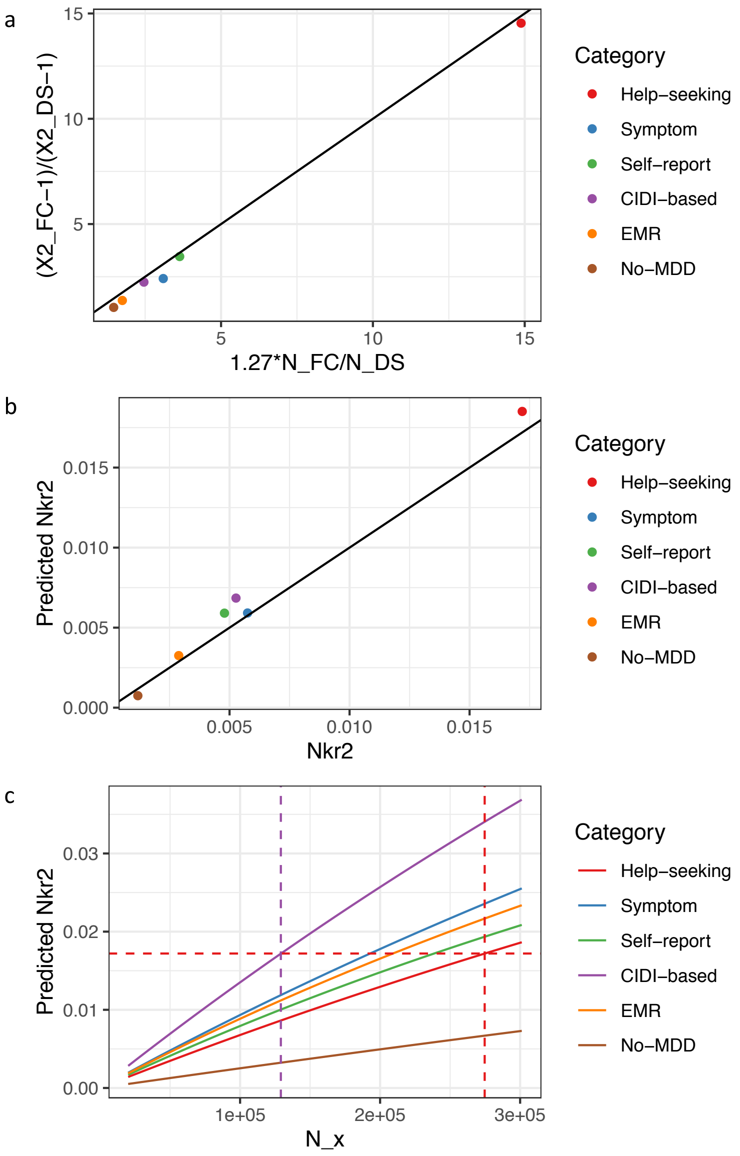

**Supplemental Figure S14: Relationship between effective sample size and prediction accuracy**

a) This figure shows the relationship between the ratio of effective sample sizes between the full cohort ($N_{FC}$) and down-sampled ($N_{DS}$) data for each definition of depression and the ratio of their mean Chi-square ($\chi^2$) statistic from GWAS, with black line x=y for reference. Across all definitions of depression, $\frac{\overline{\chi^2{}_{FC}-1}}{\overline{\chi^2{}_{DS}-1}}$ is highly correlated with $\frac{N_{FC}}{N_{DS}}$ (Pearson r2 = 0.999, P = 5.50 x $10^{-7}$), and $\frac{N_{FC}}{N_{DS}}$ has an effect of beta = 1.27 (se = 0.02) on $\frac{\overline{\chi^2{}_{FC}-1}}{\overline{\chi^2{}_{DS}-1}}$. b) This figure shows the Nagelkerke's r2 (Nkr2) for MDD status in PGC29 cohorts predicted for PRS of different definitions of depression at $N_{FC}$, plotted against their respective empirical Nkr2 at $N_{FC}$, both at P value threshold =1. The Pearson correlation $r^2$ between predicted and actual NKr2 across all definitions were 0.989 (P = 4.46 x $10^{-5}$). c) This figure shows for each definition of depression the effective sample size $N_X$ required for each predicted Nkr2 in out-of-sample prediction of MDD status in PGC29 cohorts. While $N_x$= 274677 for GPpsy to achieve a Nkr2 of 0.0172, a smaller $N_x$= 129106 is needed to achieve the same Nkr2 for LifetimeMDD.

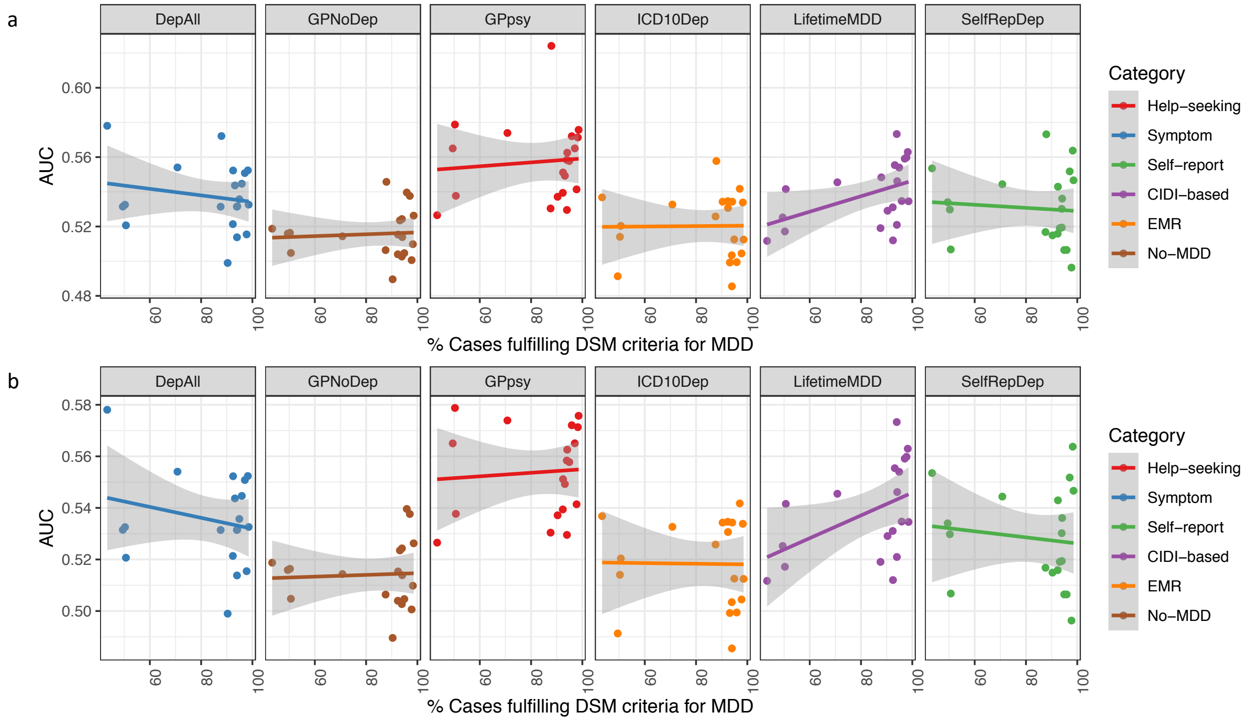

**Supplemental Figure S15: Prediction accuracy in cohorts with different percentage DSM MDD cases**

a) This figure shows the area under the curve (AUC) of polygenic risk scores (PRS) calculated for each definition of depression in UKBiobank and MDD status indicated in 20 PGC29-MDD cohorts at P-value threshold of 1 (using all SNPs after LD-clumping, see results at all P value thresholds in Supplemental Table S23), plotting AUC for each cohort against their respective percentage of cases fulfilling DSM-5 criteria A for MDD (see Supplemental Table S21). It shows that strictly defined CIDI-based LifetimeMDD is the only definition of depression in UKBiobank that shows increases in AUC as percentage of cases fulfilling DSM-5 criteria A for MDD in PGC cohorts increases, despite not giving the highest AUC. b) This figure shows the same analysis removing the PGC29-MDD cohort rad3, which is the outlier giving AUC > 0.6 in GPpsy in a). As this is a UK-based cohort, it is possible it contains relatives of individuals in UKBiobank that upwardly biased prediction accuracy in it. For all analysis shown in Figure 7, Supplemental Figure S13 and S14, and Supplemental Table S23, we have removed this cohort.

**Supplemental Tables**

| ABBREVIATION | PHENOTYPE | COLLECTION STRATEGY | STUDY TYPE | REFERENCE | N | SAMPLE PREV | POPULATION PREV | PREVALENCE REFERENCE |
|---|---|---|---|---|---|---|---|---|
| CONVERGE | MDD | Hospital based psychiatrist diagnosis | case-ascertained, screened-controls | CONVERGE Consortium, 2015 | 10640 | 0.50 | 0.08 | CONVERGE Consortium, 2015 |
| PGC1-MDD | MDD | Structured telephone interviews | case-ascertained, screened-controls (mega-analysis) | MDD Working Group of the Psychiatric Genomics Consortium, 2013 | 18759 | 0.50 | 0.15 | Kessler et al., 2003 |
| PGC29 | MDD | Structured telephone interviews/electronic health records | case-ascertained, some screened-controls (meta-analysis) | MDD Working Group of the Psychiatric Genomics Consortium, 2018 | 42455 | 0.40 | 0.15 | Kessler et al., 2003 |
| 23andMe | MDD | Minimal phenotyping: self-report via single question in questionnaire | unascertained population cohort | Hyde et al., 2016 | 307354 | 0.25 | 0.25 | Hyde et al., 2016 |
| ADHD | ADHD | Hospital based psychiatrist diagnosis | case-ascertained, screened-controls (mega-analysis) | Demontis et al., 2017 | 53293 | 0.36 | 0.05 | Demontis et al., 2017 |

| | | | | | | | | |
|---|---|---|---|---|---|---|---|---|
| BIP | BIP | Hospital based psychiatrist diagnosis | case-ascertained, 33% screened-controls (mega-analysis) | Bipolar Disorder Working Group of the Psychiatric Genomics Consortium, 2011 | 16731 | 0.45 | 0.025 | Merikangas et al., 2011 |
| SCZ | SCZ | Hospital based psychiatrist diagnosis /semi-structured interviews | case-ascertained, population-controls; trios (mega-analysis) | Schizophrenia Working Group of the Psychiatric Genomics Consortium, 2014 | 150064 | 0.25 | 0.01 | Schizophrenia Working Group of the Psychiatric Genomics Consortium, 2014 |
| AUT | AUT | Hospital based psychiatrist diagnosis | trios; case-ascertained, population-controls (mega-analysis) | Cross-Disorder Group of the Psychiatric Genomics Consortium, 2013 | 10610 | 0.50 | $9x10^{-4}$ | Cross-Disorder Group of the Psychiatric Genomics Consortium, 2013 |

**Supplemental Table S1: Other studies referenced in this paper**

This table shows the studies from which we obtain GWAS summary statistics for MDD and other psychiatric conditions that we reference in this paper. The columns are abbreviations of the studies used in this paper, the disease phenotype in question, their case collection strategy, their study type where known, the study reference, number of samples involved, case prevalence in sample, population prevalence of disease phenotypes, and reference for population prevalence of disease prevalence if different from the reference of the study itself.

| DSM CRITERIA | DATA FIELD | QUESTION ON LIFETIME MDD | QUALIFYING ANSWERS | N_YES | N_NO |
|---|---|---|---|---|---|
| A1 | 20446 | Have you ever had a time in your life when you felt sad, blue, or depressed for two weeks or more in a row? | Yes | 59409 | 50018 |
| A2 | 20441 | Have you ever had a time in your life lasting two weeks or more when you lost interest in most things like hobbies, work, or activities that usually give you pleasure? | Yes | 42565 | 66830 |
| A3 | 20536 | Did you gain or lose weight without trying, or did you stay about the same weight? | Gained weight, Lost weight, Both gained and lost some weight during the episode | 30875 | 21324 |
| A4 | 20532 | Did your sleep change? | Yes | 41955 | 10714 |
| A6 | 20449 | Did you feel more tired out or low on energy than is usual for you? | Yes | 45034 | 10072 |
| A7 | 20450 | People sometimes feel down on themselves, no good, worthless. Did you feel this way? | Yes | 28865 | 28472 |
| A8 | 20435 | Did you have a lot more trouble concentrating than usual? | Yes | 42568 | 11592 |
| A9 | 20437 | Did you think a lot about death - either your own, someone else's or death in general? | Yes | 29946 | 27856 |
| **DSM CRITERIA** | **DATA FIELD** | **QUESTION ON CURRENT MDD: Over the last 2 weeks, how often have you been bothered by any of the following problems?** | **QUALIFYING ANSWERS** | **N_YES** | **N_NO** |
| A1 | 20510 | Feeling down, depressed, or hopeless | Nearly every day | 1449 | 108261 |

| | | | | | |
|---|---|---|---|---|---|
| A2 | 20514 | Little interest or pleasure in doing things | Nearly every day | 1670 | 108040 |
| A3 | 20511 | Poor appetite or overeating | Nearly every day | 2633 | 107077 |
| A4 | 20517 | Trouble falling or staying asleep, or sleeping too much | Nearly every day | 8616 | 101094 |
| A5 | 20518 | Moving or speaking so slowly that other people could have noticed? Or the opposite - being so fidgety or restless that you have been moving around a lot more than usual | Nearly every day | 650 | 109060 |
| A6 | 20519 | Feeling tired or having little energy | Nearly every day | 5968 | 103742 |
| A7 | 20507 | Feeling bad about yourself or that you are a failure or have let yourself or your family down | Nearly every day | 2166 | 107544 |
| A8 | 20508 | Trouble concentrating on things, such as reading the newspaper or watching television | Nearly every day | 1769 | 107941 |
| A9 | 20513 | Thoughts that you would be better off dead or of hurting yourself in some way | Nearly every day | 357 | 109353 |

**Supplemental Table S2: CIDI-based criteria for lifetime and current MDD**

This table lists questions in the online mental health follow up questionnaire in UKBiobank forming the DSM-V diagnostic criteria for lifetime MDD[5] (Question on lifetime MDD) and current MDD (Question on current MDD), along with their data field in UKBiobank data showcase (datafield), the DSM criteria they correspond to, and the multiple-choice answer to each of the questions which indicates a subject as self-reporting having the symptom.

| Other criteria | | GPNoDep | | |
|---|---|---|---|---|
| | | Controls | Cases | Total |
| **MHQ** | Answered MHQ | 13514 | 1541 (10.2%) | 15055 |
| | Did not answer MHQ | 35979 | 7091 (16.5%) | 43070 |
| **Self-report** | Self-reported illness | 36474 | 7139 (16.4%) | 43613 |
| | Did not self-report illness | 13019 | 1439 (10.0%) | 14458 |

**Supplemental Table S3: Characterizing no-MDD definition GPNoDep**

This table shows the number of cases and controls in the no-MDD definition GPNoDep that answered or did not answer either the mental health questionnaire (MHQ) in the online mental health follow-up, or self-reported any illness in the verbal interview. The proportion of GPNoDep cases who answered the self-reported non-cancer illnesses during the verbal interview (16.4%) is significantly different from the proportion who did not answer (10.0%, Chisq test $P < 10^{-16}$, df = 1). Similarly, the proportion of GPNoDep cases who asnwered the MHQ (10.2%) is significantly different from the proportion who did not (16.5%, Chisq test $P < 10^{-16}$, df = 1).

| CENTRE | DEFINITION OF DEPRESSION (N_CASES/N_CONTROLS) | | | | | | | |
|---|---|---|---|---|---|---|---|---|
| | CIDI-BASED | | SYMPTOM | SELF-REPORT | HELP-SEEKING | | NO-MDD | EMR |
| | LifetimeMDD | MDDRecur | DepAll | SelfRepDep | GPpsy | Psypsy | GPNoDep | ICD10Dep |
| Barts | 399/1075 | 274/1040 | NA | 292/3623 | 1941/3795 | 1025/4728 | NA | 146/3154 |
| Birmingham | 869/2333 | 563/2239 | 3198/7995 | 816/10902 | 5536/10139 | 1922/13782 | 1186/6769 | 450/9247 |
| Bristol | 1613/5047 | 1001/4880 | 1649/4643 | 1534/21046 | 9817/20411 | 2960/27335 | 655/3972 | 738/18452 |
| Bury | 826/2436 | 512/2347 | NA | 1506/15186 | 6971/13005 | 2118/17939 | NA | 708/13533 |
| Cardiff | 554/1660 | 361/1599 | NA | 603/8663 | 4357/8239 | 1345/11285 | NA | 364/8022 |
| Croydon | 801/2937 | 500/2842 | 2744/7922 | 811/10644 | 4706/9985 | 1791/12915 | 1074/6821 | 360/8331 |
| Edinburgh | 568/2092 | 336/2033 | NA | 551/7368 | 3851/8325 | 1305/10889 | NA | 106/5784 |
| Glasgow | 480/1605 | 309/1565 | NA | 771/8505 | 4550/7961 | 1553/10998 | NA | 158/7406 |
| Hounslow | 743/3060 | 472/2954 | 2489/7862 | 619/9990 | 4401/9948 | 1741/12642 | 1055/6775 | 309/7978 |
| Leeds | 1503/4312 | 932/4146 | NA | 2058/20840 | 10701/20079 | 3379/27457 | NA | 805/18691 |
| Liverpool | 1080/3024 | 700/2916 | 2651/7765 | 1609/16421 | 7792/14444 | 2377/19930 | 1221/6500 | 875/14622 |
| Manchester | 469/1282 | 321/1218 | NA | 821/6855 | 3244/5654 | 1091/7818 | NA | 304/5961 |
| Middlesborough | 788/2175 | 504/2093 | 2934/8017 | 826/10477 | 5319/9595 | 1447/13520 | 1239/6727 | 650/9660 |
| Newcastle | 1165/3363 | 746/3246 | NA | 1355/18166 | 9120/16690 | 2836/23035 | NA | 908/17522 |
| Nottingham | 1208/3541 | 744/3420 | 676/1929 | 1325/17305 | 8178/15766 | 2515/21492 | 281/1639 | 636/15239 |
| Oxford | 455/1695 | 274/1639 | NA | 656/6793 | 3236/6577 | 1148/8690 | NA | 205/5427 |
| Reading | 999/4126 | 614/3995 | NA | 947/13561 | 6298/14662 | 1950/19033 | NA | 379/11477 |
| Sheffield | 1125/3218 | 731/3081 | 4370/11248 | 1664/16219 | 7656/13696 | 2070/19329 | 1763/9436 | 581/12965 |
| Stockport_pilot | 45/90 | 27/87 | NA | 33/247 | 108/207 | 30/287 | NA | 10/196 |
| Stoke | 485/1519 | 299/1474 | NA | 874/9823 | 4700/8913 | 1443/12210 | NA | 424/8216 |
| Swansea | 93/228 | 59/221 | 360/762 | 107/1147 | 609/964 | 188/1385 | 127/632 | 39/1048 |
| Wrexham | 33/52 | 23/48 | 106/255 | 27/333 | 169/307 | 52/427 | 31/222 | 21/304 |

**Supplemental Table S4: Number of samples by assessment centre**

This table lists the number of cases (N_cases) and controls (N_controls) collected for each of the definitions in UKBiobank in the categories CIDI-based, Symptom, Self-report, Help-seeking, and no-MDD collected in each assessment centre. "NA" denotes no collection of answers to questions that form diagnostic criteria to the relevant definition of MDD in an assessment centre.

| CATEGORY | DEFINITION | NSAMPLES | PREV | CORRECTED_PREV |
|---|---|---|---|---|
| CIDI-based | LifetimeMDD | 67171 | 0.243 | 0.268 |
| CIDI-based | MDDRecur | 59385 | 0.173 | 0.196 |
| Symptom | DepAll | 79576 | 0.266 | 0.279 |
| Self-report | SelfRepDep | 253926 | 0.078 | 0.084 |
| EMR | ICD10Dep | 212411 | 0.043 | 0.043 |
| Help-seeking | GPpsy | 332629 | 0.341 | 0.343 |
| Help-seeking | Psypsy | 333419 | 0.109 | 0.111 |

**Supplemental Table S5: Prevalence of each definition of MDD**
This table shows for each definition of MDD in UKBiobank the category we assign it to (CIDI-based, Symptom, Self-report, Help-seeking, or no-MDD), the number of samples (both cases and controls), the case prevalence (prev), and the case prevalence when corrected for age group, sex and population in each assessment centre where samples are collected. Correction factors for age, sex and population in each assessment centre are shown in Supplemental Tables S5,6.

| CENTRE | UKBIOBANK | | CENSUS_2011 | | CORRECTION |
|---|---|---|---|---|---|
| | N_SAMPLE | FRACTION | POPULATION | POP_FRACTION | |
| Cardiff | 12678 | 0.038 | 346090 | 0.044 | 1.151 |
| Sheffield | 21479 | 0.064 | 552698 | 0.070 | 1.085 |
| Hounslow | 14439 | 0.043 | 253957 | 0.032 | 0.742 |
| Leeds | 30964 | 0.093 | 751485 | 0.095 | 1.023 |
| Croydon | 14778 | 0.044 | 363378 | 0.046 | 1.037 |
| Birmingham | 15783 | 0.047 | 1073045 | 0.135 | 2.866 |
| Bristol | 30406 | 0.091 | 428234 | 0.054 | 0.594 |
| Nottingham | 24106 | 0.072 | 305680 | 0.039 | 0.535 |
| Reading | 21050 | 0.063 | 155698 | 0.020 | 0.312 |
| Liverpool | 22399 | 0.067 | 466415 | 0.059 | 0.878 |
| Newcastle | 25963 | 0.078 | 280177 | 0.035 | 0.455 |
| Stoke | 13705 | 0.041 | 249008 | 0.031 | 0.766 |
| Middlesborough | 15024 | 0.045 | 138412 | 0.017 | 0.388 |
| Manchester | 8949 | 0.027 | 503127 | 0.063 | 2.370 |
| Bury | 20138 | 0.060 | 185060 | 0.023 | 0.387 |
| Glasgow | 12589 | 0.038 | 593245 | 0.075 | 1.987 |
| Oxford | 9865 | 0.029 | 151906 | 0.019 | 0.649 |
| Edinburgh | 12224 | 0.037 | 476626 | 0.060 | 1.644 |
| Barts | 5779 | 0.017 | 7375 | 0.001 | 0.054 |
| Wrexham | 481 | 0.001 | 134844 | 0.017 | 11.819 |
| Stockport_pilot | 317 | 0.001 | 283275 | 0.036 | 37.675 |
| Swansea | 1583 | 0.005 | 239023 | 0.030 | 6.366 |

**Supplemental Table S6: Correction for population in assessment centres**

This stable shows the number of samples (N_sample) and the fraction of total sample size (fraction) from each assessment centre in UKBiobank, as well as the population in cities where each of the collection centres are located (population) and the fraction of the UK population they represent (pop_fraction) according to the UK Census 2011. Only cities where there are UKBiobank assessment centres are included in this table and considered when calculating fractions of the UK population. The correction factor (correction) is the ratio between the population fraction and sample fraction, and is multiplied to the case prevalence of each definition of MDD from each assessment centre when calculating the corrected prevalence for each definition of MDD across all assessment centres, along with correction factors for age and sex as shown in Supplemental Table S6.

| AGE | UKBIOBANK | | | CENSUS_2011 | | CORRECTION | |
|---|---|---|---|---|---|---|---|
| | N_SAMPLE | % AGE | % FEMALE | % AGE | % FEMALE | AGE | FEMALE |
| 39to44 | 31412 | 9.4 | 53.3 | 17.6 | 50.1 | 1.875 | 0.940 |
| 45to49 | 41684 | 12.5 | 55.3 | 17.7 | 50.3 | 1.421 | 0.909 |
| 50to54 | 49730 | 14.9 | 56.3 | 15.6 | 50.6 | 1.050 | 0.898 |
| 55to59 | 60517 | 18.1 | 55.4 | 13.8 | 50.6 | 0.763 | 0.913 |
| 60to64 | 84850 | 25.4 | 53.7 | 14.5 | 50.4 | 0.572 | 0.938 |
| above65 | 66506 | 19.9 | 49.6 | 26.0 | 50.8 | 1.308 | 1.023 |

**Supplemental Table S7: Correction for age and sex**

This stable shows the number of samples (N_sample) and the fraction of total sample size of each age group (% age) as well as fraction of females (% female) per age group in UKBiobank, as well as the fraction of the UK population from each age group (% age) and who are female (% female) per age group according to the UK Census 2011. Only age groups of those ages of samples included in UKBiobank are shown in this table and considered when calculating fractions of each age group and sex. Correction factors (age, sex) are the ratios between the population fraction and sample fraction of each age group and sex, and are multiplied by the case prevalence of each definition of MDD along with correction factors for assessment centre population as shown in Supplemental Table S5 when calculating the corrected prevalence for each definition of MDD.

| CHR | RSID | BP | A1 | A0 | A1FREQ | OR | SE | P | USED IN MR | REASON NOT USED |
|---|---|---|---|---|---|---|---|---|---|---|
| 1 | rs7542974 | 72,544,704 | A | G | 0.25 | 1.032 | 0.0053 | 3.80E-08 | YES | |
| 1 | rs485929 | 74,678,285 | G | A | 0.39 | 1.028 | 0.0048 | 3.70E-08 | YES | |
| 1 | rs532246 | 84,411,238 | G | A | 0.74 | 0.968 | 0.0051 | 7.00E-09 | YES | |
| 1 | rs2789111 | 243,346,404 | C | T | 0.38 | 0.968 | 0.0054 | 1.50E-10 | NO | INFO < 0.9 |
| 2 | rs35028061 | 49,479,987 | GT | G | 0.38 | 1.029 | 0.005 | 1.90E-08 | NO | INDEL |
| 3 | rs9917656 | 48,581,513 | C | T | 0.3 | 1.03 | 0.0056 | 3.20E-08 | YES | |
| 3 | rs13082026 | 52,962,681 | T | C | 0.44 | 0.972 | 0.005 | 2.40E-08 | NO | INFO < 0.9 |
| 4 | rs57692580 | 106,214,476 | A | T | 0.39 | 0.973 | 0.0046 | 2.80E-08 | NO | INFO < 0.9 |
| 5 | rs34635 | 60,513,501 | G | A | 0.42 | 0.972 | 0.0045 | 1.20E-08 | NO | INFO < 0.9 |
| 5 | rs146681214 | 133,867,867 | AC | A | 0.18 | 1.039 | 0.0065 | 3.60E-09 | NO | INDEL |
| 5 | rs2336897 | 167,050,276 | T | C | 0.69 | 1.031 | 0.0061 | 5.20E-09 | YES | |
| 6 | rs3993747 | 31,580,507 | G | A | 0.35 | 0.969 | 0.0044 | 9.50E-10 | YES | |
| 6 | rs59732267 | 98,432,302 | CA | C | 0.52 | 0.972 | 0.0047 | 2.50E-08 | NO | INDEL |
| 8 | rs28716319 | 83,269,854 | G | A | 0.28 | 1.031 | 0.0057 | 2.70E-08 | YES | |
| 8 | rs13262595 | 143,316,970 | G | A | 0.56 | 1.03 | 0.005 | 1.00E-09 | YES | |
| 9 | rs6474966 | 15,757,537 | A | G | 0.46 | 1.028 | 0.0049 | 2.80E-08 | YES | |
| 9 | rs11793831 | 23,362,311 | T | G | 0.42 | 1.027 | 0.0053 | 4.30E-08 | YES | |
| 11 | rs1984389 | 31,740,989 | C | A | 0.54 | 0.973 | 0.0046 | 2.40E-08 | YES | |
| 11 | rs10791143 | 131,278,676 | G | A | 0.62 | 1.034 | 0.0046 | 1.50E-11 | YES | |
| 16 | rs4616299 | 7,657,432 | G | A | 0.4 | 0.972 | 0.005 | 1.20E-08 | YES | |
| 17 | rs56058331 | 56,427,128 | A | G | 0.42 | 1.029 | 0.0047 | 1.00E-08 | NO | INFO < 0.9 |
| 18 | rs1261078 | 52,866,791 | G | A | 0.05 | 0.927 | 0.0107 | 5.60E-12 | YES | |
| 19 | rs34232444 | 4,965,404 | C | T | 0.35 | 1.029 | 0.0057 | 2.50E-08 | NO | INFO < 0.9 |
| 19 | rs3746187 | 18,279,816 | G | A | 0.4 | 0.968 | 0.0049 | 9.80E-11 | YES | |
| 19 | rs429358 | 45,411,941 | C | T | 0.15 | 0.942 | 0.0067 | 4.60E-19 | YES | |

**Supplemental Table S8: GWAS hits for participation in the MHQ (adapted from Adams et al 2019)**

This table shows the genome-wide significant loci (top SNP in 1MB regions) in GWAS for participation in the MHQ, as reported in Adams et al 2019 [6]. For each locus, we show the chromosome (CHR), rsid for independent significant variants (RSID), position on the chromosome (BP), test allele (A1), other allele (A0), allele frequency of the test allele (A1FREQ) in all White-British samples in UKBiobank, odds ratio of the test allele on the phenotype (OR), standard error of the OR (SE), p-value of the association (P), and whether the locus is used in the Mendelian Randomisation analysis with definitions of depression in UKBiobank (USED IN MR), and reason if it is not used (REASON NOT USED).

| CATEGORY | DEFINITION | METHOD | All 16 MHQ GWAS SNPs | | | | REMOVE 2 OUTLIERS | | | |
|---|---|---|---|---|---|---|---|---|---|---|
| | | | ESTIMATE | SE | 95% CI | P | ESTIMATE | SE | 95% CI | P |
| CIDI-based | LifetimeMDD | Simple Median | 0.035 | 0.147 | [-0.252,0.323] | 0.810 | 0.035 | 0.165 | [-0.288,0.359] | 0.831 |
| | | Weighted Median | 0.033 | 0.141 | [-0.244,0.310] | 0.816 | 0.035 | 0.162 | [-0.282,0.351] | 0.829 |
| | | IVW | 0.062 | 0.144 | [-0.220,0.343] | 0.668 | 0.094 | 0.168 | [-0.234,0.423] | 0.574 |
| | | MR-Egger | -0.229 | 0.606 | [-1.417,0.960] | 0.706 | -0.910 | 2.464 | [-5.739,3.919] | 0.712 |
| | | MR-Egger intercept | 0.010 | 0.020 | [-0.029,0.048] | 0.622 | 0.030 | 0.073 | [-0.114,0.174] | 0.683 |
| | MDDRecur | Simple Median | -0.033 | 0.175 | [-0.376,0.310] | 0.851 | -0.033 | 0.196 | [-0.417,0.352] | 0.867 |
| | | Weighted Median | -0.036 | 0.172 | [-0.373,0.300] | 0.833 | -0.033 | 0.193 | [-0.411,0.345] | 0.864 |
| | | IVW | -0.007 | 0.156 | [-0.313,0.298] | 0.963 | -0.032 | 0.182 | [-0.389,0.326] | 0.862 |
| | | MR-Egger | 0.069 | 0.658 | [-1.220,1.358] | 0.917 | -2.290 | 2.624 | [-7.432,2.853] | 0.383 |
| | | MR-Egger intercept | -0.003 | 0.021 | [-0.044,0.039] | 0.905 | 0.067 | 0.078 | [-0.086,0.221] | 0.388 |
| Symptom | DepAll | Simple Median | -0.061 | 0.136 | [-0.327,0.205] | 0.651 | -0.061 | 0.157 | [-0.369,0.246] | 0.695 |
| | | Weighted Median | -0.010 | 0.132 | [-0.269,0.248] | 0.937 | -0.065 | 0.152 | [-0.363,0.234] | 0.672 |
| | | IVW | 0.001 | 0.113 | [-0.220,0.222] | 0.993 | -0.008 | 0.134 | [-0.270,0.255] | 0.954 |
| | | MR-Egger | -0.048 | 0.471 | [-0.971,0.875] | 0.919 | -1.363 | 1.947 | [-5.180,2.454] | 0.484 |
| | | MR-Egger intercept | 0.002 | 0.015 | [-0.028,0.032] | 0.915 | 0.041 | 0.058 | [-0.073,0.154] | 0.485 |
| Self-report | SelfRepDep | Simple Median | -0.278 | 0.127 | [-0.527,-0.029] | 0.028 | -0.278 | 0.150 | [-0.573,0.017] | 0.065 |
| | | Weighted Median | -0.276 | 0.124 | [-0.519,-0.033] | 0.026 | -0.190 | 0.142 | [-0.468,0.088] | 0.180 |
| | | IVW | -0.226 | 0.096 | [-0.414,-0.037] | 0.019 | -0.202 | 0.114 | [-0.425,0.021] | 0.075 |
| | | MR-Egger | -0.410 | 0.401 | [-1.196, 0.376] | 0.307 | -0.480 | 1.665 | [-3.743,2.783] | 0.773 |
| | | MR-Egger intercept | 0.006 | 0.013 | [-0.019, 0.032] | 0.636 | 0.008 | 0.050 | [-0.089,0.106] | 0.867 |
| EMR | ICD10Dep | Simple Median | -0.331 | 0.169 | [-0.662,-0.001] | 0.050 | -0.427 | 0.190 | [-0.800,-0.055] | 0.025 |
| | | Weighted Median | -0.307 | 0.166 | [-0.633, 0.018] | 0.064 | -0.405 | 0.187 | [-0.771,-0.039] | 0.030 |
| | | IVW | -0.335 | 0.120 | [-0.571,-0.099] | 0.005 | -0.369 | 0.135 | [-0.633,-0.105] | 0.006 |
| | | MR-Egger | 0.033 | 0.485 | [-0.919, 0.984] | 0.947 | 0.922 | 1.916 | [-2.832,4.677] | 0.630 |
| | | MR-Egger intercept | -0.012 | 0.016 | [-0.043, 0.019] | 0.434 | -0.039 | 0.057 | [-0.151,0.073] | 0.499 |
| Help-seeking | GPpsy | Simple Median | -0.033 | 0.067 | [-0.165,0.098] | 0.619 | 0.000 | 0.081 | [-0.159,0.159] | 1.000 |

| | | | | | | | | | | |
|---|---|---|---|---|---|---|---|---|---|---|
| | | Weighted Median | -0.038 | 0.065 | [-0.166,0.090] | 0.563 | -0.006 | 0.078 | [-0.159,0.147] | 0.936 |
| | | IVW | -0.075 | 0.078 | [-0.227,0.078] | 0.336 | -0.074 | 0.093 | [-0.255,0.108] | 0.426 |
| | | MR-Egger | -0.097 | 0.327 | [-0.738,0.544] | 0.766 | -0.114 | 1.368 | [-2.796,2.568] | 0.934 |
| | | MR-Egger intercept | 0.001 | 0.011 | [-0.020,0.022] | 0.944 | 0.001 | 0.041 | [-0.079,0.081] | 0.977 |
| | Psypsy | Simple Median | -0.327 | 0.108 | [-0.539,-0.115] | 0.003 | -0.371 | 0.126 | [-0.618,-0.124] | 0.003 |
| | | Weighted Median | -0.209 | 0.102 | [-0.409,-0.008] | 0.042 | -0.364 | 0.122 | [-0.604,-0.125] | 0.003 |
| | | IVW | -0.189 | 0.098 | [-0.380, 0.003] | 0.053 | -0.213 | 0.116 | [-0.440,0.013] | 0.065 |
| | | MR-Egger | -0.105 | 0.408 | [-0.904, 0.694] | 0.797 | -1.568 | 1.665 | [-4.832,1.696] | 0.346 |
| | | MR-Egger intercept | -0.003 | 0.013 | [-0.029, 0.023] | 0.833 | 0.040 | 0.050 | [-0.057,0.138] | 0.415 |
| Non-MDD | GPNoDep | Simple Median | -0.094 | 0.200 | [-0.487,0.299] | 0.638 | -0.032 | 0.233 | [-0.489,0.425] | 0.892 |
| | | Weighted Median | -0.102 | 0.195 | [-0.483,0.280] | 0.602 | -0.012 | 0.221 | [-0.446,0.422] | 0.958 |
| | | IVW | -0.062 | 0.160 | [-0.376,0.252] | 0.698 | -0.012 | 0.188 | [-0.381,0.358] | 0.951 |
| | | MR-Egger | -0.259 | 0.670 | [-1.572,1.054] | 0.699 | 2.350 | 2.695 | [-2.932,7.632] | 0.383 |
| | | MR-Egger intercept | 0.007 | 0.022 | [-0.036,0.049] | 0.762 | -0.071 | 0.080 | [-0.228,0.087] | 0.380 |

**Supplemental Table S9: Assessing casusal relationship between MHQ participation and definitions of depression**

This table shows the results from Mendelian Randomization (MR) assessment of SNPs significantly associated with MHQ participation for their effects on definitions of depression in UKBiobank, using the following estimators: inverse variance weighted (IVW), MR-Egger, Simple median, and Weighted median. The analysis is performed using both all 16 MHQ associated SNPs as shown in Supplemental Table S8, or removing two SNPs rs429358 and rs1261078 which have large effects on MHQ participation.

| CATEGORY | PHENO | NSAMPLES | CHR | SNP | BP | A1 | A0 | A1FREQ | OR | SE | P | MHC/SV |
|---|---|---|---|---|---|---|---|---|---|---|---|---|
| DSM-MDD | LifetimeMDD | 67171 | 6 | rs926552 | 29548089 | A | G | 0.140 | 0.903 | 0.019 | 4.46E-08 | MHC |
| Help-seeking | GPpsy | 323344 | 1 | rs6699744 | 72825144 | A | T | 0.388 | 0.960 | 0.005 | 6.55E-14 | |
| Help-seeking | GPpsy | 332112 | 1 | rs6697602 | 177039372 | G | C | 0.083 | 1.056 | 0.009 | 6.36E-09 | |
| Help-seeking | GPpsy | 331205 | 2 | rs11123030 | 124976163 | T | C | 0.491 | 1.032 | 0.005 | 1.13E-09 | |
| Help-seeking | GPpsy | 326346 | 3 | rs66511648 | 117515519 | C | T | 0.284 | 1.033 | 0.006 | 2.42E-08 | |
| Help-seeking | GPpsy | 329391 | 5 | rs30266 | 103972357 | A | G | 0.328 | 1.039 | 0.006 | 3.50E-12 | |
| Help-seeking | GPpsy | 331857 | 6 | rs12205083 | 24275483 | G | A | 0.104 | 1.053 | 0.008 | 6.24E-10 | |
| Help-seeking | GPpsy | 332546 | 6 | rs75782365 | 26408551 | G | T | 0.110 | 0.951 | 0.008 | 1.36E-09 | MHC |
| Help-seeking | GPpsy | 332629 | 6 | rs7772160 | 27412386 | C | T | 0.478 | 0.965 | 0.005 | 3.27E-12 | MHC |
| Help-seeking | GPpsy | 332629 | 6 | rs3135296 | 28795856 | T | A | 0.120 | 0.946 | 0.008 | 3.75E-12 | MHC |
| Help-seeking | GPpsy | 326902 | 6 | rs3115631 | 29986324 | A | T | 0.126 | 0.944 | 0.008 | 2.86E-13 | MHC |
| Help-seeking | GPpsy | 332629 | 6 | rs1625792 | 31306420 | A | G | 0.147 | 0.961 | 0.007 | 4.59E-08 | MHC |
| Help-seeking | GPpsy | 327948 | 6 | rs236346 | 36832103 | C | T | 0.095 | 0.950 | 0.009 | 1.19E-08 | MHC |
| Help-seeking | GPpsy | 331741 | 6 | rs9345737 | 66676938 | G | A | 0.439 | 0.969 | 0.005 | 3.03E-09 | |
| Help-seeking | GPpsy | 332629 | 7 | rs3807866 | 12250378 | A | G | 0.410 | 1.039 | 0.005 | 5.44E-13 | |
| Help-seeking | GPpsy | 327847 | 9 | rs393488 | 17044971 | A | T | 0.469 | 0.967 | 0.005 | 2.14E-10 | |
| Help-seeking | GPpsy | 331280 | 9 | rs12057031 | 25235063 | T | C | 0.108 | 0.952 | 0.008 | 5.29E-09 | |
| Help-seeking | GPpsy | 322805 | 10 | rs11599236 | 106454672 | C | T | 0.408 | 0.971 | 0.005 | 2.91E-08 | |
| Help-seeking | GPpsy | 332122 | 11 | rs537635 | 88705235 | T | C | 0.484 | 1.033 | 0.005 | 6.24E-10 | SV |
| Help-seeking | GPpsy | 329151 | 11 | rs578174 | 89959637 | G | A | 0.102 | 0.953 | 0.009 | 2.10E-08 | SV |

| | | | | | | | | | | | | |
|---|---|---|---|---|---|---|---|---|---|---|---|---|
| Help-seeking | GPpsy | 332489 | 14 | rs12889665 | 75234830 | T | G | 0.462 | 0.972 | 0.005 | 3.16E-08 | |
| Help-seeking | GPpsy | 329763 | 14 | rs61997596 | 104511206 | A | G | 0.191 | 1.037 | 0.007 | 4.15E-08 | |
| Help-seeking | GPpsy | 332629 | 16 | rs11646401 | 21609978 | G | C | 0.442 | 1.029 | 0.005 | 3.58E-08 | |
| Help-seeking | GPpsy | 327186 | 18 | rs12967855 | 35138245 | A | G | 0.329 | 1.034 | 0.006 | 1.57E-09 | |
| Help-seeking | GPpsy | 331002 | 18 | rs8097498 | 53449667 | G | A | 0.387 | 1.031 | 0.005 | 9.58E-09 | |
| Help-seeking | Psypsy | 332603 | 6 | rs66975207 | 26942146 | C | A | 0.111 | 0.925 | 0.013 | 1.23E-09 | MHC |
| Help-seeking | Psypsy | 333419 | 6 | rs4713145 | 28106827 | C | T | 0.244 | 0.941 | 0.009 | 5.32E-11 | MHC |
| Help-seeking | Psypsy | 333295 | 6 | rs3129120 | 29111775 | C | T | 0.124 | 0.925 | 0.012 | 1.45E-10 | MHC |
| Help-seeking | Psypsy | 333387 | 6 | rs2517622 | 30155149 | C | G | 0.137 | 0.926 | 0.012 | 4.20E-11 | MHC |
| Help-seeking | Psypsy | 319776 | 6 | rs535777 | 32577633 | C | G | 0.138 | 0.930 | 0.012 | 9.94E-10 | MHC |
| No-MDD | GPNoDep | 57572 | 6 | rs3094146 | 29970960 | C | G | 0.130 | 0.870 | 0.025 | 4.74E-08 | MHC |

**Supplemental Table S10: Genome-wide significant loci in GWAS for definitions of MDD**

This table shows the genome-wide significant loci (top SNP in 1MB regions) in GWAS for all definitions of MDD in UKBiobank. Only four definitions show genome-wide significant hits: CIDI-based LifetimeMDD, minimal phenotyping, help-seeking based definitions GPpsy and Psypsy, and minimal phenotyping, help-seeking based no-MDD definition that exclude MDD symptoms GPNoDep. For each locus we show the chromosome (CHR), rsid for the top SNP in 1MB window (SNP), position on the chromosome (BP), test and minor allele (A1), major allele (A0), allele frequency of the test allele (A1FREQ) in all White-British samples in UKBiobank, number of samples included in the linear regression at the locus with no missing genotypes, phenotype or covariate data (NMISS), standardized effect size of the minor allele on the phenotype (BETA), standard error of the effect (SE), p-value of the association (P), and whether the locus is in the MHC region on chr6:25-35MB or in any SV regions as listed in Price et al 2008[9] (MHC/SV). We note that rs3094146 in GPNoDep lies in the same locus as rs3115631 in GPpsy, and they lie in the same MB region as rs926552 in LifetimeMDD and rs3129120 in Psypsy, with GPNoDep showing the greatest size of effect (OR=0.84, SE=0.025) demonstrating this locus is not specific to MDD but shared and potentially driven by conditions other than MDD captured by GPNoDep.

| CATEGORY | PHENO | NSAMPLES | CHR | SNP | BP | A1 | A0 | A1FREQ | OR | SE | P | MHC/SV |
|---|---|---|---|---|---|---|---|---|---|---|---|---|
| DSM-MDD | MDD2 | 63196 | 6 | rs1233396 | 29546799 | A | G | 0.140 | 0.902 | 0.019 | 4.79E-08 | MHC |
| DSM-MDD | MDD2recur | 55054 | 5 | rs1833718 | 164635015 | T | C | 0.364 | 0.914 | 0.016 | 4.01E-08 | |
| Help-seeking | GPpsy | 318392 | 1 | rs6699744 | 72825144 | A | T | 0.388 | 0.960 | 0.005 | 5.98E-14 | |
| Help-seeking | GPpsy | 327022 | 1 | rs6697602 | 177039372 | G | C | 0.083 | 1.056 | 0.009 | 5.34E-09 | |
| Help-seeking | GPpsy | 326131 | 2 | rs11123030 | 124976163 | T | C | 0.491 | 1.032 | 0.005 | 9.95E-10 | |
| Help-seeking | GPpsy | 321342 | 3 | rs66511648 | 117515519 | C | T | 0.284 | 1.033 | 0.006 | 2.89E-08 | |
| Help-seeking | GPpsy | 322712 | 5 | rs40465 | 103981726 | G | T | 0.331 | 1.040 | 0.006 | 2.34E-12 | |
| Help-seeking | GPpsy | 321888 | 6 | rs3115631 | 29986324 | A | T | 0.126 | 0.942 | 0.008 | 5.95E-14 | MHC |
| Help-seeking | GPpsy | 327532 | 6 | rs2232423 | 28366151 | G | A | 0.117 | 0.944 | 0.008 | 1.73E-12 | MHC |
| Help-seeking | GPpsy | 327281 | 6 | rs67859638 | 27357978 | G | A | 0.113 | 0.948 | 0.008 | 1.08E-10 | MHC |
| Help-seeking | GPpsy | 326771 | 6 | rs12205083 | 24275483 | G | A | 0.104 | 1.053 | 0.008 | 1.39E-09 | |
| Help-seeking | GPpsy | 326658 | 6 | rs9345737 | 66676938 | G | A | 0.439 | 0.969 | 0.005 | 2.97E-09 | MHC |
| Help-seeking | GPpsy | 322923 | 6 | rs236346 | 36832103 | C | T | 0.095 | 0.950 | 0.009 | 1.13E-08 | MHC |
| Help-seeking | GPpsy | 323806 | 6 | rs34158769 | 26336572 | A | G | 0.106 | 0.954 | 0.009 | 2.36E-08 | MHC |
| Help-seeking | GPpsy | 327532 | 6 | rs1625792 | 31306420 | A | G | 0.147 | 0.960 | 0.007 | 3.51E-08 | MHC |
| Help-seeking | GPpsy | 327532 | 7 | rs3807866 | 12250378 | A | G | 0.410 | 1.039 | 0.005 | 5.11E-13 | |
| Help-seeking | GPpsy | 322818 | 9 | rs393488 | 17044971 | A | T | 0.469 | 0.968 | 0.005 | 7.69E-10 | |
| Help-seeking | GPpsy | 326201 | 9 | rs12057031 | 25235063 | T | C | 0.108 | 0.953 | 0.008 | 9.98E-09 | |
| Help-seeking | GPpsy | 317841 | 10 | rs11599236 | 106454672 | C | T | 0.408 | 0.970 | 0.005 | 2.04E-08 | |
| Help-seeking | GPpsy | 321170 | 11 | rs10765180 | 88740827 | T | G | 0.482 | 1.034 | 0.005 | 2.35E-10 | SV |
| Help-seeking | GPpsy | 324121 | 11 | rs578174 | 89959637 | G | A | 0.102 | 0.953 | 0.009 | 2.22E-08 | SV |
| Help-seeking | GPpsy | 326354 | 11 | rs4244537 | 28617622 | A | C | 0.379 | 0.971 | 0.005 | 4.97E-08 | SV |
| Help-seeking | GPpsy | 327394 | 14 | rs12889665 | 75234830 | T | G | 0.462 | 0.972 | 0.005 | 2.89E-08 | |
| Help-seeking | GPpsy | 324700 | 14 | rs61997596 | 104511206 | A | G | 0.191 | 1.037 | 0.007 | 3.64E-08 | |
| Help-seeking | GPpsy | 327532 | 16 | rs11646401 | 21609978 | G | C | 0.442 | 1.030 | 0.005 | 2.12E-08 | |
| Help-seeking | GPpsy | 322176 | 18 | rs12967855 | 35138245 | A | G | 0.329 | 1.034 | 0.006 | 1.64E-09 | |
| Help-seeking | GPpsy | 325934 | 18 | rs8097498 | 53449667 | G | A | 0.387 | 1.031 | 0.005 | 1.05E-08 | |
| Help-seeking | Psypsy | 281202 | 5 | rs542852 | 78409396 | T | C | 0.362 | 1.047 | 0.008 | 2.82E-08 | |

| | | | | | | | | | | | | |
|---|---|---|---|---|---|---|---|---|---|---|---|---|
| Help-seeking | Psypsy | 284937 | 6 | rs1235162 | 29537224 | G | A | 0.131 | 0.922 | 0.012 | 1.76E-11 | MHC |
| Help-seeking | Psypsy | 284937 | 6 | rs4713145 | 28106827 | C | T | 0.244 | 0.941 | 0.009 | 6.54E-11 | MHC |
| Help-seeking | Psypsy | 273221 | 6 | rs535777 | 32577633 | C | G | 0.138 | 0.929 | 0.012 | 6.38E-10 | MHC |
| Help-seeking | Psypsy | 284244 | 6 | rs66975207 | 26942146 | C | A | 0.111 | 0.924 | 0.013 | 8.87E-10 | MHC |
| Help-seeking | Psypsy | 284760 | 6 | rs3095327 | 30699022 | A | G | 0.159 | 0.935 | 0.011 | 1.07E-09 | MHC |
| Help-seeking | Psypsy | 282552 | 7 | rs6460894 | 12247330 | C | T | 0.338 | 1.048 | 0.008 | 1.80E-08 | |
| Help-seeking | Psypsy | 284120 | 14 | rs10144051 | 103885931 | A | C | 0.287 | 0.952 | 0.009 | 3.07E-08 | |
| No-MDD | GPNoDep | 56766 | 6 | rs3094146 | 29970960 | C | G | 0.130 | 0.869 | 0.025 | 3.53E-08 | MHC |

**Supplemental Table S11: Genome-wide significant loci in GWAS for definitions of MDD using "clean" controls**

This table shows the genome-wide significant loci (top SNP in 1MB regions) in GWAS for all definitions of MDD in UKBiobank, using "clean" controls rather than all controls. There are more significant loci for GPpsy and Psypsy, and one significant locus for MDDrecur that was not observed in GWAS using all controls at rs1833718. For each locus we show the chromosome (CHR), rsid for the top SNP in 1MB window (SNP), position on the chromosome (BP), test and minor allele (A1), major allele (A0), allele frequency of the test allele (A1FREQ) in all White-British samples in UKBiobank, number of samples included in the linear regression at the locus with no missing genotype, phenotype or covariate data (NMISS), standardized effect size of the minor allele on the phenotype (BETA), standard error of the effect (SE), p-value of the association (P), and whether the locus is in the MHC region on chr6:25-35MB or in any SV regions as listed in Price et al 2008[9] (MHC/SV).

| DATAFIELD | QUESTION ON CHILDHOOD TRAUMA "When I was growing up…" | RESPONSE AND SCORE (0/1) | | | | | | OR for LifetimeMDD |
|---|---|---|---|---|---|---|---|---|
| | | Prefer not to answer | Never true | Rarely true | Sometimes true | Often true | Very often true | |
| 20487 | I felt that someone in my family hated me | NA | 0 | 0 | 0 | 1 | 1 | 1.799 |
| 20488 | People in my family hit me so hard that it left me with bruises or marks | NA | 0 | 0 | 1 | 1 | 1 | 1.215 |
| 20489 | I felt loved | NA | 1 | 1 | 1 | 0 | 0 | 1.664 |
| 20490 | Someone molested me (sexually) | NA | 0 | 1 | 1 | 1 | 1 | 1.174 |
| 20491 | There was someone to take me to the doctor if I needed it | Question omitted | | | | | | |
| **DATAFIELD** | **QUESTION ON ADULTHOOD TRAUMA "Since I was sixteen…"** | **RESPONSE AND SCORE (0/1)** | | | | | | **OR for LifetimeMDD** |
| | | **Prefer not to answer** | **Never true** | **Rarely true** | **Sometimes true** | **Often true** | **Very often true** | |
| 20521 | A partner or ex-partner repeatedly belittled me to the extent that I felt worthless | NA | 0 | 0 | 1 | 1 | 1 | 2.388 |
| 20522 | I have been in a confiding relationship | NA | 1 | 1 | 0 | 0 | 0 | 1.039 |
| 20525 | There was money to pay the rent or mortgage when I needed it | NA | 1 | 1 | 1 | 0 | 0 | 1.311 |
| 20523 | A partner or ex-partner deliberately hit me or used violence in any other way | Question omitted | | | | | | |
| 20524 | A partner or ex-partner sexually interfered with me, or forced me to have sex against my wishes | Question omitted | | | | | | |

| DATAFIELD | QUESTION FOR LIFETIME TRAUMA "In your life, have you...?" | RESPONSE AND SCORE (0/1) | | | | OR for LifetimeMDD |
|---|---|---|---|---|---|---|
| | | Prefer not to answer | Never | Yes, but not in the last 12 months | Yes, within the last 12 months | |
| 20526 | Been in a serious accident that you believed to be life-threatening at the time | NA | 0 | 1 | 1 | 1.326 |
| 20527 | Been involved in combat or exposed to a war-zone (either in the military or as a civilian) | Question omitted | | | | |
| 20528 | Been diagnosed with a life-threatening illness | NA | 0 | 1 | 1 | 1.275 |
| 20529 | Been attacked, mugged, robbed, or been the victim of a physically violent crime | NA | 0 | 1 | 1 | 1.145 |
| 20530 | Witnessed a sudden violent death (eg. murder, suicide, aftermath of an accident) | NA | 0 | 1 | 1 | 1.288 |
| 20531 | Been a victim of a sexual assault, whether by a stranger or someone you knew | NA | 0 | 1 | 1 | 1.339 |

**Supplemental Table S12: Derivation of lifetime trauma score**

This table lists questions in the online mental health follow up questionnaire in UKBiobank forming the derivation of lifetime trauma score used in this paper (Supplemental Methods), along with their data field in the UKBiobank data showcase. The questions come in three categories, “Questions on childhood trauma”, “Questions on adulthood trauma”, and “Question for lifetime trauma”. Answers to questions that are of high similarity with another question are excluded from the calculation of lifetime trauma score (Question omitted). We show the multiple-choice answer to each of the questions which indicates a subject as self-reporting having experienced the traumatic life event. We also show the odds ratio (OR) of having experienced each of the traumatic life event on CIDI-based LifetimeMDD, when modelled jointly with all other traumatic life events, as well as age, sex, years of education, neuroticism, Townsend deprivation index, experience of any traumatic life event in the past 2 years (data field 6145), and total number of traumatic life events reported. We then weight each of the traumatic life events by its OR and sum all weighted scores to arrive at a weighted lifetime trauma score for each individual.

| PHENOTYPE | | | | | | PCGCs | | | | LDSC | | BOLT-REML | |
|---|---|---|---|---|---|---|---|---|---|---|---|---|---|
| CATEGORY | DEFINITION | OBS PREV | POP PREV | POPPREV ORIGIN | N | H2SNP | SE | COND H2SNP | SE | H2SNP | SE | H2SNP | SE |
| | | | 0.243 | observed prev | | 0.263 | 0.022 | 0.262 | 0.022 | 0.189 | 0.028 | 0.184 | 0.012 |
| | | | 0.268 | census correction | | 0.270 | 0.023 | 0.269 | 0.023 | 0.194 | 0.029 | 0.189 | 0.013 |
| CIDI-based | LifetimeMDD | 0.243 | 0.150 | MDD prev = 0.15 | 67171 | 0.228 | 0.019 | 0.227 | 0.019 | 0.163 | 0.024 | 0.159 | 0.011 |
| | | | 0.173 | observed prev | | 0.321 | 0.026 | 0.320 | 0.026 | 0.228 | 0.035 | 0.218 | 0.016 |
| | | | 0.196 | census correction | | 0.332 | 0.026 | 0.331 | 0.026 | 0.237 | 0.036 | 0.226 | 0.017 |
| CIDI-based | MDDRecur | 0.173 | 0.150 | MDD prev = 0.15 | 59385 | 0.306 | 0.024 | 0.305 | 0.024 | 0.218 | 0.033 | 0.208 | 0.015 |
| | | | 0.266 | observed prev | | 0.185 | 0.015 | 0.185 | 0.015 | 0.131 | 0.018 | 0.132 | 0.010 |
| | | | 0.279 | census correction | | 0.188 | 0.015 | 0.187 | 0.015 | 0.132 | 0.019 | 0.134 | 0.010 |
| Symptom | DepAll | 0.266 | 0.150 | MDD prev = 0.15 | 79576 | 0.157 | 0.013 | 0.156 | 0.012 | 0.111 | 0.015 | 0.112 | 0.008 |
| | | | 0.078 | observed prev | | 0.110 | 0.009 | 0.110 | 0.009 | 0.079 | 0.010 | 0.077 | 0.006 |
| | | | 0.084 | census correction | | 0.112 | 0.009 | 0.112 | 0.009 | 0.081 | 0.011 | 0.079 | 0.006 |
| Self-report | SelfRepDep | 0.078 | 0.150 | MDD prev = 0.15 | 253926 | 0.135 | 0.010 | 0.135 | 0.010 | 0.097 | 0.013 | 0.095 | 0.007 |
| | | | 0.043 | observed prev | | 0.127 | 0.012 | 0.127 | 0.012 | 0.114 | 0.014 | 0.075 | 0.010 |
| | | | 0.043 | census correction | | 0.126 | 0.011 | 0.125 | 0.011 | 0.114 | 0.014 | 0.075 | 0.010 |
| EMR | ICD10Dep | 0.043 | 0.150 | MDD prev = 0.15 | 212411 | 0.185 | 0.017 | 0.184 | 0.017 | 0.164 | 0.021 | 0.111 | 0.014 |
| | | | 0.341 | observed prev | | 0.144 | 0.008 | 0.143 | 0.008 | 0.120 | 0.009 | 0.109 | 0.003 |
| | | | 0.343 | census correction | | 0.144 | 0.008 | 0.144 | 0.008 | 0.120 | 0.009 | 0.109 | 0.002 |
| Help-seeking | GPpsy | 0.341 | 0.150 | MDD prev = 0.15 | 332629 | 0.114 | 0.007 | 0.114 | 0.007 | 0.096 | 0.007 | 0.087 | 0.002 |
| | | | 0.109 | observed prev | | 0.126 | 0.012 | 0.128 | 0.012 | 0.103 | 0.014 | 0.090 | 0.004 |
| | | | 0.111 | census correction | | 0.127 | 0.012 | 0.126 | 0.012 | 0.103 | 0.014 | 0.090 | 0.004 |
| Help-seeking | Psypsy | 0.109 | 0.150 | MDD prev = 0.15 | 333419 | 0.139 | 0.014 | 0.139 | 0.014 | 0.113 | 0.015 | 0.099 | 0.004 |
| | | | 0.149 | observed prev | | 0.130 | 0.028 | 0.130 | 0.028 | 0.084 | 0.038 | 0.052 | 0.015 |
| | | | 0.146 | census correction | | 0.129 | 0.031 | 0.129 | 0.031 | 0.083 | 0.038 | 0.052 | 0.015 |
| Non-MDD | GPNoDep | 0.149 | 0.150 | MDD prev = 0.15 | 58125 | 0.130 | 0.031 | 0.130 | 0.031 | 0.084 | 0.038 | 0.052 | 0.015 |

**Supplemental Table S13: Heritability estimates of definitions of MDD from different methods**

This table shows the $h^2_{SNP}$ estimates of each definition of MDD calculated using three different methods, PCGCs, LDSC, and BOLD-REML, as detailed in Supplemental Methods. For each definition of MDD we show the sample size (N), observed prevalence (PREV), assumed population prevalence (POP PREV), the origin of the assumed population prevalence (POP PREV ORIGIN). We show results from three different assumed population prevalences: the observed prevalence, the census corrected observed prevalence, and previously reported population prevalence of 0.15[8,32]. For each method we show the $h^2_{SNP}$ estimate (H2SNP) and its standard error (SE). For PCGCs, we also show the conditional $h^2_{SNP}$ (COND H2) and its standard error (SE) for completeness [20].

| RISK FACTOR | PHENOTYPE | | N (PREV) | H2SNP (SE) | N (PREV) | H2SNP (SE) | DIFFERENCE | | | |
|---|---|---|---|---|---|---|---|---|---|---|
| | CATEGORY | DEFINITION | AGE >= 60 | | AGE < 60 | | ΔH2SNP | DF | T-TEST STAT | T-TEST PVALUE |
| AGE | CIDI-based | LifetimeMDD | 27671 (0.166) | 0.298 (0.039) | 39500 (0.297) | 0.312 (0.027) | 0.015 | 52168 | -0.310 | 3.78E-01 |
| | CIDI-based | MDDRecur | 24894 (0.106) | 0.256 (0.048) | 34491 (0.222) | 0.387 (0.038) | 0.132 | 51023 | -2.142 | 1.61E-02 |
| | Symptom | DepAll | 38683 (0.224) | 0.215 (0.027) | 40893 (0.306) | 0.232 (0.022) | 0.017 | 76179 | -0.503 | 3.07E-01 |
| | Self-report | SelfRepDep | 126331 (0.058) | 0.141 (0.016) | 127595 (0.098) | 0.129 (0.013) | -0.012 | 243413 | 0.587 | 2.79E-01 |
| | EMR | ICD10Dep | 107665 (0.036) | 0.150 (0.025) | 104746 (0.050) | 0.170 (0.021) | 0.020 | 205648 | -0.604 | 2.73E-01 |
| | Help-seeking | GPpsy | 150368 (0.318) | 0.151 (0.014) | 182261 (0.359) | 0.153 (0.008) | 0.002 | 233073 | -0.143 | 4.43E-01 |
| | Help-seeking | Psypsy | 150840 (0.104) | 0.150 (0.015) | 182579 (0.113) | 0.124 (0.012) | -0.026 | 301479 | 1.303 | 9.63E-02 |
| | No-MDD | GPNoDep | 29866 (0.152) | 0.208 (0.044) | 28259 (0.145) | 0.153 (0.041) | -0.055 | 58049 | 0.920 | 1.79E-01 |
| | CATEGORY | DEFINITION | AGE >= 60 | | AGE < 60 | | ΔH2SNP | DF | T-TEST STAT | T-TEST PVALUE |
| DEPRIVATION | CIDI-based | LifetimeMDD | 13969 (0.304) | 0.418 (0.059) | 53120 (0.226) | 0.252 (0.025) | -0.166 | 19489 | 2.596 | 4.71E-03 |
| | CIDI-based | MDDRecur | 12308 (0.239) | 0.434 (0.073) | 47013 (0.156) | 0.338 (0.032) | -0.096 | 17400 | 1.212 | 1.13E-01 |
| | Symptom | DepAll | 20435 (0.309) | 0.325 (0.038) | 59023 (0.251) | 0.189 (0.018) | -0.136 | 30436 | 3.264 | 5.50E-04 |

| | | | | | | | | | | |
|---|---|---|---|---|---|---|---|---|---|---|
| | Self-report | SelfRepDep | 65877 (0.099) | 0.178 (0.023) | 187750 (0.071) | 0.119 (0.011) | -0.059 | 94849 | 2.297 | 1.08E-02 |
| | EMR | ICD10Dep | 56571 (0.063) | 0.216 (0.034) | 155596 (0.036) | 0.130 (0.017) | 0.086 | 88100 | 2.276 | 1.14E-02 |
| | Help-seeking | GPpsy | 83671 (0.390) | 0.149 (0.011) | 248566 (0.324) | 0.145 (0.008) | -0.004 | 182828 | 0.309 | 3.79E-01 |
| | Help-seeking | Psypsy | 83835 (0.149) | 0.154 (0.016) | 249190 (0.095) | 0.119 (0.011) | -0.034 | 172056 | 1.724 | 4.23E-02 |
| | No-MDD | GPNoDep | 14046 (0.168) | 0.311 (0.071) | 44004 (0.142) | 0.143 (0.029) | -0.168 | 18912 | 2.203 | 1.38E-02 |
| **TRAUMA** | **CATEGORY** | **DEFINITION** | **AGE >= 60** | | **AGE < 60** | | **ΔH2SNP** | **DF** | **T-TEST STAT** | **T-TEST PVALUE** |
| | CIDI-based | LifetimeMDD | 19110 (0.424) | 0.401 (0.047) | 48057 (0.171) | 0.232 (0.025) | -0.169 | 30581 | 3.185 | 7.24E-04 |
| | CIDI-based | MDDRecur | 16139 (0.353) | 0.468 (0.051) | 47013 (0.156) | 0.338 (0.032) | -0.130 | 35988 | 2.849 | 2.20E-03 |
| | Symptom | DepAll | 8448 (0.408) | 0.344 (0.076) | 20544 (0.219) | 0.295 (0.049) | -0.048 | 15766 | 0.535 | 2.96E-01 |
| | Self-report | SelfRepDep | 26454 (0.117) | 0.220 (0.047) | 54827 (0.057) | 0.252 (0.032) | 0.032 | 51237 | -0.550 | 2.91E-01 |
| | EMR | ICD10Dep | 21986 (0.049) | 0.321 (0.093) | 41337 (0.021) | 0.286 (0.095) | 0.035 | 58316 | 0.266 | 3.95E-01 |
| | Help-seeking | GPpsy | 33451 (0.447) | 0.233 (0.028) | 75790 (0.277) | 0.169 (0.015) | -0.064 | 53301 | 1.998 | 2.28E-02 |
| | Help-seeking | Psypsy | 33502 (0.160) | 0.203 (0.033) | 75929 (0.072) | 0.193 (0.031) | -0.011 | 89536 | 0.238 | 4.06E-01 |

| | CATEGORY | DEFINITION | AGE >= 60 | | AGE < 60 | | ΔH2SNP | DF | T-TEST STAT | T-TEST PVALUE |
|---|---|---|---|---|---|---|---|---|---|---|
| | No-MDD | GPNoDep | 4980 (0.186) | 0.759 (0.181) | 16007 (0.119) | 0.308 (0.065) | -0.452 | 6333 | 2.349 | 9.43E-03 |

| | CATEGORY | DEFINITION | AGE >= 60 | | AGE < 60 | | ΔH2SNP | DF | T-TEST STAT | T-TEST PVALUE |
|---|---|---|---|---|---|---|---|---|---|---|
| NEUROTICISM | CIDI-based | LifetimeMDD | 19043 (0.439) | 0.381 (0.043) | 38003 (0.136) | 0.289 (0.030) | -0.092 | 37284 | 1.746 | 4.04E-02 |
| | CIDI-based | MDDRecur | 16148 (0.370) | 0.451 (0.054) | 34367 (0.072) | 0.315 (0.045) | -0.136 | 37543 | 1.948 | 2.57E-02 |
| | Symptom | DepAll | 25691 (0.420) | 0.212 (0.030) | 41428 (0.171) | 0.183 (0.027) | -0.029 | 58976 | 0.711 | 2.39E-01 |
| | Self-report | SelfRepDep | 87606 (0.141) | 0.106 (0.013) | 117206 (0.027) | 0.194 (0.032) | 0.088 | 150998 | -2.567 | 5.12E-03 |
| | EMR | ICD10Dep | 73570 (0.074) | 0.156 (0.023) | 97407 (0.042) | 0.176 (0.046) | 0.020 | 141169 | -0.378 | 3.53E-01 |
| | Help-seeking | GPpsy | 110335 (0.513) | 0.129 (0.008) | 160959 (0.206) | 0.115 (0.010) | -0.014 | 270728 | 1.067 | 1.43E-01 |
| | Help-seeking | Psypsy | 110374 (0.183) | 0.118 (0.012) | 161225 (0.052) | 0.139 (0.015) | 0.021 | 271596 | -1.097 | 1.36E-01 |
| | No-MDD | GPNoDep | 14841 (0.257) | 0.228 (0.054) | 34261 (0.088) | 0.120 (0.046) | -0.108 | 35650 | 1.526 | 6.35E-02 |

| | CATEGORY | DEFINITION | AGE >= 60 | | AGE < 60 | | ΔH2SNP | DF | T-TEST STAT | T-TEST PVALUE |
|---|---|---|---|---|---|---|---|---|---|---|
| SEX | CIDI-based | LifetimeMDD | 32933 (0.167) | 0.294 (0.039) | 34238 (0.315) | 0.328 (0.025) | 0.033 | 57293 | -0.719 | 2.36E-01 |
| | CIDI-based | MDDRecur | 29760 (0.110) | 0.360 (0.047) | 29625 (0.237) | 0.429 (0.033) | 0.069 | 53839 | -1.204 | 1.14E-01 |
| | Symptom | DepAll | 37728 (0.203) | 0.203 (0.034) | 41848 (0.323) | 0.218 (0.021) | 0.015 | 64013 | -0.383 | 3.51E-01 |

| | CATEGORY | DEFINITION | AGE >= 60 | | AGE < 60 | | ΔH2SNP | DF | T-TEST STAT | T-TEST PVALUE |
|---|---|---|---|---|---|---|---|---|---|---|
| | Self-report | SelfRepDep | 117712 (0.058) | 0.133 (0.015) | 136214 (0.095) | 0.125 (0.012) | -0.008 | 229531 | 0.432 | 3.33E-01 |
| | EMR | ICD10Dep | 95851 (0.036) | 0.171 (0.025) | 116560 (0.049) | NA (NA) | NA | NA | NA | NA |
| | Help-seeking | GPpsy | 153917 (0.257) | 0.149 (0.011) | 178712 (0.413) | 0.170 (0.009) | 0.021 | 307791 | -1.512 | 6.52E-02 |
| | Help-seeking | Psypsy | 154190 (0.097) | 0.131 (0.016) | 179229 (0.119) | 0.145 (0.012) | 0.014 | 293736 | -0.699 | 2.42E-01 |
| | No-MDD | GPNoDep | 29968 (0.112) | 0.188 (0.041) | 28157 (0.187) | 0.228 (0.069) | 0.041 | 46298 | -0.507 | 3.06E-01 |

| | CATEGORY | DEFINITION | AGE >= 60 | | AGE < 60 | | ΔH2SNP | DF | T-TEST STAT | T-TEST PVALUE |
|---|---|---|---|---|---|---|---|---|---|---|
| **EDUCATION** | CIDI-based | LifetimeMDD | 17452 (0.244) | 0.356 (0.057) | 19815 (0.241) | 0.295 (0.042) | -0.061 | 6872 | -2.259 | 1.20E-02 |
| | CIDI-based | MDDRecur | 15446 (0.174) | 0.424 (0.071) | 17652 (0.174) | 0.298 (0.056) | -0.126 | 6152 | -1.607 | 5.40E-02 |
| | Symptom | DepAll | 19804 (0.261) | 0.306 (0.044) | 33136 (0.256) | 0.188 (0.027) | -0.118 | 19841 | -1.055 | 1.46E-01 |
| | Self-report | SelfRepDep | 59341 (0.078) | 0.132 (0.027) | 115285 (0.077) | 0.126 (0.014) | -0.006 | 70255 | -1.419 | 7.79E-02 |
| | EMR | ICD10Dep | 48482 (0.040) | 0.205 (0.046) | 101513 (0.050) | 0.145 (0.020) | 0.060 | 67284 | 1.207 | 1.14E-01 |
| | Help-seeking | GPpsy | 79630 (0.332) | 0.145 (0.013) | 144159 (0.353) | 0.144 (0.009) | -0.001 | 128193 | -0.890 | 1.87E-01 |
| | Help-seeking | Psypsy | 79791 (0.100) | 0.153 (0.024) | 144659 (0.114) | NA (NA) | NA | NA | NA | NA |

| | | | | | | | | | | |
|---|---|---|---|---|---|---|---|---|---|---|
| | No-MDD | GPNoDep | 14570 (0.140) | 0.372 (0.117) | 24492 (0.164) | 0.122 (0.043) | -0.251 | 14865 | -1.277 | 1.01E-01 |

**Supplemental Table S14: Heritability of definitions of MDD stratified by risk factors**

This table shows the $h^2_{SNP}$ estimates of each definition of MDD calculated using PCGCs in subgroups stratified by risk factors (age, deprivation, trauma, neuroticism, sex and education). For each stratum, we show the number of samples (N_sample), case prevalence (prev), $h^2_{SNP}$ estimate (H2SNP) and its standard error (SE). We show the difference between heritabilities from the two strata per risk factor (ΔH2SNP), and calculate the degree of freedom (DF), t-test statistic (T TEST STAT), and significance of this difference (T TEST PVALUE) using both $h^2_{SNP}$ estimates, standard errors and sample sizes. Items are "NA" when we were not able to get an estimate from PCGCs. None of the differences are significant in a two-tailed t-test at $p < 0{\cdot}05$ after multiple testing correction.

| Definitions | | PCGCs | | | | LDSC | | | | | | |
|---|---|---|---|---|---|---|---|---|---|---|---|---|
| **Definition1** | **Definition2** | **rG** | **rG SE** | **rho** | **rho SE** | **Intercept** | **rG** | **rG SE** | **rho** | **rho SE** | **Z Score** | **P value** |
| ICD10Dep | LifetimeMDD | 0.653 | 0.052 | 0.119 | 0.011 | 0.086 | 0.646 | 0.079 | 0.031 | 0.004 | 8.166 | 3.20E-16 |
| ICD10Dep | MDDRecur | 0.685 | 0.054 | 0.138 | 0.012 | 0.099 | 0.663 | 0.082 | 0.033 | 0.004 | 8.120 | 4.66E-16 |
| ICD10Dep | GPpsy | 0.762 | 0.036 | 0.103 | 0.006 | 0.190 | 0.798 | 0.055 | 0.033 | 0.003 | 14.442 | 2.82E-47 |
| ICD10Dep | Psypsy | 0.751 | 0.041 | 0.095 | 0.007 | 0.211 | 0.775 | 0.059 | 0.023 | 0.002 | 13.236 | 5.42E-40 |
| ICD10Dep | DepAll | 0.604 | 0.056 | 0.093 | 0.010 | 0.094 | 0.615 | 0.085 | 0.025 | 0.004 | 7.275 | 3.46E-13 |
| ICD10Dep | SelfRepDep | 0.778 | 0.050 | 0.092 | 0.007 | 0.265 | 0.893 | 0.079 | 0.021 | 0.002 | 11.265 | 1.94E-29 |
| ICD10Dep | GPNoDep | 0.539 | 0.088 | 0.069 | 0.011 | 0.028 | 0.662 | 0.183 | 0.019 | 0.004 | 3.624 | 3.00E-04 |
| LifetimeMDD | MDDRecur | 0.988 | 0.008 | 0.287 | 0.023 | 0.855 | 0.999 | 0.014 | 0.102 | 0.015 | 71.579 | 0 |
| LifetimeMDD | GPpsy | 0.811 | 0.026 | 0.158 | 0.012 | 0.259 | 0.903 | 0.044 | 0.077 | 0.008 | 20.333 | 6.55E-92 |
| LifetimeMDD | Psypsy | 0.772 | 0.035 | 0.141 | 0.015 | 0.179 | 0.883 | 0.054 | 0.054 | 0.008 | 16.254 | 2.10E-59 |
| LifetimeMDD | DepAll | 0.858 | 0.039 | 0.189 | 0.016 | 0.175 | 1.011 | 0.074 | 0.086 | 0.010 | 13.619 | 3.11E-42 |
| LifetimeMDD | SelfRepDep | 0.790 | 0.043 | 0.134 | 0.011 | 0.170 | 0.936 | 0.071 | 0.045 | 0.006 | 13.121 | 2.48E-39 |
| LifetimeMDD | GPNoDep | 0.577 | 0.078 | 0.107 | 0.021 | 0.042 | 0.725 | 0.142 | 0.043 | 0.013 | 5.108 | 3.26E-07 |
| MDDRecur | GPpsy | 0.852 | 0.029 | 0.183 | 0.013 | 0.245 | 0.956 | 0.047 | 0.083 | 0.009 | 20.516 | 1.55E-93 |
| MDDRecur | Psypsy | 0.799 | 0.040 | 0.160 | 0.017 | 0.186 | 0.909 | 0.061 | 0.057 | 0.009 | 14.951 | 1.52E-50 |
| MDDRecur | DepAll | 0.875 | 0.040 | 0.213 | 0.018 | 0.156 | 1.072 | 0.075 | 0.093 | 0.011 | 14.300 | 2.19E-46 |
| MDDRecur | SelfRepDep | 0.845 | 0.045 | 0.158 | 0.013 | 0.188 | 0.983 | 0.075 | 0.049 | 0.006 | 13.184 | 1.09E-39 |
| MDDRecur | GPNoDep | 0.619 | 0.081 | 0.126 | 0.024 | 0.038 | 0.809 | 0.156 | 0.049 | 0.015 | 5.202 | 1.97E-07 |
| GPpsy | Psypsy | 0.903 | 0.014 | 0.121 | 0.010 | 0.488 | 0.915 | 0.019 | 0.048 | 0.006 | 47.077 | 0 |
| GPpsy | DepAll | 0.853 | 0.027 | 0.139 | 0.010 | 0.410 | 0.918 | 0.046 | 0.067 | 0.007 | 20.070 | 1.34E-89 |
| GPpsy | SelfRepDep | 0.859 | 0.025 | 0.108 | 0.007 | 0.346 | 0.959 | 0.044 | 0.040 | 0.004 | 21.760 | 5.54E-105 |
| GPpsy | GPNoDep | 0.840 | 0.051 | 0.115 | 0.014 | 0.322 | 1.022 | 0.160 | 0.052 | 0.009 | 6.410 | 1.46E-10 |
| Psypsy | DepAll | 0.783 | 0.036 | 0.119 | 0.011 | 0.217 | 0.878 | 0.060 | 0.046 | 0.007 | 14.639 | 1.60E-48 |
| Psypsy | SelfRepDep | 0.830 | 0.029 | 0.098 | 0.008 | 0.326 | 0.950 | 0.049 | 0.028 | 0.003 | 19.546 | 4.50E-85 |
| Psypsy | GPNoDep | 0.697 | 0.069 | 0.089 | 0.017 | 0.092 | 0.882 | 0.124 | 0.032 | 0.009 | 7.138 | 9.50E-13 |
| DepAll | SelfRepDep | 0.790 | 0.044 | 0.113 | 0.009 | 0.151 | 1.010 | 0.085 | 0.041 | 0.005 | 11.953 | 6.27E-33 |
| DepAll | GPNoDep | 0.620 | 0.079 | 0.096 | 0.017 | 0.002 | 0.875 | 0.168 | 0.044 | 0.011 | 5.208 | 1.91E-07 |

| | | | | | | | | | | | | |
|---|---|---|---|---|---|---|---|---|---|---|---|---|
| SelfRepDep | GPNoDep | 0.587 | 0.080 | 0.070 | 0.013 | 0.043 | 0.793 | 0.139 | 0.023 | 0.006 | 5.703 | 1.18E-08 |

**Supplemental Table S15: Genetic correlation between definitions of depression in UKBiobank**

This table shows the genetic correlation between definitions of depression in UKBiobank from both PCGCs and LDSC. For each pair of definitions, we show the genetic correlation (rG), its standard error (rG SE), genetic covariance (rho) and its standard error (rho SE) for both methods. In addition, for LDSC, we show the intercept, Z score and P value of the estimation.

| CHR | SNP | BP | A1 | A0 | A1FREQ | NMISS | OR | SE | P | MHC/SV |
|---|---|---|---|---|---|---|---|---|---|---|
| 1 | rs1324481 | 33892964 | T | G | 0.317 | 333782 | 0.9708 | 0.005 | 4.01E-08 | |
| 1 | rs1697593 | 190453672 | T | C | 0.483 | 332979 | 1.028 | 0.005 | 3.70E-08 | |
| 1 | rs10863714 | 208710593 | A | G | 0.4238 | 335411 | 1.029 | 0.005 | 2.59E-08 | |
| 2 | rs1004787 | 45159091 | G | A | 0.4694 | 329067 | 0.9677 | 0.005 | 8.80E-11 | |
| 2 | rs4671381 | 60496910 | T | C | 0.4366 | 334992 | 0.9728 | 0.005 | 4.77E-08 | |
| 2 | rs67716713 | 104267572 | A | C | 0.4866 | 334796 | 0.9676 | 0.005 | 4.85E-11 | |
| 2 | rs1427499 | 145400317 | A | G | 0.289 | 336066 | 1.043 | 0.006 | 1.67E-14 | |
| 3 | rs11711099 | 25225163 | C | T | 0.4839 | 330036 | 1.029 | 0.005 | 1.75E-08 | |
| 3 | rs591988 | 61847695 | T | C | 0.2663 | 334385 | 1.032 | 0.006 | 4.25E-08 | |
| 3 | rs73121433 | 83154013 | A | G | 0.1632 | 328418 | 1.039 | 0.007 | 2.01E-08 | |
| 3 | rs12714592 | 84387950 | C | A | 0.2734 | 335552 | 1.038 | 0.006 | 4.85E-11 | SV |
| 3 | rs62250491 | 85616009 | G | T | 0.3713 | 333650 | 1.042 | 0.005 | 2.02E-15 | SV |
| 3 | rs1995245 | 117776020 | C | T | 0.1497 | 333233 | 0.959 | 0.007 | 2.45E-09 | |
| 4 | rs899632 | 57749347 | C | T | 0.3867 | 332773 | 0.9702 | 0.005 | 4.86E-09 | |
| 5 | rs1549212 | 166996722 | C | T | 0.3744 | 333822 | 1.032 | 0.005 | 2.22E-09 | |
| 6 | rs2892512 | 98744946 | C | T | 0.2684 | 334016 | 1.033 | 0.006 | 7.58E-09 | |
| 6 | rs465646 | 111620758 | G | A | 0.1588 | 334411 | 1.055 | 0.007 | 7.74E-15 | |
| 7 | rs13237637 | 3503207 | C | G | 0.4967 | 331983 | 0.9718 | 0.005 | 1.41E-08 | |
| 7 | rs1174864 | 53127559 | G | A | 0.4483 | 334045 | 0.9726 | 0.005 | 3.68E-08 | |
| 7 | rs2404324 | 99023461 | G | A | 0.1547 | 335670 | 0.9627 | 0.007 | 3.57E-08 | |
| 7 | rs10244364 | 117529641 | C | T | 0.3201 | 330304 | 1.039 | 0.005 | 1.03E-12 | |
| 8 | rs13258512 | 92777433 | G | A | 0.4221 | 331316 | 0.9709 | 0.005 | 6.40E-09 | |
| 9 | rs56209921 | 38276428 | T | C | 0.163 | 335889 | 1.039 | 0.007 | 1.42E-08 | |
| 9 | rs323740 | 102083530 | G | C | 0.4737 | 335303 | 0.9706 | 0.005 | 2.80E-09 | |
| 9 | rs7870475 | 128134034 | C | T | 0.4743 | 334109 | 1.032 | 0.005 | 6.28E-10 | |
| 10 | rs12244388 | 104640052 | A | G | 0.3357 | 335056 | 1.036 | 0.005 | 1.91E-11 | |
| 11 | rs4756044 | 45961114 | C | A | 0.2525 | 334624 | 0.9689 | 0.006 | 4.10E-08 | SV |
| 11 | rs10896972 | 59212884 | G | T | 0.3302 | 336010 | 0.9687 | 0.005 | 2.30E-09 | |

| | | | | | | | | | | |
|---|---|---|---|---|---|---|---|---|---|---|
| 11 | rs2155290 | 112851068 | G | C | 0.3826 | 335943 | 1.067 | 0.005 | 9.47E-36 | |
| 12 | rs4766578 | 111904371 | T | A | 0.4958 | 335926 | 1.028 | 0.005 | 3.62E-08 | SV |
| 13 | rs56081685 | 59454140 | G | T | 0.314 | 335337 | 0.9704 | 0.005 | 2.76E-08 | |
| 13 | rs837333 | 101179012 | C | T | 0.4763 | 326768 | 1.029 | 0.005 | 1.34E-08 | |
| 15 | rs150294 | 89931148 | G | A | 0.3996 | 329261 | 0.9701 | 0.005 | 4.04E-09 | |
| 17 | rs7216173 | 51891405 | A | T | 0.2139 | 323640 | 1.036 | 0.006 | 1.13E-08 | |
| 18 | rs1221983 | 49993266 | T | G | 0.394 | 335304 | 1.029 | 0.005 | 1.63E-08 | |

**Supplemental Table S16: Genome-wide significant loci in GWAS for smoking**

This table shows the genome-wide significant loci (top SNP in 1MB regions) in GWAS for smoking (data field 20160: “Ever smoked”). For each locus we show the chromosome (CHR), rsid for the top SNP in 1MB window (SNP), position on the chromosome (BP), test and minor allele (A1), major allele (A0), allele frequency of the test allele (A1FREQ) in all White-British samples in UKBiobank, number of samples included in the logistic regression at the locus with no missing genotype, phenotype or covariate data (NMISS), odds ratio of the minor allele on the phenotype (OR), standard error of the OR (SE), p-value of the association (P), and whether the locus is in the MHC region on chr6:25-35MB or in any SV regions as listed in Price et al 2008[9] (MHC/SV).

| CHR | SNP | BP | A1 | A0 | A1FREQ | NMISS | BETA | SE | P | MHC/SV |
|---|---|---|---|---|---|---|---|---|---|---|
| 1 | rs1002656 | 37192741 | C | T | 0.2905 | 270132 | 0.01183 | 0.001924 | 7.84E-10 | |
| 1 | rs4561025 | 91263757 | A | C | 0.3521 | 273426 | 0.01059 | 0.001912 | 3.04E-08 | |
| 1 | rs12145171 | 171843249 | C | A | 0.4283 | 273985 | -0.01105 | 0.00191 | 7.22E-09 | |
| 2 | rs12466146 | 16077199 | T | C | 0.4219 | 271236 | 0.01055 | 0.00192 | 3.88E-08 | |
| 2 | rs6743916 | 58704449 | A | G | 0.2927 | 271257 | -0.01136 | 0.001921 | 3.34E-09 | |
| 2 | rs2042555 | 148555489 | A | G | 0.418 | 273328 | 0.01323 | 0.001914 | 4.77E-12 | |
| 2 | rs7567451 | 157053380 | G | T | 0.2672 | 272984 | -0.01054 | 0.001915 | 3.67E-08 | |
| 2 | rs10497655 | 185462041 | C | T | 0.3206 | 274114 | -0.0106 | 0.001909 | 2.84E-08 | SV |
| 3 | rs2278609 | 16924440 | C | T | 0.216 | 270342 | 0.01191 | 0.001923 | 5.86E-10 | |
| 3 | rs1542212 | 35683935 | G | T | 0.3917 | 271082 | 0.01205 | 0.001921 | 3.54E-10 | |
| 3 | rs57838764 | 50374568 | C | T | 0.1142 | 273935 | 0.01062 | 0.001911 | 2.75E-08 | |
| 3 | rs836927 | 107201428 | A | C | 0.4257 | 265524 | 0.01067 | 0.001941 | 3.88E-08 | |
| 3 | rs10935184 | 136153468 | C | T | 0.4093 | 274114 | -0.01165 | 0.00191 | 1.06E-09 | |
| 4 | rs13102162 | 90939567 | G | A | 0.3591 | 264987 | -0.01124 | 0.001942 | 7.15E-09 | |
| 4 | rs10032297 | 139013401 | A | T | 0.3958 | 269788 | -0.01136 | 0.001925 | 3.54E-09 | |
| 4 | rs7696796 | 183166469 | A | G | 0.2464 | 274114 | 0.01116 | 0.00191 | 5.05E-09 | |
| 5 | rs4413518 | 107738001 | A | G | 0.2189 | 271136 | 0.01234 | 0.00192 | 1.29E-10 | |
| 6 | rs2148254 | 11994762 | G | A | 0.4928 | 270744 | -0.01133 | 0.001921 | 3.67E-09 | |
| 6 | rs7772160 | 27412386 | C | T | 0.4783 | 274114 | -0.01061 | 0.00191 | 2.72E-08 | MHC |
| 6 | rs2269426 | 32076499 | A | G | 0.3594 | 274114 | 0.01334 | 0.001911 | 2.95E-12 | MHC |
| 6 | rs6916891 | 98457595 | T | G | 0.1176 | 271957 | 0.01064 | 0.001917 | 2.89E-08 | |
| 7 | rs56226325 | 2078981 | T | C | 0.1534 | 274114 | -0.01052 | 0.00191 | 3.61E-08 | |
| 7 | rs11509880 | 12261911 | A | G | 0.3278 | 273194 | 0.01211 | 0.001913 | 2.43E-10 | |
| 7 | rs73167875 | 82939096 | A | T | 0.2022 | 269308 | -0.01083 | 0.001926 | 1.90E-08 | |
| 7 | rs274632 | 86269181 | A | C | 0.4229 | 273225 | -0.01153 | 0.001915 | 1.71E-09 | |
| 7 | rs6976111 | 117495667 | A | C | 0.2993 | 264510 | 0.01101 | 0.001943 | 1.45E-08 | |
| 7 | rs13226841 | 126389408 | C | T | 0.4896 | 274064 | 0.0125 | 0.00191 | 6.03E-11 | |
| 8 | rs7845515 | 4946228 | A | G | 0.2875 | 265208 | 0.01267 | 0.001941 | 6.69E-11 | |

| | | | | | | | | | | |
|---|---|---|---|---|---|---|---|---|---|---|
| 8 | rs2921036 | 8363897 | T | C | 0.4898 | 269177 | 0.01775 | 0.001927 | 3.32E-20 | SV |
| 8 | rs477860 | 9811765 | T | C | 0.2968 | 271822 | -0.01439 | 0.001918 | 6.25E-14 | SV |
| 8 | rs11250117 | 10972740 | A | C | 0.4667 | 272432 | 0.0172 | 0.001915 | 2.69E-19 | SV |
| 9 | rs2380937 | 4145781 | C | T | 0.4011 | 272440 | -0.01119 | 0.001916 | 5.23E-09 | |
| 9 | rs4977844 | 23295899 | C | G | 0.3598 | 266189 | 0.01136 | 0.001937 | 4.48E-09 | |
| 9 | rs7869969 | 96217447 | G | A | 0.3321 | 274057 | -0.01087 | 0.00191 | 1.28E-08 | |
| 9 | rs28377268 | 98225056 | T | G | 0.1071 | 272736 | 0.01142 | 0.001914 | 2.42E-09 | |
| 9 | rs2094580 | 120490857 | G | T | 0.2825 | 270567 | 0.01071 | 0.001922 | 2.50E-08 | |
| 11 | rs34796300 | 13315205 | T | C | 0.4254 | 272562 | 0.01137 | 0.001916 | 2.96E-09 | |
| 11 | rs297343 | 16354653 | T | G | 0.3604 | 272965 | 0.01107 | 0.001914 | 7.23E-09 | |
| 11 | rs2071754 | 31812582 | C | T | 0.201 | 274114 | 0.01262 | 0.00191 | 3.94E-11 | |
| 11 | rs7107356 | 47676170 | A | G | 0.4943 | 273602 | -0.01319 | 0.001911 | 5.21E-12 | SV |
| 11 | rs10896636 | 57448032 | G | C | 0.3449 | 268952 | 0.01281 | 0.001928 | 3.02E-11 | |
| 11 | rs674437 | 88689953 | G | A | 0.4843 | 273301 | 0.01098 | 0.001913 | 9.46E-09 | SV |
| 11 | rs35738585 | 113386347 | G | T | 0.4334 | 271900 | -0.01561 | 0.001918 | 3.95E-16 | |
| 12 | rs11608355 | 109879292 | C | T | 0.3132 | 272760 | 0.01181 | 0.001914 | 6.75E-10 | SV |
| 12 | rs3741475 | 117669914 | A | G | 0.1944 | 273597 | 0.01194 | 0.001911 | 4.14E-10 | |
| 13 | rs2210903 | 69576975 | G | A | 0.3692 | 274114 | -0.01168 | 0.00191 | 9.64E-10 | |
| 14 | rs4140799 | 72170969 | G | A | 0.47 | 272113 | 0.01268 | 0.001917 | 3.68E-11 | |
| 14 | rs10144845 | 75237770 | C | T | 0.317 | 274031 | -0.01389 | 0.001911 | 3.60E-13 | |
| 15 | rs12903563 | 78033735 | T | C | 0.4798 | 269275 | 0.01176 | 0.001927 | 1.03E-09 | |
| 17 | rs12938775 | 2574821 | G | A | 0.4989 | 274114 | 0.0114 | 0.00191 | 2.36E-09 | |
| 17 | rs35982947 | 38214275 | C | A | 0.3692 | 267464 | -0.01109 | 0.001933 | 9.51E-09 | |
| 17 | rs62062288 | 44096553 | A | G | 0.2181 | 267423 | 0.01863 | 0.001937 | 6.72E-22 | |
| 17 | rs56084168 | 79084574 | T | C | 0.1478 | 273326 | -0.0123 | 0.001913 | 1.27E-10 | |
| 18 | rs9952522 | 31286129 | C | T | 0.4084 | 272613 | -0.01074 | 0.001915 | 2.04E-08 | |
| 18 | rs11665070 | 35152563 | G | A | 0.3313 | 272412 | 0.01565 | 0.001916 | 3.10E-16 | |
| 18 | rs8097041 | 50898217 | A | T | 0.3644 | 273465 | -0.01159 | 0.001953 | 2.95E-09 | |
| 18 | rs56403421 | 52765283 | C | A | 0.3309 | 268579 | 0.01207 | 0.001931 | 4.15E-10 | |

| | | | | | | | | | |
|---|---|---|---|---|---|---|---|---|---|
| 22 | rs1028321 | 39926929 | A | G | 0.3052 | 272405 | -0.0111 | 0.001918 | 7.09E-09 |
| 22 | rs11090045 | 41753603 | A | G | 0.3031 | 263768 | 0.01321 | 0.001945 | 1.13E-11 |

**Supplemental Table S17: Genome-wide significant loci in GWAS for neuroticism**

This table shows the genome-wide significant loci (top SNP in 1MB regions) in GWAS for neuroticism score (data field 20127: "Neuroticism score"). For each locus we show the chromosome (CHR), rsid for the top SNP in 1MB window (SNP), position on the chromosome (BP), test and minor allele (A1), major allele (A0), allele frequency of the test allele (A1FREQ) in all White-British samples in UKBiobank, number of samples included in the linear regression at the locus with no missing genotype, phenotype or covariate data (NMISS), standardized effect size of the minor allele on the phenotype (BETA), standard error of the effect (SE), p-value of the association (P), and whether the locus is in the MHC region on chr6:25-35MB or in any SV regions as listed in Price et al 2008[9] (MHC/SV).

| CONDITION | CATEGORY | DEFINITION | LDSC | | | | rho-HESS | |
|---|---|---|---|---|---|---|---|---|
| | | | INTERCEPT | RHO (SE) | RG (SE) | P | RHO (SE) | RG (SE) |
| ADHD | CIDI-based | LifetimeMDD | 0.022 | 0.048 (0.011) | 0.297 (0.076) | 8.93E-05 | 0.054 (0.016) | 0.271 (0.026) |
| | Help-seeking | GPpsy | 0.026 | 0.058 (0.007) | 0.389 (0.045) | 5.47E-18 | 0.043 (0.008) | 0.272 (0.016) |
| | Symptom | DepAll | 0.026 | 0.038 (0.011) | 0.280 (0.081) | 5.00E-04 | 0.044 (0.015) | 0.263 (0.029) |
| | Self-report | SelfRepDep | 0.017 | 0.017 (0.006) | 0.227 (0.077) | 3.20E-03 | 0.020 (0.009) | 0.217 (0.030) |
| | EMR | ICD10Dep | 0.011 | 0.039 (0.006) | 0.503 (0.072) | 3.66E-12 | 0.030 (0.010) | 0.334 (0.035) |
| | No-MDD | GPNoDep | 0.008 | 0.027 (0.010) | 0.307 (0.224) | 1.71E-01 | 0.022 (0.019) | 0.206 (0.053) |
| BIP | CIDI-based | LifetimeMDD | 0.019 | 0.043 (0.013) | 0.344 (0.095) | 3.00E-04 | 0.038 (0.012) | 0.286 (0.035) |
| | Help-seeking | GPpsy | 0.026 | 0.033 (0.006) | 0.320 (0.052) | 5.19E-10 | 0.028 (0.006) | 0.264 (0.021) |
| | Symptom | DepAll | 0.001 | 0.052 (0.009) | 0.534 (0.091) | 4.66E-09 | 0.034 (0.011) | 0.290 (0.036) |
| | Self-report | SelfRepDep | 0.021 | 0.020 (0.005) | 0.341 (0.088) | 1.00E-04 | 0.019 (0.007) | 0.291 (0.037) |
| | EMR | ICD10Dep | 0.003 | 0.019 (0.005) | 0.306 (0.085) | 3.00E-04 | 0.014 (0.007) | 0.225 (0.042) |
| | No-MDD | GPNoDep | 0.003 | 0.021 (0.011) | 0.305 (0.155) | 4.87E-02 | 0.011 (0.014) | 0.163 (0.070) |
| SCZ | CIDI-based | LifetimeMDD | 0.023 | 0.061 (0.010) | 0.372 (0.055) | 1.38E-11 | 0.059 (0.008) | 0.247 (0.017) |
| | Help-seeking | GPpsy | 0.028 | 0.046 (0.007) | 0.326 (0.037) | 5.01E-19 | 0.038 (0.004) | 0.197 (0.010) |
| | Symptom | DepAll | 0.010 | 0.056 (0.009) | 0.421 (0.053) | 2.08E-15 | 0.041 (0.007) | 0.201 (0.017) |
| | Self-report | SelfRepDep | 0.019 | 0.028 (0.005) | 0.357 (0.059) | 1.11E-09 | 0.025 (0.004) | 0.226 (0.018) |
| | EMR | ICD10Dep | 0.001 | 0.030 (0.004) | 0.354 (0.049) | 5.51E-13 | 0.019 (0.005) | 0.180 (0.020) |
| | No-MDD | GPNoDep | 0.006 | 0.031 (0.010) | 0.351 (0.111) | 1.60E-03 | 0.024 (0.009) | 0.203 (0.035) |
| AUT | CIDI-based | LifetimeMDD | 0.010 | 0.030 (0.016) | 0.164 (0.090) | 6.66E-02 | 0.039 (0.040) | 0.295 (0.077) |
| | Help-seeking | GPpsy | 0.014 | 0.005 (0.009) | 0.033 (0.057) | 5.56E-01 | 0.016 (0.019) | 0.147 (0.047) |

| | | | | | | | | |
|---|---|---|---|---|---|---|---|---|
| | Symptom | DepAll | -0.002 | 0.041 (0.014) | 0.272 (0.100) | 6.30E-03 | 0.017 (0.039) | 0.146 (0.080) |
| | Self-report | SelfRepDep | 0.007 | 0.009 (0.009) | 0.098 (0.096) | 3.06E-01 | 0.014 (0.023) | 0.227 (0.084) |
| | EMR | ICD10Dep | 0.001 | -0.008 (0.009) | -0.077 (0.091) | 3.98E-01 | 0.002 (0.025) | 0.032 (0.095) |
| | No-MDD | GPNoDep | 0.011 | -0.023 (0.017) | -0.225 (0.175) | 1.98E-01 | 0.009 (0.048) | 0.146 (0.168) |
| PGC1-MDD | CIDI-based | LifetimeMDD | 0.008 | 0.107 (0.013) | 0.857 (0.127) | 1.51E-11 | 0.076 (0.039) | 0.796 (0.099) |
| | Help-seeking | GPpsy | 0.013 | 0.103 (0.008) | 0.940 (0.100) | 6.52E-21 | 0.077 (0.018) | 0.996 (0.073) |
| | Symptom | DepAll | 0.003 | 0.103 (0.012) | 0.968 (0.130) | 1.12E-13 | 0.069 (0.037) | 0.856 (0.104) |
| | Self-report | SelfRepDep | 0.016 | 0.050 (0.008) | 0.811 (0.124) | 6.27E-11 | 0.048 (0.022) | 1.110 (0.117) |
| | EMR | ICD10Dep | 0.010 | 0.044 (0.007) | 0.673 (0.114) | 3.80E-09 | 0.039 (0.024) | 0.935 (0.126) |
| | No-MDD | GPNoDep | -0.003 | 0.073 (0.013) | 1.058 (0.265) | 6.46E-05 | 0.040 (0.046) | 0.809 (0.198) |
| 23andMe (depression) | CIDI-based | LifetimeMDD | 0.020 | 0.066 (0.006) | 0.826 (0.060) | 6.41E-43 | 0.049 (0.010) | 0.512 (0.025) |
| | Help-seeking | GPpsy | 0.045 | 0.057 (0.003) | 0.853 (0.030) | 8.74E-179 | 0.044 (0.004) | 0.571 (0.015) |
| | Symptom | DepAll | 0.028 | 0.048 (0.005) | 0.732 (0.064) | 3.53E-30 | 0.041 (0.009) | 0.503 (0.028) |
| | Self-report | SelfRepDep | 0.030 | 0.028 (0.002) | 0.735 (0.062) | 2.24E-32 | 0.025 (0.005) | 0.568 (0.032) |
| | EMR | ICD10Dep | 0.015 | 0.023 (0.002) | 0.599 (0.066) | 1.07E-19 | 0.018 (0.006) | 0.422 (0.035) |
| | No-MDD | GPNoDep | 0.015 | 0.031 (0.005) | 0.680 (0.141) | 1.42E-06 | 0.026 (0.011) | 0.533 (0.059) |
| Neuroticism | CIDI-based | LifetimeMDD | 0.179 | 0.080 (0.007) | 0.671 (0.041) | 2.69E-60 | 0.156 (0.009) | 1.150 (0.025) |
| | Help-seeking | GPpsy | 0.384 | 0.072 (0.005) | 0.705 (0.019) | 1.88E-298 | 0.147 (0.004) | 1.350 (0.015) |
| | Symptom | DepAll | 0.161 | 0.063 (0.007) | 0.617 (0.043) | 2.22E-46 | 0.126 (0.009) | 1.090 (0.028) |
| | Self-report | SelfRepDep | 0.231 | 0.041 (0.004) | 0.717 (0.040) | 2.48E-71 | 0.096 (0.005) | 1.570 (0.053) |
| | EMR | ICD10Dep | 0.159 | 0.029 (0.003) | 0.499 (0.060) | 5.70E-17 | 0.070 (0.006) | 1.210 (0.057) |
| | No-MDD | GPNoDep | 0.109 | 0.046 (0.007) | 0.642 (0.128) | 5.69E-07 | 0.099 (0.011) | 1.370 (0.092) |

| | | | | | | | | |
|---|---|---|---|---|---|---|---|---|
| Smoking | CIDI-based | LifetimeMDD | 0.037 | 0.019 (0.006) | 0.192 (0.055) | 5.00E-04 | 0.030 (0.009) | 0.283 (0.022) |
| | Help-seeking | GPpsy | 0.071 | 0.023 (0.004) | 0.279 (0.037) | 3.95E-14 | 0.031 (0.004) | 0.365 (0.014) |
| | Symptom | DepAll | 0.040 | 0.023 (0.005) | 0.286 (0.057) | 5.79E-07 | 0.031 (0.009) | 0.348 (0.025) |
| | Self-report | SelfRepDep | 0.022 | 0.008 (0.003) | 0.182 (0.052) | 5.00E-04 | 0.012 (0.005) | 0.246 (0.026) |
| | EMR | ICD10Dep | 0.016 | 0.015 (0.002) | 0.331 (0.047) | 2.55E-12 | 0.014 (0.006) | 0.301 (0.031) |
| | No-MDD | GPNoDep | 0.012 | 0.015 (0.007) | 0.262 (0.100) | 8.80E-03 | 0.015 (0.011) | 0.274 (0.045) |

**Supplemental Table S18: Genetic correlation between definitions of MDD and other conditions**

This table shows the genetic correlation between each definition of MDD in UKBiobank with other conditions: attention deficit hyperactive disorder (ADHD), bipolar disorder (BIP), schizophrenia (SCZ), autism (AUT), first meta-analysis of MDD by PGC (PGC1), minimal phenotyping based MDD study by 23andMe (23andMe), and neuroticism and smoking in UKBiobank. This table shows results from two different methods: LDSC and rho-HESS. For each of the methods we show the genetic covariance (rho) and its standard error (rho_se), as well as the genetic correlation (rG) and its standard error (rG_se); genetic correlation is genetic covariance divided by the product of the heritabilities of the pair of traits involved. For estimates from LDSC, we show the p-value (P) for the genetic correlation being different from 0. For estimates from rho-HESS, both genetic covariance and genetic correlation are summed from regional genetic covariance and genetic correlations in 1703 independent genomic regions (Supplemental Methods). The Pearson r between rG estimated from LDSC and rho-HESS is 0.70 ($P=2.48 \times 10^{-8}$).

| CATEGORY | PHENOTYPE | SNP | CHR | BP | A1 | OR | L95 | U95 | P | REPLICATION |
|---|---|---|---|---|---|---|---|---|---|---|
| Help seeking | GPpsy | rs6699744 | 1 | 72825144 | A | 0.960 | 0.950 | 0.971 | 6.55E-14 | Replicated |
| | | rs6697602 | 1 | 177039372 | G | 1.056 | 1.037 | 1.075 | 6.36E-09 | Replicated |
| | | rs11123030 | 2 | 124976163 | T | 1.032 | 1.022 | 1.043 | 1.13E-09 | Replicated |
| | | rs66511648 | 3 | 117515519 | C | 1.033 | 1.021 | 1.045 | 2.42E-08 | Replicated |
| | | rs30266 | 5 | 103972357 | A | 1.039 | 1.028 | 1.050 | 3.50E-12 | Replicated |
| | | rs12205083 | 6 | 24275483 | G | 1.053 | 1.036 | 1.071 | 6.24E-10 | Replicated |
| | | rs75782365 | 6 | 26408551 | G | 0.951 | 0.936 | 0.967 | 1.36E-09 | Replicated |
| | | rs7772160 | 6 | 27412386 | C | 0.965 | 0.955 | 0.974 | 3.27E-12 | Replicated |
| | | rs4713145 | 6 | 28106827 | C | 0.969 | 0.957 | 0.980 | 1.51E-07 | Replicated |
| | | rs3135296 | 6 | 28795856 | T | 0.946 | 0.931 | 0.961 | 3.75E-12 | Replicated |
| | | rs3129120 | 6 | 29111775 | C | 0.948 | 0.934 | 0.963 | 1.48E-11 | Replicated |
| | | rs3115631 | 6 | 29986324 | A | 0.944 | 0.929 | 0.959 | 2.86E-13 | Replicated |
| | | rs2517622 | 6 | 30155149 | C | 0.950 | 0.936 | 0.964 | 1.26E-11 | Replicated |
| | | rs1625792 | 6 | 31306420 | A | 0.961 | 0.947 | 0.975 | 4.59E-08 | Replicated |
| | | rs535777 | 6 | 32577633 | C | 0.965 | 0.951 | 0.980 | 4.66E-06 | Replicated |
| | | rs236346 | 6 | 36832103 | C | 0.950 | 0.934 | 0.967 | 1.19E-08 | Replicated |
| | | rs9345737 | 6 | 66676938 | G | 0.969 | 0.960 | 0.979 | 3.03E-09 | Replicated |
| | | rs3807866 | 7 | 12250378 | A | 1.039 | 1.028 | 1.049 | 5.44E-13 | Replicated |
| | | rs393488 | 9 | 17044971 | A | 0.967 | 0.958 | 0.977 | 2.14E-10 | Replicated |
| | | rs12057031 | 9 | 25235063 | T | 0.952 | 0.936 | 0.968 | 5.29E-09 | Replicated |
| | | rs11599236 | 10 | 106454672 | C | 0.971 | 0.961 | 0.981 | 2.91E-08 | Replicated |
| | | rs537635 | 11 | 88705235 | T | 1.033 | 1.022 | 1.043 | 6.24E-10 | Replicated |
| | | rs578174 | 11 | 89959637 | G | 0.953 | 0.937 | 0.969 | 2.10E-08 | Replicated |
| | | rs12889665 | 14 | 75234830 | T | 0.972 | 0.962 | 0.982 | 3.16E-08 | Replicated |
| | | rs61997596 | 14 | 104511206 | A | 1.037 | 1.023 | 1.050 | 4.15E-08 | Replicated |
| | | rs11646401 | 16 | 21609978 | G | 1.029 | 1.019 | 1.040 | 3.58E-08 | Replicated |
| | | rs12967855 | 18 | 35138245 | A | 1.034 | 1.023 | 1.045 | 1.57E-09 | Replicated |
| DSM based | LifetimeMDD | rs6699744 | 1 | 72825144 | A | 0.984 | 0.959 | 1.010 | 2.20E-01 | NotReplicated |
| | | rs6697602 | 1 | 177039372 | G | 1.061 | 1.015 | 1.110 | 9.30E-03 | NotReplicated |
| | | rs11123030 | 2 | 124976163 | T | 1.024 | 0.999 | 1.050 | 6.35E-02 | NotReplicated |

| | | | | | | | | | | |
|---|---|---|---|---|---|---|---|---|---|---|
| | | rs66511648 | 3 | 117515519 | C | 1.034 | 1.006 | 1.064 | 1.79E-02 | NotReplicated |
| | | rs30266 | 5 | 103972357 | A | 1.047 | 1.020 | 1.076 | 6.55E-04 | Replicated |
| | | rs12205083 | 6 | 24275483 | G | 0.992 | 0.952 | 1.033 | 6.89E-01 | NotReplicated |
| | | rs75782365 | 6 | 26408551 | G | 0.939 | 0.902 | 0.978 | 2.32E-03 | NotReplicated |
| | | rs7772160 | 6 | 27412386 | C | 0.957 | 0.934 | 0.982 | 6.19E-04 | Replicated |
| | | rs4713145 | 6 | 28106827 | C | 0.962 | 0.934 | 0.990 | 9.04E-03 | NotReplicated |
| | | rs3135296 | 6 | 28795856 | T | 0.905 | 0.870 | 0.941 | 5.48E-07 | Replicated |
| | | rs3129120 | 6 | 29111775 | C | 0.905 | 0.871 | 0.941 | 4.08E-07 | Replicated |
| | | rs3115631 | 6 | 29986324 | A | 0.906 | 0.872 | 0.941 | 4.49E-07 | Replicated |
| | | rs2517622 | 6 | 30155149 | C | 0.923 | 0.890 | 0.958 | 2.10E-05 | Replicated |
| | | rs1625792 | 6 | 31306420 | A | 0.939 | 0.906 | 0.973 | 4.91E-04 | Replicated |
| | | rs535777 | 6 | 32577633 | C | 0.952 | 0.918 | 0.988 | 9.68E-03 | NotReplicated |
| | | rs236346 | 6 | 36832103 | C | 0.955 | 0.915 | 0.997 | 3.65E-02 | NotReplicated |
| | | rs9345737 | 6 | 66676938 | G | 0.994 | 0.970 | 1.020 | 6.63E-01 | NotReplicated |
| | | rs3807866 | 7 | 12250378 | A | 1.049 | 1.023 | 1.076 | 2.33E-04 | Replicated |
| | | rs393488 | 9 | 17044971 | A | 0.976 | 0.952 | 1.001 | 6.00E-02 | NotReplicated |
| | | rs12057031 | 9 | 25235063 | T | 0.938 | 0.900 | 0.977 | 1.94E-03 | NotReplicated |
| | | rs11599236 | 10 | 106454672 | C | 0.970 | 0.945 | 0.996 | 2.13E-02 | NotReplicated |
| | | rs537635 | 11 | 88705235 | T | 1.041 | 1.016 | 1.068 | 1.44E-03 | Replicated |
| | | rs578174 | 11 | 89959637 | G | 0.974 | 0.934 | 1.016 | 2.17E-01 | NotReplicated |
| | | rs12889665 | 14 | 75234830 | T | 0.973 | 0.949 | 0.998 | 3.31E-02 | NotReplicated |
| | | rs61997596 | 14 | 104511206 | A | 1.064 | 1.031 | 1.099 | 1.18E-04 | Replicated |
| | | rs11646401 | 16 | 21609978 | G | 1.019 | 0.994 | 1.045 | 1.37E-01 | NotReplicated |
| | | rs12967855 | 18 | 35138245 | A | 1.001 | 0.975 | 1.029 | 9.23E-01 | NotReplicated |
| | | rs6699744 | 1 | 72825144 | A | 0.972 | 0.940 | 1.006 | 1.02E-01 | NotReplicated |
| | | rs6697602 | 1 | 177039372 | G | 1.073 | 1.013 | 1.136 | 1.61E-02 | NotReplicated |
| | | rs11123030 | 2 | 124976163 | T | 1.019 | 0.986 | 1.052 | 2.63E-01 | NotReplicated |
| **No-MDD** | **GPNoDep** | rs66511648 | 3 | 117515519 | C | 1.016 | 0.979 | 1.053 | 4.02E-01 | NotReplicated |
| | | rs30266 | 5 | 103972357 | A | 1.047 | 1.012 | 1.084 | 8.94E-03 | NotReplicated |
| | | rs12205083 | 6 | 24275483 | G | 1.043 | 0.989 | 1.099 | 1.18E-01 | NotReplicated |
| | | rs75782365 | 6 | 26408551 | G | 0.907 | 0.860 | 0.956 | 2.98E-04 | Replicated |

| | | | | | | | | | |
|---|---|---|---|---|---|---|---|---|---|
| | | rs7772160 | 6 | 27412386 | C | 0.930 | 0.900 | 0.960 | 1.05E-05 | Replicated |
| | | rs4713145 | 6 | 28106827 | C | 0.919 | 0.885 | 0.955 | 1.33E-05 | Replicated |
| | | rs3135296 | 6 | 28795856 | T | 0.881 | 0.837 | 0.928 | 1.49E-06 | Replicated |
| | | rs3129120 | 6 | 29111775 | C | 0.895 | 0.851 | 0.941 | 1.54E-05 | Replicated |
| | | rs3115631 | 6 | 29986324 | A | 0.869 | 0.826 | 0.914 | 5.62E-08 | Replicated |
| | | rs2517622 | 6 | 30155149 | C | 0.891 | 0.849 | 0.935 | 2.80E-06 | Replicated |
| | | rs1625792 | 6 | 31306420 | A | 0.913 | 0.872 | 0.957 | 1.38E-04 | Replicated |
| | | rs535777 | 6 | 32577633 | C | 0.926 | 0.882 | 0.973 | 2.15E-03 | NotReplicated |
| | | rs236346 | 6 | 36832103 | C | 0.990 | 0.937 | 1.046 | 7.21E-01 | NotReplicated |
| | | rs9345737 | 6 | 66676938 | G | 0.949 | 0.919 | 0.981 | 1.77E-03 | Replicated |
| | | rs3807866 | 7 | 12250378 | A | 1.024 | 0.991 | 1.058 | 1.59E-01 | NotReplicated |
| | | rs393488 | 9 | 17044971 | A | 0.964 | 0.933 | 0.996 | 2.86E-02 | NotReplicated |
| | | rs12057031 | 9 | 25235063 | T | 0.991 | 0.940 | 1.044 | 7.24E-01 | NotReplicated |
| | | rs11599236 | 10 | 106454672 | C | 0.985 | 0.953 | 1.019 | 3.82E-01 | NotReplicated |
| | | rs537635 | 11 | 88705235 | T | 0.999 | 0.967 | 1.031 | 9.33E-01 | NotReplicated |
| | | rs578174 | 11 | 89959637 | G | 0.962 | 0.911 | 1.016 | 1.66E-01 | NotReplicated |
| | | rs12889665 | 14 | 75234830 | T | 0.989 | 0.957 | 1.022 | 5.08E-01 | NotReplicated |
| | | rs61997596 | 14 | 104511206 | A | 1.043 | 1.001 | 1.087 | 4.57E-02 | NotReplicated |
| | | rs11646401 | 16 | 21609978 | G | 1.047 | 1.014 | 1.082 | 5.11E-03 | NotReplicated |
| | | rs12967855 | 18 | 35138245 | A | 1.008 | 0.974 | 1.044 | 6.49E-01 | NotReplicated |
| **Other Condition** | **SCZ** | rs6699744 | 1 | 72825144 | A | 0.978 | 0.956 | 1.000 | 4.34E-02 | NotReplicated |
| | | rs6697602 | 1 | 177039372 | G | 0.993 | 0.955 | 1.030 | 7.06E-01 | NotReplicated |
| | | rs11123030 | 2 | 124976163 | T | 1.026 | 1.006 | 1.047 | 1.36E-02 | NotReplicated |
| | | rs66511648 | 3 | 117515519 | C | 0.997 | 0.973 | 1.021 | 8.26E-01 | NotReplicated |
| | | rs30266 | 5 | 103972357 | A | 1.023 | 1.000 | 1.045 | 5.02E-02 | NotReplicated |
| | | rs12205083 | 6 | 24275483 | G | 0.985 | 0.952 | 1.019 | 3.92E-01 | NotReplicated |
| | | rs75782365 | 6 | 26408551 | G | 0.754 | 0.714 | 0.794 | 8.89E-27 | Replicated |
| | | rs7772160 | 6 | 27412386 | C | 0.981 | 0.960 | 1.001 | 6.87E-02 | NotReplicated |
| | | rs4713145 | 6 | 28106827 | C | 0.923 | 0.898 | 0.948 | 8.73E-09 | Replicated |
| | | rs3135296 | NA | NA | T | NA | NA | NA | NA | NotReplicated |
| | | rs3129120 | NA | NA | C | NA | NA | NA | NA | NotReplicated |

| | | | | | | | | | | |
|---|---|---|---|---|---|---|---|---|---|---|
| | | rs3115631 | NA | NA | A | NA | NA | NA | NA | NotReplicated |
| | | rs2517622 | NA | NA | C | NA | NA | NA | NA | NotReplicated |
| | | rs1625792 | NA | NA | A | NA | NA | NA | NA | NotReplicated |
| | | rs535777 | 6 | 32577633 | C | 0.874 | 0.839 | 0.910 | 1.78E-13 | Replicated |
| | | rs236346 | 6 | 36832103 | C | 0.999 | 0.965 | 1.034 | 9.72E-01 | NotReplicated |
| | | rs9345737 | 6 | 66676938 | G | 0.976 | 0.955 | 0.997 | 2.84E-02 | NotReplicated |
| | | rs3807866 | 7 | 12250378 | A | 1.016 | 0.995 | 1.037 | 1.36E-01 | NotReplicated |
| | | rs393488 | 9 | 17044971 | A | 0.982 | 0.961 | 1.003 | 9.64E-02 | NotReplicated |
| | | rs12057031 | 9 | 25235063 | T | 0.954 | 0.920 | 0.989 | 7.57E-03 | NotReplicated |
| | | rs11599236 | 10 | 106454672 | C | 0.961 | 0.939 | 0.982 | 3.63E-04 | Replicated |
| | | rs537635 | 11 | 88705235 | T | 1.014 | 0.993 | 1.035 | 1.97E-01 | NotReplicated |
| | | rs578174 | 11 | 89959637 | G | 0.989 | 0.951 | 1.027 | 5.83E-01 | NotReplicated |
| | | rs12889665 | 14 | 75234830 | T | 0.981 | 0.960 | 1.002 | 7.26E-02 | NotReplicated |
| | | rs61997596 | 14 | 104511206 | A | 1.057 | 1.030 | 1.085 | 7.48E-05 | Replicated |
| | | rs11646401 | 16 | 21609978 | G | 1.026 | 1.004 | 1.048 | 1.66E-02 | NotReplicated |
| | | rs12967855 | 18 | 35138245 | A | 1.015 | 0.993 | 1.037 | 1.77E-01 | NotReplicated |
| **Other Condition** | **Smoking** | rs6699744 | 1 | 72825144 | A | 0.996 | 0.986 | 1.006 | 4.22E-01 | NotReplicated |
| | | rs6697602 | 1 | 177039372 | G | 1.001 | 0.984 | 1.019 | 8.82E-01 | NotReplicated |
| | | rs11123030 | 2 | 124976163 | T | 0.998 | 0.988 | 1.008 | 6.74E-01 | NotReplicated |
| | | rs66511648 | 3 | 117515519 | C | 1.000 | 0.990 | 1.012 | 9.33E-01 | NotReplicated |
| | | rs30266 | 5 | 103972357 | A | 1.020 | 1.009 | 1.031 | 2.01E-04 | Replicated |
| | | rs12205083 | 6 | 24275483 | G | 1.005 | 0.989 | 1.021 | 5.48E-01 | NotReplicated |
| | | rs75782365 | 6 | 26408551 | G | 0.967 | 0.952 | 0.982 | 2.55E-05 | Replicated |
| | | rs7772160 | 6 | 27412386 | C | 0.982 | 0.973 | 0.992 | 3.08E-04 | Replicated |
| | | rs4713145 | 6 | 28106827 | C | 0.978 | 0.967 | 0.989 | 1.53E-04 | Replicated |
| | | rs3135296 | 6 | 28795856 | T | 0.970 | 0.955 | 0.984 | 5.95E-05 | Replicated |
| | | rs3129120 | 6 | 29111775 | C | 0.970 | 0.956 | 0.985 | 6.00E-05 | Replicated |
| | | rs3115631 | 6 | 29986324 | A | 0.979 | 0.964 | 0.993 | 4.67E-03 | NotReplicated |
| | | rs2517622 | 6 | 30155149 | C | 0.976 | 0.962 | 0.990 | 6.61E-04 | Replicated |
| | | rs1625792 | 6 | 31306420 | A | 0.979 | 0.966 | 0.993 | 3.04E-03 | NotReplicated |
| | | rs535777 | 6 | 32577633 | C | 0.981 | 0.967 | 0.995 | 9.58E-03 | NotReplicated |

| | | | | | | | | | | |
|---|---|---|---|---|---|---|---|---|---|---|
| | | rs236346 | 6 | 36832103 | C | 0.992 | 0.976 | 1.009 | 3.73E-01 | NotReplicated |
| | | rs9345737 | 6 | 66676938 | G | 0.994 | 0.984 | 1.004 | 2.07E-01 | NotReplicated |
| | | rs3807866 | 7 | 12250378 | A | 1.004 | 0.995 | 1.015 | 3.78E-01 | NotReplicated |
| | | rs393488 | 9 | 17044971 | A | 0.997 | 0.987 | 1.007 | 5.18E-01 | NotReplicated |
| | | rs12057031 | 9 | 25235063 | T | 0.982 | 0.966 | 0.998 | 2.29E-02 | NotReplicated |
| | | rs11599236 | 10 | 106454672 | C | 0.988 | 0.978 | 0.998 | 1.79E-02 | NotReplicated |
| | | rs537635 | 11 | 88705235 | T | 1.009 | 0.999 | 1.019 | 7.48E-02 | NotReplicated |
| | | rs578174 | 11 | 89959637 | G | 0.982 | 0.967 | 0.999 | 3.19E-02 | NotReplicated |
| | | rs12889665 | 14 | 75234830 | T | 0.988 | 0.978 | 0.998 | 1.38E-02 | NotReplicated |
| | | rs61997596 | 14 | 104511206 | A | 1.010 | 0.997 | 1.023 | 1.19E-01 | NotReplicated |
| | | rs11646401 | 16 | 21609978 | G | 1.000 | 0.990 | 1.010 | 9.81E-01 | NotReplicated |
| | | rs12967855 | 18 | 35138245 | A | 1.011 | 1.000 | 1.022 | 4.10E-02 | NotReplicated |
| **Other Condition** | **Neuroticism** | rs6699744 | 1 | 72825144 | A | 0.996 | 0.993 | 1.000 | 5.50E-02 | NotReplicated |
| | | rs6697602 | 1 | 177039372 | G | 1.006 | 1.002 | 1.010 | 2.75E-03 | NotReplicated |
| | | rs11123030 | 2 | 124976163 | T | 1.006 | 1.002 | 1.010 | 2.23E-03 | NotReplicated |
| | | rs66511648 | 3 | 117515519 | C | 1.005 | 1.001 | 1.009 | 1.33E-02 | NotReplicated |
| | | rs30266 | 5 | 103972357 | A | 1.009 | 1.005 | 1.013 | 3.75E-06 | Replicated |
| | | rs12205083 | 6 | 24275483 | G | 1.008 | 1.004 | 1.012 | 3.17E-05 | Replicated |
| | | rs75782365 | 6 | 26408551 | G | 0.992 | 0.988 | 0.996 | 3.35E-05 | Replicated |
| | | rs7772160 | 6 | 27412386 | C | 0.989 | 0.986 | 0.993 | 2.72E-08 | Replicated |
| | | rs4713145 | 6 | 28106827 | C | 0.993 | 0.989 | 0.996 | 9.14E-05 | Replicated |
| | | rs3135296 | 6 | 28795856 | T | 0.991 | 0.988 | 0.995 | 6.03E-06 | Replicated |
| | | rs3129120 | 6 | 29111775 | C | 0.992 | 0.988 | 0.995 | 1.53E-05 | Replicated |
| | | rs3115631 | 6 | 29986324 | A | 0.992 | 0.988 | 0.996 | 3.67E-05 | Replicated |
| | | rs2517622 | 6 | 30155149 | C | 0.993 | 0.989 | 0.997 | 2.15E-04 | Replicated |
| | | rs1625792 | 6 | 31306420 | A | 0.994 | 0.990 | 0.997 | 7.07E-04 | Replicated |
| | | rs535777 | 6 | 32577633 | C | 0.992 | 0.988 | 0.996 | 4.71E-05 | Replicated |
| | | rs236346 | 6 | 36832103 | C | 0.995 | 0.991 | 0.999 | 8.75E-03 | NotReplicated |
| | | rs9345737 | 6 | 66676938 | G | 0.997 | 0.993 | 1.001 | 9.08E-02 | NotReplicated |
| | | rs3807866 | 7 | 12250378 | A | 1.012 | 1.008 | 1.016 | 9.08E-10 | Replicated |
| | | rs393488 | 9 | 17044971 | A | 0.994 | 0.990 | 0.998 | 1.25E-03 | Replicated |

| | | | | | | | | | |
|---|---|---|---|---|---|---|---|---|---|
| | rs12057031 | 9 | 25235063 | T | 0.995 | 0.991 | 0.999 | 6.92E-03 | NotReplicated |
| | rs11599236 | 10 | 106454672 | C | 0.989 | 0.986 | 0.993 | 5.16E-08 | Replicated |
| | rs537635 | 11 | 88705235 | T | 1.011 | 1.007 | 1.015 | 1.74E-08 | Replicated |
| | rs578174 | 11 | 89959637 | G | 0.994 | 0.990 | 0.997 | 7.41E-04 | Replicated |
| | rs12889665 | 14 | 75234830 | T | 0.987 | 0.983 | 0.991 | 9.88E-12 | Replicated |
| | rs61997596 | 14 | 104511206 | A | 1.005 | 1.001 | 1.009 | 1.29E-02 | NotReplicated |
| | rs11646401 | 16 | 21609978 | G | 1.003 | 0.999 | 1.007 | 1.38E-01 | NotReplicated |
| | rs12967855 | 18 | 35138245 | A | 1.015 | 1.012 | 1.019 | 1.47E-15 | Replicated |

**Supplemental Table S19: Effects of GWAS loci from help-seeking definitions on other definitions of MDD in UKBiobank and psychiatric conditions**

This table shows the genome-wide significant loci (top SNP in 1MB regions) in GWAS minimal phenotyping, help-seeking definitions GPpsy and Psypsy and their effects in the following phenotypes: help-seeking definition GPpsy, CIDI-based definition LifetimeMDD, help-seeking no-MDD definition that specifically exclude MDD symptoms GPNoDep, and other psychiatric conditions SCZ, neuroticism and smoking. For each SNP we show the chromosome (CHR), rsid (SNP), position on the chromosome (BP), and test and minor allele (A1). For each SNP-phenotype association we show the odds ratio (OR, in the case of neuroticism which is a quantitative trait, we show exp(BETA) as OR), the lower and upper bounds of the 95% confidence interval of OR (L95 and U95), and the p-value of association (P). An association is considered “Replicated’ if its p-value is below $3{\cdot}09x10^{-4}$ ($p < 0{\cdot}05$ after multiple testing correction for 162 tests in total) and its direction of effect is the same as that in the GPpsy and Psypsy, where the association is discovered.

| CATEGORY | PHENOTYPE | SNP | CHR | BP | A1 | OR | L95 | U95 | P | REPLICATION |
|---|---|---|---|---|---|---|---|---|---|---|
| **Help seeking** | **GPpsy** | rs301806 | 1 | 8482078 | C | 1.021 | 1.010 | 1.031 | 1.00E-04 | Replicated |
| | | rs11209948 | 1 | 72811904 | G | 0.963 | 0.953 | 0.973 | 1.62E-12 | Replicated |
| | | rs2422321 | 1 | 73293393 | G | 1.021 | 1.010 | 1.032 | 8.96E-05 | Replicated |
| | | rs12065553 | 1 | 80793118 | G | 1.014 | 1.003 | 1.025 | 1.58E-02 | NotReplicated |
| | | rs1518395 | 2 | 58208074 | A | 0.989 | 0.978 | 0.999 | 3.21E-02 | NotReplicated |
| | | rs1656369 | 3 | 158280085 | A | 0.979 | 0.969 | 0.990 | 1.23E-04 | Replicated |
| | | rs10514299 | 5 | 87663610 | T | 1.017 | 1.005 | 1.029 | 4.77E-03 | NotReplicated |
| | | rs454214 | 5 | 88003403 | C | 1.018 | 1.008 | 1.029 | 6.71E-04 | Replicated |
| | | rs4543289 | 5 | 164484948 | T | 0.977 | 0.967 | 0.987 | 6.52E-06 | Replicated |
| | | rs1475120 | 6 | 105389953 | G | 0.981 | 0.971 | 0.991 | 2.71E-04 | Replicated |
| | | rs7044150 | 9 | 2982931 | T | 0.987 | 0.977 | 0.998 | 1.90E-02 | NotReplicated |
| | | rs6476606 | 9 | 37005561 | A | 1.023 | 1.012 | 1.034 | 2.54E-05 | Replicated |
| | | rs10786831 | 10 | 106614571 | A | 1.026 | 1.015 | 1.036 | 1.51E-06 | Replicated |
| | | rs2125716 | 12 | 84941429 | A | 1.018 | 1.006 | 1.030 | 4.03E-03 | NotReplicated |
| | | rs12552 | 13 | 53625781 | A | 1.011 | 1.001 | 1.022 | 3.25E-02 | NotReplicated |
| | | rs8025231 | 15 | 37648402 | C | 1.018 | 1.008 | 1.029 | 5.68E-04 | Replicated |
| | | rs2179744 | 22 | 41621714 | A | 1.015 | 1.004 | 1.027 | 7.78E-03 | NotReplicated |
| **DSM based** | **LifetimeMDD** | rs301806 | 1 | 8482078 | C | 0.998 | 0.973 | 1.023 | 8.44E-01 | NotReplicated |
| | | rs11209948 | 1 | 72811904 | G | 0.977 | 0.952 | 1.002 | 6.88E-02 | NotReplicated |
| | | rs2422321 | 1 | 73293393 | G | 1.038 | 1.012 | 1.065 | 4.52E-03 | NotReplicated |
| | | rs12065553 | 1 | 80793118 | G | 1.032 | 1.004 | 1.060 | 2.37E-02 | NotReplicated |
| | | rs1518395 | 2 | 58208074 | A | 0.990 | 0.965 | 1.016 | 4.63E-01 | NotReplicated |
| | | rs1656369 | 3 | 158280085 | A | 0.978 | 0.952 | 1.004 | 8.95E-02 | NotReplicated |
| | | rs10514299 | 5 | 87663610 | T | 1.008 | 0.979 | 1.037 | 6.03E-01 | NotReplicated |
| | | rs454214 | 5 | 88003403 | C | 1.029 | 1.003 | 1.056 | 2.77E-02 | NotReplicated |
| | | rs4543289 | 5 | 164484948 | T | 0.949 | 0.926 | 0.974 | 5.19E-05 | Replicated |
| | | rs1475120 | 6 | 105389953 | G | 0.987 | 0.962 | 1.012 | 2.92E-01 | NotReplicated |
| | | rs7044150 | 9 | 2982931 | T | 1.006 | 0.980 | 1.032 | 6.77E-01 | NotReplicated |
| | | rs6476606 | 9 | 37005561 | A | 1.021 | 0.995 | 1.048 | 1.22E-01 | NotReplicated |
| | | rs10786831 | 10 | 106614571 | A | 1.024 | 0.999 | 1.051 | 6.34E-02 | NotReplicated |

| | | | | | | | | | | |
|---|---|---|---|---|---|---|---|---|---|---|
| | | rs2125716 | 12 | 84941429 | A | 1.013 | 0.983 | 1.044 | 3.97E-01 | NotReplicated |
| | | rs12552 | 13 | 53625781 | A | 1.042 | 1.016 | 1.068 | 1.62E-03 | Replicated |
| | | rs8025231 | 15 | 37648402 | C | 1.027 | 1.002 | 1.053 | 3.58E-02 | NotReplicated |
| | | rs2179744 | 22 | 41621714 | A | 1.044 | 1.015 | 1.073 | 2.47E-03 | Replicated |
| **No-MDD** | **GPNoDep** | rs301806 | 1 | 8482078 | C | 0.985 | 0.953 | 1.018 | 3.64E-01 | NotReplicated |
| | | rs11209948 | 1 | 72811904 | G | 0.984 | 0.952 | 1.017 | 3.32E-01 | NotReplicated |
| | | rs2422321 | 1 | 73293393 | G | 1.014 | 0.981 | 1.048 | 4.16E-01 | NotReplicated |
| | | rs12065553 | 1 | 80793118 | G | 0.980 | 0.946 | 1.015 | 2.65E-01 | NotReplicated |
| | | rs1518395 | 2 | 58208074 | A | 0.999 | 0.966 | 1.032 | 9.38E-01 | NotReplicated |
| | | rs1656369 | 3 | 158280085 | A | 0.984 | 0.951 | 1.018 | 3.61E-01 | NotReplicated |
| | | rs10514299 | 5 | 87663610 | T | 1.040 | 1.002 | 1.079 | 4.05E-02 | NotReplicated |
| | | rs454214 | 5 | 88003403 | C | 1.020 | 0.986 | 1.054 | 2.49E-01 | NotReplicated |
| | | rs4543289 | 5 | 164484948 | T | 0.958 | 0.928 | 0.990 | 9.89E-03 | NotReplicated |
| | | rs1475120 | 6 | 105389953 | G | 1.022 | 0.989 | 1.056 | 1.89E-01 | NotReplicated |
| | | rs7044150 | 9 | 2982931 | T | 1.005 | 0.972 | 1.040 | 7.61E-01 | NotReplicated |
| | | rs6476606 | 9 | 37005561 | A | 0.999 | 0.966 | 1.034 | 9.65E-01 | NotReplicated |
| | | rs10786831 | 10 | 106614571 | A | 1.050 | 1.016 | 1.085 | 4.02E-03 | NotReplicated |
| | | rs2125716 | 12 | 84941429 | A | 1.045 | 1.005 | 1.086 | 2.57E-02 | NotReplicated |
| | | rs12552 | 13 | 53625781 | A | 1.006 | 0.974 | 1.040 | 7.20E-01 | NotReplicated |
| | | rs8025231 | 15 | 37648402 | C | 1.023 | 0.990 | 1.057 | 1.74E-01 | NotReplicated |
| | | rs2179744 | 22 | 41621714 | A | 1.012 | 0.976 | 1.049 | 5.18E-01 | NotReplicated |
| **Other condition** | **SCZ** | rs301806 | 1 | 8482078 | C | 0.946 | 0.925 | 0.967 | 1.38E-06 | NotReplicated |
| | | rs11209948 | 1 | 72811904 | G | 0.980 | 0.958 | 1.002 | 7.77E-02 | NotReplicated |
| | | rs2422321 | 1 | 73293393 | G | 1.059 | 1.037 | 1.080 | 2.18E-08 | Replicated |
| | | rs12065553 | 1 | 80793118 | G | 1.016 | 0.993 | 1.039 | 1.77E-01 | NotReplicated |
| | | rs1518395 | 2 | 58208074 | A | 0.942 | 0.921 | 0.963 | 3.43E-08 | Replicated |
| | | rs1656369 | 3 | 158280085 | A | 1.000 | 0.977 | 1.022 | 9.72E-01 | NotReplicated |
| | | rs10514299 | 5 | 87663610 | T | 0.964 | 0.940 | 0.988 | 2.75E-03 | NotReplicated |
| | | rs454214 | 5 | 88003403 | C | 0.973 | 0.952 | 0.993 | 1.09E-02 | NotReplicated |
| | | rs4543289 | 5 | 164484948 | T | 0.995 | 0.974 | 1.016 | 6.67E-01 | NotReplicated |
| | | rs1475120 | 6 | 105389953 | G | 1.049 | 1.029 | 1.070 | 1.87E-06 | NotReplicated |

| | | | | | | | | | | |
|---|---|---|---|---|---|---|---|---|---|---|
| | | rs7044150 | 9 | 2982931 | T | 0.985 | 0.963 | 1.007 | 1.74E-01 | NotReplicated |
| | | rs6476606 | 9 | 37005561 | A | 0.978 | 0.956 | 0.999 | 4.24E-02 | NotReplicated |
| | | rs10786831 | 10 | 106614571 | A | 1.047 | 1.025 | 1.068 | 2.34E-05 | Replicated |
| | | rs2125716 | 12 | 84941429 | A | 0.991 | 0.965 | 1.016 | 4.72E-01 | NotReplicated |
| | | rs12552 | 13 | 53625781 | A | 0.993 | 0.972 | 1.014 | 5.22E-01 | NotReplicated |
| | | rs8025231 | 15 | 37648402 | C | 1.003 | 0.982 | 1.024 | 7.63E-01 | NotReplicated |
| | | rs2179744 | 22 | 41621714 | A | 1.064 | 1.041 | 1.087 | 8.31E-08 | Replicated |
| **Other condition** | **Smoking** | rs301806 | 1 | 8482078 | C | 0.989 | 0.979 | 0.999 | 2.59E-02 | NotReplicated |
| | | rs11209948 | 1 | 72811904 | G | 0.994 | 0.984 | 1.004 | 2.47E-01 | NotReplicated |
| | | rs2422321 | 1 | 73293393 | G | 1.012 | 1.001 | 1.022 | 2.44E-02 | NotReplicated |
| | | rs12065553 | 1 | 80793118 | G | 1.024 | 1.013 | 1.035 | 1.99E-05 | Replicated |
| | | rs1518395 | 2 | 58208074 | A | 0.989 | 0.979 | 0.999 | 2.85E-02 | NotReplicated |
| | | rs1656369 | 3 | 158280085 | A | 0.981 | 0.971 | 0.991 | 3.07E-04 | Replicated |
| | | rs10514299 | 5 | 87663610 | T | 0.974 | 0.963 | 0.985 | 3.95E-06 | NotReplicated |
| | | rs454214 | 5 | 88003403 | C | 0.995 | 0.985 | 1.005 | 3.06E-01 | NotReplicated |
| | | rs4543289 | 5 | 164484948 | T | 0.990 | 0.980 | 0.999 | 3.74E-02 | NotReplicated |
| | | rs1475120 | 6 | 105389953 | G | 0.999 | 0.989 | 1.008 | 7.76E-01 | NotReplicated |
| | | rs7044150 | 9 | 2982931 | T | 0.994 | 0.984 | 1.004 | 2.49E-01 | NotReplicated |
| | | rs6476606 | 9 | 37005561 | A | 0.995 | 0.985 | 1.005 | 3.16E-01 | NotReplicated |
| | | rs10786831 | 10 | 106614571 | A | 1.006 | 0.996 | 1.016 | 2.40E-01 | NotReplicated |
| | | rs2125716 | 12 | 84941429 | A | 1.019 | 1.007 | 1.031 | 1.93E-03 | Replicated |
| | | rs12552 | 13 | 53625781 | A | 1.018 | 1.008 | 1.028 | 5.80E-04 | Replicated |
| | | rs8025231 | 15 | 37648402 | C | 1.004 | 0.995 | 1.014 | 3.78E-01 | NotReplicated |
| | | rs2179744 | 22 | 41621714 | A | 1.000 | 0.990 | 1.011 | 9.40E-01 | NotReplicated |
| **Other condition** | **Neuroticism** | rs301806 | 1 | 8482078 | C | 1.009 | 1.005 | 1.013 | 4.26E-06 | Replicated |
| | | rs11209948 | 1 | 72811904 | G | 0.997 | 0.994 | 1.001 | 1.56E-01 | NotReplicated |
| | | rs2422321 | 1 | 73293393 | G | 1.003 | 0.999 | 1.007 | 1.37E-01 | NotReplicated |
| | | rs12065553 | 1 | 80793118 | G | 1.005 | 1.002 | 1.009 | 5.13E-03 | NotReplicated |
| | | rs1518395 | 2 | 58208074 | A | 0.991 | 0.987 | 0.995 | 1.38E-06 | Replicated |
| | | rs1656369 | 3 | 158280085 | A | 0.995 | 0.991 | 0.999 | 6.42E-03 | NotReplicated |
| | | rs10514299 | 5 | 87663610 | T | 1.005 | 1.001 | 1.009 | 7.18E-03 | NotReplicated |

| | | | | | | | | | |
|---|---|---|---|---|---|---|---|---|---|
| | rs454214 | 5 | 88003403 | C | 1.004 | 1.001 | 1.008 | 2.09E-02 | NotReplicated |
| | rs4543289 | 5 | 164484948 | T | 0.992 | 0.988 | 0.995 | 8.95E-06 | Replicated |
| | rs1475120 | 6 | 105389953 | G | 0.994 | 0.990 | 0.997 | 7.84E-04 | Replicated |
| | rs7044150 | 9 | 2982931 | T | 0.995 | 0.991 | 0.999 | 6.36E-03 | NotReplicated |
| | rs6476606 | 9 | 37005561 | A | 1.005 | 1.001 | 1.009 | 6.82E-03 | NotReplicated |
| | rs10786831 | 10 | 106614571 | A | 1.007 | 1.003 | 1.011 | 4.39E-04 | Replicated |
| | rs2125716 | 12 | 84941429 | A | 1.002 | 0.999 | 1.006 | 2.30E-01 | NotReplicated |
| | rs12552 | 13 | 53625781 | A | 1.002 | 0.999 | 1.006 | 2.25E-01 | NotReplicated |
| | rs8025231 | 15 | 37648402 | C | 1.003 | 0.999 | 1.006 | 1.67E-01 | NotReplicated |
| | rs2179744 | 22 | 41621714 | A | 1.011 | 1.007 | 1.015 | 2.24E-08 | Replicated |

**Supplemental Table S20: Effects of GWAS loci from minimal phenotyping definition of MDD in 23andMe on definitions of MDD in UKBiobank and psychiatric conditions**

This table shows the genome-wide significant loci (top SNP in 1MB regions) in GWAS minimal phenotyping definitions of MDD in 23andMe and their effects in the following phenotypes: help-seeking definition GPpsy, CIDI-based definition LifetimeMDD, help-seeking no-MDD definition that specifically exclude MDD symptoms GPNoDep, and other psychiatric conditions SCZ, neuroticism and smoking. For each SNP we show the chromosome (CHR), rsid (SNP), position on the chromosome (BP), and test and minor allele (A1). For each SNP-phenotype association we show the odds ratio (OR, in the case of neuroticism which is a quantitative trait, we show exp(BETA) as OR), the lower and upper bounds of the 95% confidence interval of OR (L95 and U95), and the p-value of association (P). An association is considered “Replicated’ if its p-value is below $4·90x10^{-4}$ ($p < 0·05$ after multiple testing correction for 102 tests in total) and its direction of effect is the same as that in 23andMe, where the association is discovered.

| PGC COHORTS | | | | | PRS Analysis | | DSM-MDD Criteria A Symptoms | | |
|---|---|---|---|---|---|---|---|---|---|
| **Name** | **Cohort** | **Ncases** | **Ncontrols** | **N** | **Used** | **Reason Not Used** | **Symptoms Available** | **DSM-MDD Cases** | **% DSM-MDD Cases** |
| BOMA | boma | 586 | 1062 | 1648 | 1 | | Yes | 541 | 92.48 |
| CoFams | cof3 | 120 | 126 | 246 | 0 | N < 500 | NA | | |
| PsyCoLaus | col3 | 507 | 1445 | 1952 | 1 | | Yes | 485 | 95.85 |
| Edinburgh | edi2 | 372 | 285 | 657 | 1 | | No | | |
| GenRED1 | gens | 1019 | 1344 | 2363 | 1 | | Yes | 1002 | 98.23 |
| GenPod/Newmeds | gep3 | 482 | 2836 | 3318 | 1 | | Yes | 233 | 50.43 |
| Depression Genetic Network (DGN) | grdg | 471 | 470 | 941 | 1 | | Yes | 461 | 97.05 |
| GenRED2 | grnd | 830 | 474 | 1304 | 1 | | Yes | 808 | 98.54 |
| GSK/Max Planck Inst Psychiatry | gsk2 | 880 | 861 | 1741 | 0 | PERMISSION | NA | | |
| i2b2 | i2b3 | 806 | 1067 | 1873 | 1 | | No | | |
| Janssen | jjp2 | 466 | 1380 | 1846 | 0 | PERMISSION | NA | | |
| MPIP/MARS old | mmi2 | 584 | 517 | 1101 | 1 | | Yes | 291 | 49.83 |
| MPIP/MARS new | mmo4 | 264 | 371 | 635 | 1 | | Yes | 113 | 42.80 |
| NESDA | nes1 | 1494 | 1602 | 3096 | 1 | | Yes | 1294 | 94.11 |
| Pfizer | pfm2 | 281 | 820 | 1101 | 0 | PERMISSION | NA | | |
| QIMR | qi3c | 864 | 579 | 1443 | 1 | | Yes | 556 | 95.53 |
| QIMR | qi6c | 499 | 590 | 1089 | 1 | | Yes | 465 | 93.19 |
| QIMR | qio2 | 565 | 526 | 1091 | 1 | | Yes | 531 | 93.98 |
| RADIANT-UK | rad3 | 1872 | 1528 | 3400 | 1 | | Yes | 1552 | 88.08 |
| RADIANT-GER | rage | 322 | 227 | 549 | 1 | | Yes | 297 | 92.24 |
| RADIANT-Irish | rai2 | 109 | 340 | 449 | 0 | N < 500 | NA | | |
| RADIANT-US | rau2 | 223 | 378 | 601 | 1 | | Yes | 217 | 97.75 |
| RADIANT-DE | rde4 | 133 | 516 | 649 | 1 | | Yes | 124 | 93.94 |
| Roche | roc3 | 271 | 92 | 363 | 0 | PERMISSION | NA | | |
| Rotterdam | rot4 | 241 | 1028 | 1269 | 1 | | Yes | 113 | 50.67 |

| | | | | | | | | | |
|---|---|---|---|---|---|---|---|---|---|
| Ship0 | shp0 | 366 | 1087 | 1453 | 1 | | Yes | 337 | 90.35 |
| ShipTrend | shpt | 163 | 484 | 647 | 1 | | No | | |
| STAR*D | stm2 | 936 | 934 | 1870 | 1 | | Yes | 817 | 87.57 |
| TwinGene | twg2 | 1097 | 2663 | 3760 | 1 | | Yes | 806 | 70.83 |

**Supplemental Table S21: PGC cohorts for out-of-sample predictions of MDD**

This table shows the 29 cohorts in PGC29, as shown in Supplemental Table 3 of Wray et al 2018[32]. We obtained access to 22 of the 29 cohorts through the MDD Working Group of the PGC, and used 20 out of the 22 in our out-of-sample prediction analysis (Used), removing 2 because of their small sample sizes (<500 samples). We indicated the reasons for each of the cohorts we did not use (Reason Not Used). Of note, rad3 is removed from all our analyses post-hoc due to its likely containing individuals that are relatives of individuals in UKBiobank, due to the much higher predictive power in this cohort as compared to other cohorts (see Supplemental Table S21 and Supplemental Figure S13). For each of the cohorts, we indicated their cohort name (Name), abbreviation (Cohort), number of individuals indicated as MDD cases (Ncases), number of individuals indicated as controls (Ncontrols), and total sample size (N). Further, we indicate whether individual level symptom endorsement of DSM-V criterion A symptoms are recorded and available for each cohort (Symptoms Available), and the number of DSM-MDD cases we were able to derive from the individual level symptoms data (DSM-MDD Cases). Using this we calculated the percentage of individuals indicated in the cohort as MDD cases (Ncases) that fulfilled DSM-V criterion A (% DSM-MDD Cases).

| PGC Cohorts | | | Downsampled GPpsy | | | GPpsy | | |
|---|---|---|---|---|---|---|---|---|
| | | | P value threshold = 0.05 | | | P value threshold = 0.05 | | |
| **Cohort** | **Nsamples** | **%Cases DSM-MDD** | **Nagelkerke's r2 (P)** | **AUC** | **h2_Liab (SE)** | **Nagelkerke's r2 (P)** | **AUC** | **h2_Liab (SE)** |
| boma | 1648 | 92.48 | 1.72E-03 (1.56E-01) | 0.51 | 1.63E-03 (2.22E-03) | 1.37E-02 (5.87E-05) | 0.56 | 1.31E-02 (6.26E-03) |
| col3 | 1952 | 95.85 | 2.07E-06 (9.58E-01) | 0.50 | 2.20E-06 (8.45E-05) | 1.96E-02 (3.04E-07) | 0.58 | 2.09E-02 (8.04E-03) |
| edi2 | 657 | NA | 2.04E-04 (7.59E-01) | 0.50 | 1.85E-04 (8.55E-04) | 3.48E-02 (5.39E-05) | 0.59 | 3.18E-02 (1.49E-02) |
| gens | 2363 | 98.24 | 2.01E-03 (6.10E-02) | 0.52 | 1.82E-03 (1.93E-03) | 3.15E-02 (8.70E-14) | 0.58 | 2.88E-02 (7.52E-03) |
| gep3 | 3318 | 50.43 | 1.13E-03 (1.54E-01) | 0.52 | 1.54E-03 (2.13E-03) | 9.56E-03 (3.24E-05) | 0.56 | 1.30E-02 (6.24E-03) |
| grdg | 941 | NA | 1.00E-05 (9.33E-01) | 0.50 | 8.98E-06 (2.15E-04) | 1.85E-02 (2.95E-04) | 0.56 | 1.67E-02 (9.06E-03) |
| grnd | 1304 | 98.54 | 2.03E-04 (6.62E-01) | 0.51 | 1.92E-04 (8.69E-04) | 2.35E-02 (2.26E-06) | 0.58 | 2.23E-02 (9.23E-03) |
| i2b3 | 1873 | NA | 8.31E-04 (2.89E-01) | 0.51 | 7.55E-04 (1.40E-03) | 3.55E-03 (2.83E-02) | 0.53 | 3.23E-03 (2.86E-03) |
| mmi2 | 1101 | 49.83 | 1.32E-04 (7.42E-01) | 0.51 | 1.18E-04 (7.01E-04) | 1.78E-02 (1.23E-04) | 0.56 | 1.61E-02 (8.26E-03) |
| mmo4 | 635 | 42.8 | 2.14E-03 (3.66E-01) | 0.49 | 1.96E-03 (3.28E-03) | 5.98E-03 (1.31E-01) | 0.53 | 5.47E-03 (5.75E-03) |
| nes1 | 3096 | 94.11 | 2.91E-03 (9.44E-03) | 0.53 | 2.61E-03 (2.00E-03) | 1.66E-02 (4.74E-10) | 0.56 | 1.50E-02 (4.75E-03) |
| qi3c | 1443 | 95.53 | 4.06E-04 (5.90E-01) | 0.51 | 3.75E-04 (1.37E-03) | 1.56E-02 (8.37E-04) | 0.56 | 1.44E-02 (8.48E-03) |
| qi6c | 1089 | 93.19 | 8.48E-03 (8.52E-03) | 0.55 | 7.66E-03 (5.77E-03) | 1.83E-02 (1.11E-04) | 0.56 | 1.65E-02 (8.43E-03) |
| qio2 | 1091 | 93.98 | 4.61E-03 (5.23E-02) | 0.54 | 4.14E-03 (4.23E-03) | 5.37E-03 (3.62E-02) | 0.54 | 4.83E-03 (4.58E-03) |
| rad3 | 3400 | 88.08 | 6.20E-03 (1.19E-04) | 0.54 | 5.61E-03 (2.88E-03) | 5.46E-02 (9.97E-31) | 0.61 | 4.99E-02 (8.32E-03) |
| rage | 549 | 92.24 | 4.38E-03 (1.90E-01) | 0.54 | 4.01E-03 (5.59E-03) | 5.84E-03 (1.30E-01) | 0.54 | 5.36E-03 (6.64E-03) |
| rau2 | 601 | 97.75 | 1.24E-03 (4.62E-01) | 0.48 | 1.16E-03 (3.04E-03) | 3.30E-03 (2.30E-01) | 0.54 | 3.10E-03 (5.11E-03) |
| rde4 | 649 | 93.94 | 1.17E-03 (4.88E-01) | 0.52 | 1.37E-03 (3.85E-03) | 1.05E-02 (3.71E-02) | 0.55 | 1.23E-02 (1.18E-02) |
| rot4 | 1269 | 50.67 | 1.12E-03 (3.48E-01) | 0.52 | 1.35E-03 (2.86E-03) | 5.37E-03 (3.94E-02) | 0.54 | 6.50E-03 (6.26E-03) |
| shp0 | 1453 | 90.35 | 1.05E-06 (9.74E-01) | 0.50 | 1.13E-06 (6.99E-05) | 7.18E-03 (7.88E-03) | 0.55 | 7.72E-03 (5.77E-03) |
| shpt | 647 | NA | 6.00E-04 (6.10E-01) | 0.49 | 6.44E-04 (2.53E-03) | 1.46E-02 (1.16E-02) | 0.56 | 1.57E-02 (1.23E-02) |
| stm2 | 1870 | 87.57 | 2.39E-04 (5.65E-01) | 0.51 | 2.14E-04 (7.34E-04) | 4.08E-03 (1.73E-02) | 0.53 | 3.66E-03 (3.03E-03) |
| twg2 | 3760 | 70.83 | 6.39E-04 (1.95E-01) | 0.51 | 6.48E-04 (1.00E-03) | 1.56E-02 (1.30E-10) | 0.56 | 1.59E-02 (4.91E-03) |
| **PGC Cohorts** | | | **Downsampled DepAll** | | | **DepAll** | | |
| | | | **P value threshold = 0.05** | | | **P value threshold = 0.05** | | |
| **Cohort** | **Nsamples** | **%Cases DSM-MDD** | **Nagelkerke's r2 (P)** | **AUC** | **h2_Liab (SE)** | **Nagelkerke's r2 (P)** | **AUC** | **h2_Liab (SE)** |

| | | | | | | | | |
|---|---|---|---|---|---|---|---|---|
| boma | 1648 | 92.48 | 4.24E-03 (2.57E-02) | 0.54 | 4.04E-03 (3.45E-03) | 8.05E-03 (2.11E-03) | 0.55 | 7.66E-03 (4.86E-03) |
| col3 | 1952 | 95.85 | 1.05E-03 (6.61E-01) | 0.53 | 9.43E-04 (4.19E-03) | 3.20E-03 (3.92E-02) | 0.53 | 3.39E-03 (3.28E-03) |
| edi2 | 657 | NA | 9.68E-04 (5.03E-01) | 0.51 | 8.79E-04 (2.55E-03) | 1.67E-03 (3.79E-01) | 0.52 | 1.52E-03 (3.06E-03) |
| gens | 2363 | 98.24 | 2.80E-03 (2.68E-02) | 0.53 | 2.55E-03 (2.27E-03) | 7.63E-03 (2.55E-04) | 0.54 | 6.95E-03 (3.75E-03) |
| gep3 | 3318 | 50.43 | 7.78E-04 (2.36E-01) | 0.52 | 1.06E-03 (1.79E-03) | 3.93E-03 (7.73E-03) | 0.54 | 5.36E-03 (4.00E-03) |
| grdg | 941 | NA | 1.07E-03 (3.86E-01) | 0.51 | 9.60E-04 (2.20E-03) | 7.14E-03 (2.50E-02) | 0.55 | 6.42E-03 (5.66E-03) |
| grnd | 1304 | 98.54 | 2.27E-04 (6.43E-01) | 0.50 | 2.15E-04 (9.15E-04) | 3.80E-03 (5.82E-02) | 0.53 | 3.59E-03 (3.74E-03) |
| i2b3 | 1873 | NA | 4.84E-03 (1.05E-02) | 0.53 | 4.40E-03 (3.30E-03) | 5.95E-03 (4.52E-03) | 0.54 | 5.42E-03 (3.65E-03) |
| mmi2 | 1101 | 49.83 | 2.03E-04 (6.83E-01) | 0.51 | 1.83E-04 (8.85E-04) | 1.39E-03 (2.85E-01) | 0.52 | 1.25E-03 (2.33E-03) |
| mmo4 | 635 | 42.8 | 1.41E-02 (2.00E-02) | 0.55 | 1.30E-02 (8.55E-03) | 2.16E-02 (3.95E-03) | 0.54 | 1.99E-02 (1.17E-02) |
| nes1 | 3096 | 94.11 | 4.40E-04 (3.13E-01) | 0.52 | 3.95E-04 (7.79E-04) | 2.86E-03 (1.01E-02) | 0.52 | 2.57E-03 (1.98E-03) |
| qi3c | 1443 | 95.53 | 2.77E-03 (1.60E-01) | 0.52 | 2.55E-03 (3.58E-03) | 5.44E-03 (4.86E-02) | 0.54 | 5.03E-03 (5.05E-03) |
| qi6c | 1089 | 93.19 | 1.09E-03 (3.46E-01) | 0.51 | 9.86E-04 (2.08E-03) | 1.04E-02 (3.62E-03) | 0.55 | 9.37E-03 (6.38E-03) |
| qio2 | 1091 | 93.98 | 5.20E-05 (8.37E-01) | 0.51 | 4.67E-05 (4.43E-04) | 9.18E-04 (3.87E-01) | 0.52 | 8.25E-04 (1.88E-03) |
| rad3 | 3400 | 88.08 | 4.98E-03 (5.65E-04) | 0.53 | 4.50E-03 (2.59E-03) | 1.85E-02 (2.77E-11) | 0.57 | 1.68E-02 (4.95E-03) |
| rage | 549 | 92.24 | 4.18E-04 (6.86E-01) | 0.52 | 3.82E-04 (2.16E-03) | 3.36E-03 (2.50E-01) | 0.52 | 3.08E-03 (4.91E-03) |
| rau2 | 601 | 97.75 | 9.96E-04 (5.09E-01) | 0.51 | 9.35E-04 (2.78E-03) | 2.26E-03 (3.20E-01) | 0.52 | 2.13E-03 (4.27E-03) |
| rde4 | 649 | 93.94 | 2.13E-05 (9.25E-01) | 0.51 | 2.49E-05 (5.43E-04) | 3.33E-03 (2.42E-01) | 0.53 | 3.89E-03 (6.57E-03) |
| rot4 | 1269 | 50.67 | 4.97E-04 (5.31E-01) | 0.51 | 6.01E-04 (1.92E-03) | 1.31E-04 (7.48E-01) | 0.51 | 1.58E-04 (9.71E-04) |
| shp0 | 1453 | 90.35 | 8.26E-05 (7.76E-01) | 0.50 | 8.86E-05 (6.31E-04) | 2.98E-05 (8.64E-01) | 0.50 | 3.20E-05 (3.85E-04) |
| shpt | 647 | NA | 8.40E-04 (5.46E-01) | 0.52 | 9.02E-04 (2.93E-03) | 9.25E-03 (4.47E-02) | 0.55 | 9.94E-03 (9.72E-03) |
| stm2 | 1870 | 87.57 | 7.34E-04 (3.13E-01) | 0.52 | 6.59E-04 (1.27E-03) | 5.26E-03 (6.84E-03) | 0.54 | 4.72E-03 (3.43E-03) |
| twg2 | 3760 | 70.83 | 3.08E-03 (4.41E-03) | 0.53 | 3.13E-03 (2.19E-03) | 4.01E-03 (1.17E-03) | 0.53 | 4.07E-03 (2.50E-03) |

| **PGC Cohorts** | | | **Downsampled SelfRepDep** | | | **SelfRepDep** | | |
|---|---|---|---|---|---|---|---|---|
| | | | **P value threshold = 0.05** | | | **P value threshold = 0.05** | | |
| **Cohort** | **Nsamples** | **%Cases DSM-MDD** | **Nagelkerke's r2 (P)** | **AUC** | **h2_Liab (SE)** | **Nagelkerke's r2 (P)** | **AUC** | **h2_Liab (SE)** |
| boma | 1648 | 92.48 | 1.25E-03 (2.26E-01) | 0.53 | 1.19E-03 (1.90E-03) | 8.72E-03 (1.38E-03) | 0.55 | 8.30E-03 (5.02E-03) |
| col3 | 1952 | 95.85 | 5.61E-04 (7.49E-01) | 0.52 | 5.03E-04 (3.26E-03) | 2.74E-03 (5.64E-02) | 0.48 | 2.90E-03 (3.04E-03) |
| edi2 | 657 | NA | 1.31E-03 (4.36E-01) | 0.50 | 1.19E-03 (2.80E-03) | 5.48E-03 (1.11E-01) | 0.54 | 4.98E-03 (5.81E-03) |

| | | | | | | | | |
|---|---|---|---|---|---|---|---|---|
| gens | 2363 | 98.24 | 3.39E-03 (1.49E-02) | 0.54 | 3.08E-03 (2.50E-03) | 8.10E-03 (1.65E-04) | 0.55 | 7.37E-03 (3.87E-03) |
| gep3 | 3318 | 50.43 | 2.71E-03 (2.70E-02) | 0.53 | 3.70E-03 (3.34E-03) | 1.60E-03 (8.92E-02) | 0.53 | 2.18E-03 (2.57E-03) |
| grdg | 941 | NA | 7.72E-05 (8.16E-01) | 0.50 | 6.92E-05 (5.99E-04) | 9.44E-03 (9.92E-03) | 0.55 | 8.49E-03 (6.47E-03) |
| grnd | 1304 | 98.54 | 4.78E-04 (5.02E-01) | 0.52 | 4.51E-04 (1.34E-03) | 2.72E-03 (1.09E-01) | 0.54 | 2.56E-03 (3.18E-03) |
| i2b3 | 1873 | NA | 7.24E-04 (3.22E-01) | 0.52 | 6.58E-04 (1.32E-03) | 1.31E-03 (1.84E-01) | 0.52 | 1.19E-03 (1.75E-03) |
| mmi2 | 1101 | 49.83 | 5.22E-06 (9.48E-01) | 0.51 | 4.70E-06 (1.42E-04) | 2.14E-03 (1.85E-01) | 0.48 | 1.92E-03 (2.88E-03) |
| mmo4 | 635 | 42.8 | 5.45E-03 (1.49E-01) | 0.53 | 4.99E-03 (5.50E-03) | 1.11E-02 (3.92E-02) | 0.55 | 1.02E-02 (8.07E-03) |
| nes1 | 3096 | 94.11 | 2.37E-03 (1.91E-02) | 0.53 | 2.13E-03 (1.81E-03) | 4.55E-03 (1.16E-03) | 0.53 | 4.09E-03 (2.50E-03) |
| qi3c | 1443 | 95.53 | 3.28E-05 (8.78E-01) | 0.50 | 3.03E-05 (3.81E-04) | 5.45E-04 (5.33E-01) | 0.51 | 5.03E-04 (1.62E-03) |
| qi6c | 1089 | 93.19 | 2.81E-03 (1.30E-01) | 0.52 | 2.54E-03 (3.33E-03) | 7.98E-04 (4.20E-01) | 0.52 | 7.20E-04 (1.78E-03) |
| qio2 | 1091 | 93.98 | 7.24E-07 (9.81E-01) | 0.51 | 6.50E-07 (9.02E-05) | 3.77E-06 (9.56E-01) | 0.50 | 3.38E-06 (9.47E-05) |
| rad3 | 3400 | 88.08 | 4.96E-03 (5.79E-04) | 0.54 | 4.49E-03 (2.58E-03) | 1.69E-02 (1.89E-10) | 0.56 | 1.53E-02 (4.73E-03) |
| rage | 549 | 92.24 | 3.41E-03 (2.47E-01) | 0.54 | 3.12E-03 (5.07E-03) | 1.58E-05 (9.37E-01) | 0.51 | 1.45E-05 (3.36E-04) |
| rau2 | 601 | 97.75 | 3.52E-03 (2.15E-01) | 0.52 | 3.30E-03 (5.18E-03) | 9.16E-05 (8.41E-01) | 0.51 | 8.60E-05 (8.66E-04) |
| rde4 | 649 | 93.94 | 5.59E-04 (6.32E-01) | 0.50 | 6.54E-04 (2.96E-03) | 4.27E-04 (6.75E-01) | 0.53 | 4.99E-04 (2.34E-03) |
| rot4 | 1269 | 50.67 | 7.01E-03 (1.86E-02) | 0.54 | 8.48E-03 (7.17E-03) | 3.32E-03 (1.05E-01) | 0.53 | 4.02E-03 (4.94E-03) |
| shp0 | 1453 | 90.35 | 1.13E-03 (2.92E-01) | 0.52 | 1.21E-03 (2.28E-03) | 5.86E-04 (4.48E-01) | 0.51 | 6.29E-04 (1.65E-03) |
| shpt | 647 | NA | 3.38E-03 (2.26E-01) | 0.53 | 3.63E-03 (5.86E-03) | 5.95E-03 (1.08E-01) | 0.53 | 6.39E-03 (7.84E-03) |
| stm2 | 1870 | 87.57 | 3.48E-03 (2.79E-02) | 0.53 | 3.12E-03 (2.81E-03) | 3.68E-03 (2.38E-02) | 0.53 | 3.30E-03 (2.87E-03) |
| twg2 | 3760 | 70.83 | 2.13E-03 (1.80E-02) | 0.53 | 2.16E-03 (1.82E-03) | 8.91E-03 (1.27E-06) | 0.55 | 9.06E-03 (3.71E-03) |

| **PGC Cohorts** | | | **Downsampled ICD10Dep** | | | **ICD10Dep** | | |
|---|---|---|---|---|---|---|---|---|
| | | | **P value threshold = 0.05** | | | **P value threshold = 0.05** | | |
| **Cohort** | **Nsamples** | **%Cases DSM-MDD** | **Nagelkerke's r2 (P)** | **AUC** | **h2_Liab (SE)** | **Nagelkerke's r2 (P)** | **AUC** | **h2_Liab (SE)** |
| boma | 1648 | 92.48 | 6.89E-04 (3.69E-01) | 0.49 | 6.54E-04 (1.35E-03) | 3.96E-03 (3.13E-02) | 0.53 | 3.76E-03 (3.37E-03) |
| col3 | 1952 | 95.85 | 6.49E-04 (7.30E-01) | 0.51 | 5.82E-04 (3.68E-03) | 2.83E-05 (8.46E-01) | 0.50 | 3.00E-05 (3.12E-04) |
| edi2 | 657 | NA | 9.01E-03 (4.08E-02) | 0.53 | 8.19E-03 (7.94E-03) | 1.42E-02 (1.03E-02) | 0.54 | 1.29E-02 (9.79E-03) |
| gens | 2363 | 98.24 | 1.49E-03 (1.06E-01) | 0.52 | 1.36E-03 (1.66E-03) | 5.04E-03 (2.97E-03) | 0.53 | 4.59E-03 (3.05E-03) |
| gep3 | 3318 | 50.43 | 1.46E-03 (1.05E-01) | 0.53 | 1.99E-03 (2.43E-03) | 5.39E-04 (3.25E-01) | 0.52 | 7.34E-04 (1.48E-03) |
| grdg | 941 | NA | 7.75E-03 (1.95E-02) | 0.55 | 6.97E-03 (5.88E-03) | 1.39E-02 (1.76E-03) | 0.55 | 1.25E-02 (7.84E-03) |

| | | | | | | | | |
|---|---|---|---|---|---|---|---|---|
| grnd | 1304 | 98.54 | 7.46E-04 (4.02E-01) | 0.51 | 7.04E-04 (1.65E-03) | 4.37E-03 (4.22E-02) | 0.53 | 4.13E-03 (4.03E-03) |
| i2b3 | 1873 | NA | 5.51E-03 (6.32E-03) | 0.53 | 5.01E-03 (3.53E-03) | 1.01E-02 (2.08E-04) | 0.55 | 9.24E-03 (4.80E-03) |
| mmi2 | 1101 | 49.83 | 2.35E-03 (1.64E-01) | 0.52 | 2.11E-03 (3.02E-03) | 5.11E-03 (4.03E-02) | 0.53 | 4.60E-03 (4.44E-03) |
| mmo4 | 635 | 42.8 | 6.52E-03 (1.14E-01) | 0.52 | 5.97E-03 (6.17E-03) | 7.48E-03 (9.08E-02) | 0.52 | 6.85E-03 (6.63E-03) |
| nes1 | 3096 | 94.11 | 1.59E-03 (5.53E-02) | 0.52 | 1.42E-03 (1.48E-03) | 2.59E-03 (1.43E-02) | 0.53 | 2.33E-03 (1.89E-03) |
| qi3c | 1443 | 95.53 | 4.67E-03 (6.78E-02) | 0.53 | 4.31E-03 (4.67E-03) | 5.48E-04 (5.32E-01) | 0.50 | 5.06E-04 (1.59E-03) |
| qi6c | 1089 | 93.19 | 1.21E-04 (7.54E-01) | 0.50 | 1.09E-04 (6.96E-04) | 5.75E-04 (4.94E-01) | 0.51 | 5.18E-04 (1.51E-03) |
| qio2 | 1091 | 93.98 | 1.49E-05 (9.12E-01) | 0.50 | 1.34E-05 (2.66E-04) | 2.21E-03 (1.80E-01) | 0.53 | 1.98E-03 (2.94E-03) |
| rad3 | 3400 | 88.08 | 6.04E-03 (1.47E-04) | 0.54 | 5.46E-03 (2.84E-03) | 1.08E-02 (3.59E-07) | 0.55 | 9.80E-03 (3.79E-03) |
| rage | 549 | 92.24 | 1.22E-03 (4.89E-01) | 0.54 | 1.12E-03 (3.20E-03) | 5.92E-03 (1.27E-01) | 0.55 | 5.43E-03 (6.97E-03) |
| rau2 | 601 | 97.75 | 4.74E-03 (1.50E-01) | 0.53 | 4.45E-03 (6.04E-03) | 4.25E-06 (9.66E-01) | 0.50 | 3.99E-06 (1.72E-04) |
| rde4 | 649 | 93.94 | 9.95E-05 (8.40E-01) | 0.50 | 1.16E-04 (1.07E-03) | 2.65E-04 (7.41E-01) | 0.50 | 3.10E-04 (1.95E-03) |
| rot4 | 1269 | 50.67 | 4.07E-04 (5.71E-01) | 0.52 | 4.91E-04 (1.73E-03) | 6.80E-06 (9.42E-01) | 0.50 | 8.21E-06 (2.18E-04) |
| shp0 | 1453 | 90.35 | 2.72E-03 (1.02E-01) | 0.53 | 2.92E-03 (3.55E-03) | 2.38E-03 (1.26E-01) | 0.53 | 2.56E-03 (3.32E-03) |
| shpt | 647 | NA | 4.92E-03 (1.43E-01) | 0.53 | 5.29E-03 (7.08E-03) | 1.74E-04 (7.83E-01) | 0.50 | 1.87E-04 (1.28E-03) |
| stm2 | 1870 | 87.57 | 1.41E-06 (9.65E-01) | 0.51 | 1.27E-06 (4.77E-05) | 1.19E-03 (1.99E-01) | 0.51 | 1.07E-03 (1.65E-03) |
| twg2 | 3760 | 70.83 | 3.39E-03 (2.81E-03) | 0.53 | 3.45E-03 (2.29E-03) | 2.72E-03 (7.51E-03) | 0.53 | 2.76E-03 (2.05E-03) |

| **PGC Cohorts** | | | **Downsampled LifetimeMDD** | | | **LifetimeMDD** | | |
|---|---|---|---|---|---|---|---|---|
| | | | **P value threshold = 0.05** | | | **P value threshold = 0.05** | | |
| **Cohort** | **Nsamples** | **%Cases DSM-MDD** | **Nagelkerke's r2 (P)** | **AUC** | **h2_Liab (SE)** | **Nagelkerke's r2 (P)** | **AUC** | **h2_Liab (SE)** |
| boma | 1648 | 92.48 | 1.96E-03 (1.30E-01) | 0.51 | 1.86E-03 (2.38E-03) | 1.66E-03 (1.63E-01) | 0.48 | 1.58E-03 (2.18E-03) |
| col3 | 1952 | 95.85 | 1.71E-02 (7.60E-02) | 0.56 | 1.54E-02 (1.72E-02) | 2.54E-03 (6.58E-02) | 0.52 | 2.70E-03 (2.93E-03) |
| edi2 | 657 | NA | 5.81E-03 (1.01E-01) | 0.54 | 5.28E-03 (6.12E-03) | 7.46E-03 (6.28E-02) | 0.54 | 6.78E-03 (6.86E-03) |
| gens | 2363 | 98.24 | 4.47E-03 (5.13E-03) | 0.53 | 4.07E-03 (2.88E-03) | 9.27E-03 (5.54E-05) | 0.55 | 8.44E-03 (4.13E-03) |
| gep3 | 3318 | 50.43 | 1.34E-03 (1.20E-01) | 0.48 | 1.83E-03 (2.36E-03) | 2.65E-03 (2.90E-02) | 0.52 | 3.60E-03 (3.28E-03) |
| grdg | 941 | NA | 1.68E-03 (2.78E-01) | 0.52 | 1.50E-03 (2.75E-03) | 9.01E-03 (1.18E-02) | 0.55 | 8.10E-03 (6.33E-03) |
| grnd | 1304 | 98.54 | 5.97E-03 (1.76E-02) | 0.54 | 5.63E-03 (4.69E-03) | 4.21E-03 (4.61E-02) | 0.53 | 3.98E-03 (3.97E-03) |
| i2b3 | 1873 | NA | 3.99E-03 (2.01E-02) | 0.53 | 3.63E-03 (2.99E-03) | 4.36E-03 (1.51E-02) | 0.54 | 3.96E-03 (3.18E-03) |
| mmi2 | 1101 | 49.83 | 2.41E-03 (1.60E-01) | 0.52 | 2.17E-03 (3.05E-03) | 1.71E-03 (2.36E-01) | 0.48 | 1.54E-03 (2.55E-03) |

| | | | | | | | | |
|---|---|---|---|---|---|---|---|---|
| mmo4 | 635 | 42.8 | 8.39E-04 (5.71E-01) | 0.52 | 7.68E-04 (2.22E-03) | 2.57E-04 (7.54E-01) | 0.52 | 2.35E-04 (1.93E-03) |
| nes1 | 3096 | 94.11 | 5.77E-03 (2.52E-04) | 0.54 | 5.19E-03 (2.82E-03) | 1.25E-02 (6.97E-08) | 0.55 | 1.13E-02 (4.13E-03) |
| qi3c | 1443 | 95.53 | 7.03E-03 (2.50E-02) | 0.54 | 6.49E-03 (5.74E-03) | 1.74E-02 (4.17E-04) | 0.57 | 1.61E-02 (8.98E-03) |
| qi6c | 1089 | 93.19 | 1.21E-02 (1.67E-03) | 0.56 | 1.09E-02 (6.88E-03) | 1.12E-02 (2.45E-03) | 0.55 | 1.02E-02 (6.64E-03) |
| qio2 | 1091 | 93.98 | 1.38E-04 (7.37E-01) | 0.51 | 1.24E-04 (7.64E-04) | 3.51E-03 (9.07E-02) | 0.53 | 3.15E-03 (3.66E-03) |
| rad3 | 3400 | 88.08 | 4.20E-03 (1.55E-03) | 0.53 | 3.80E-03 (2.37E-03) | 8.75E-03 (4.81E-06) | 0.54 | 7.92E-03 (3.43E-03) |
| rage | 549 | 92.24 | 9.03E-03 (5.93E-02) | 0.55 | 8.28E-03 (8.38E-03) | 5.38E-03 (1.46E-01) | 0.53 | 4.93E-03 (6.75E-03) |
| rau2 | 601 | 97.75 | 1.14E-02 (2.52E-02) | 0.55 | 1.07E-02 (9.32E-03) | 1.06E-02 (3.13E-02) | 0.55 | 9.95E-03 (9.01E-03) |
| rde4 | 649 | 93.94 | 8.15E-03 (6.69E-02) | 0.55 | 9.55E-03 (1.03E-02) | 3.28E-03 (2.46E-01) | 0.52 | 3.84E-03 (6.53E-03) |
| rot4 | 1269 | 50.67 | 4.40E-06 (9.53E-01) | 0.51 | 5.31E-06 (1.79E-04) | 2.44E-03 (1.65E-01) | 0.53 | 2.95E-03 (4.23E-03) |
| shp0 | 1453 | 90.35 | 6.59E-04 (4.21E-01) | 0.51 | 7.08E-04 (1.75E-03) | 1.97E-03 (1.65E-01) | 0.52 | 2.11E-03 (3.02E-03) |
| shpt | 647 | NA | 2.51E-03 (2.97E-01) | 0.53 | 2.69E-03 (5.07E-03) | 5.65E-03 (1.17E-01) | 0.54 | 6.08E-03 (7.52E-03) |
| stm2 | 1870 | 87.57 | 1.33E-03 (1.74E-01) | 0.52 | 1.19E-03 (1.74E-03) | 7.02E-04 (3.24E-01) | 0.51 | 6.30E-04 (1.25E-03) |
| twg2 | 3760 | 70.83 | 2.58E-03 (9.21E-03) | 0.53 | 2.62E-03 (2.01E-03) | 8.77E-03 (1.52E-06) | 0.55 | 8.92E-03 (3.68E-03) |

| **PGC Cohorts** | | | **Downsampled GPNoDep** | | | **GPNoDep** | | |
|---|---|---|---|---|---|---|---|---|
| | | | **P value threshold = 0.05** | | | **P value threshold = 0.05** | | |
| **Cohort** | **Nsamples** | **%Cases DSM-MDD** | **Nagelkerke's r2 (P)** | **AUC** | **h2_Liab (SE)** | **Nagelkerke's r2 (P)** | **AUC** | **h2_Liab (SE)** |
| boma | 1648 | 92.48 | 3.91E-07 (9.83E-01) | 0.51 | 3.71E-07 (8.97E-06) | 6.75E-04 (3.74E-01) | 0.52 | 6.41E-04 (1.41E-03) |
| col3 | 1952 | 95.85 | 3.09E-07 (9.94E-01) | 0.52 | 2.78E-07 (1.31E-04) | 2.64E-03 (6.11E-02) | 0.53 | 2.80E-03 (2.98E-03) |
| edi2 | 657 | NA | 7.65E-04 (5.52E-01) | 0.51 | 6.95E-04 (2.51E-03) | 4.80E-03 (1.36E-01) | 0.54 | 4.36E-03 (6.00E-03) |
| gens | 2363 | 98.24 | 3.21E-06 (9.40E-01) | 0.51 | 2.92E-06 (8.04E-05) | 2.08E-05 (8.49E-01) | 0.50 | 1.89E-05 (1.95E-04) |
| gep3 | 3318 | 50.43 | 5.43E-04 (3.23E-01) | 0.52 | 7.40E-04 (1.51E-03) | 2.46E-05 (8.33E-01) | 0.50 | 3.35E-05 (3.25E-04) |
| grdg | 941 | NA | 6.06E-03 (3.90E-02) | 0.54 | 5.44E-03 (5.21E-03) | 1.13E-02 (4.68E-03) | 0.56 | 1.02E-02 (7.11E-03) |
| grnd | 1304 | 98.54 | 2.11E-03 (1.58E-01) | 0.52 | 1.99E-03 (2.78E-03) | 6.56E-04 (4.31E-01) | 0.51 | 6.20E-04 (1.56E-03) |
| i2b3 | 1873 | NA | 1.52E-03 (1.52E-01) | 0.51 | 1.38E-03 (1.86E-03) | 3.94E-03 (2.10E-02) | 0.53 | 3.58E-03 (2.98E-03) |
| mmi2 | 1101 | 49.83 | 1.85E-05 (9.02E-01) | 0.51 | 1.67E-05 (2.80E-04) | 3.94E-04 (5.69E-01) | 0.50 | 3.55E-04 (1.22E-03) |
| mmo4 | 635 | 42.8 | 3.48E-03 (2.49E-01) | 0.53 | 3.19E-03 (3.81E-03) | 2.91E-04 (7.39E-01) | 0.51 | 2.66E-04 (5.17E-04) |
| nes1 | 3096 | 94.11 | 8.79E-04 (1.54E-01) | 0.51 | 7.90E-04 (1.10E-03) | 1.15E-03 (1.03E-01) | 0.52 | 1.03E-03 (1.26E-03) |
| qi3c | 1443 | 95.53 | 1.26E-04 (7.64E-01) | 0.50 | 1.16E-04 (7.70E-04) | 1.32E-03 (3.32E-01) | 0.52 | 1.22E-03 (2.49E-03) |

| | | | | | | | | |
|---|---|---|---|---|---|---|---|---|
| qi6c | 1089 | 93.19 | 2.31E-03 (1.70E-01) | 0.52 | 2.09E-03 (3.03E-03) | 1.14E-03 (3.35E-01) | 0.51 | 1.03E-03 (2.13E-03) |
| qio2 | 1091 | 93.98 | 2.55E-03 (1.49E-01) | 0.53 | 2.29E-03 (3.12E-03) | 1.23E-03 (3.16E-01) | 0.52 | 1.11E-03 (2.19E-03) |
| rad3 | 3400 | 88.08 | 3.39E-03 (4.49E-03) | 0.53 | 3.06E-03 (2.13E-03) | 4.68E-03 (8.30E-04) | 0.53 | 4.23E-03 (2.50E-03) |
| rage | 549 | 92.24 | 2.27E-03 (3.45E-01) | 0.52 | 2.08E-03 (3.81E-03) | 1.73E-03 (4.10E-01) | 0.52 | 1.58E-03 (3.40E-03) |
| rau2 | 601 | 97.75 | 4.48E-03 (1.61E-01) | 0.54 | 4.21E-03 (5.98E-03) | 4.50E-03 (1.60E-01) | 0.54 | 4.23E-03 (5.99E-03) |
| rde4 | 649 | 93.94 | 1.42E-03 (4.45E-01) | 0.52 | 1.66E-03 (4.47E-03) | 2.04E-04 (7.72E-01) | 0.50 | 2.38E-04 (1.54E-03) |
| rot4 | 1269 | 50.67 | 1.83E-03 (2.29E-01) | 0.52 | 2.22E-03 (3.67E-03) | 2.46E-03 (1.64E-01) | 0.53 | 2.97E-03 (4.25E-03) |
| shp0 | 1453 | 90.35 | 7.55E-06 (9.31E-01) | 0.50 | 8.11E-06 (1.76E-04) | 4.98E-05 (8.25E-01) | 0.50 | 5.35E-05 (4.76E-04) |
| shpt | 647 | NA | 9.24E-04 (5.26E-01) | 0.51 | 9.92E-04 (3.15E-03) | 5.93E-03 (1.08E-01) | 0.53 | 6.37E-03 (7.83E-03) |
| stm2 | 1870 | 87.57 | 2.77E-04 (5.35E-01) | 0.50 | 2.48E-04 (7.96E-04) | 4.21E-05 (8.09E-01) | 0.51 | 3.77E-05 (3.06E-04) |
| twg2 | 3760 | 70.83 | 2.65E-04 (4.04E-01) | 0.49 | 2.68E-04 (6.41E-04) | 3.59E-05 (7.59E-01) | 0.50 | 3.64E-05 (2.34E-04) |

**Supplemental Table S22: Out-of-sample prediction of MDD in PGC cohorts**

This table shows the prediction of MDD status indicated in 20 PGC cohorts with polygenic risk scores (PRS) computed from GWAS summary statistics of definitions of depression in UKBiobank. The PRS are calculated using effect sizes at independent (LD r2 < 0.1) SNPs passing P-value thresholds 0.05 only (for results at other P-value thresholds, see Figure 7 and Supplemental Figure S13,S15). We show results from PRS computed from GWAS performed on down-sampled cases and controls ("Downsampled"), as well as on all cases and controls. For each cohort in PGC we show the cohort name (Cohort), number of samples (N) and percentage of cases that fulfill DSM-V criterion A for symptoms (% Cases DSM-MDD). For prediction results from each definition of depression in UKBiobank, we show the Nagelkerke's $r^2$ with the P-value of correlation (P), the area under the curve for prediction accuracy (AUC) and variance of MDD status explained by the PRS under the liability scale, assuming population prevalence of 0.15 (h2PRS_Liab) with its standard error (SE). We draw attention to results in cohort rad3, which has much higher Nagelkerke's $r^2$ and AUC than other cohorts at each P value threshold, likely due to it containing relatives of individuals in the UKBiobank, being a UK-based cohort. We removed this cohort from the calculation of prediction accuracy across all cohorts.

| Category | Definition | PRS P value threshold | Correlation between % DSM-MDD Cases in PGC Cohorts and PRS | | |
|---|---|---|---|---|---|
| | | | Pearson r | Pearson r2 | P value |
| Help-seeking | GPpsy | 0.05 | 0.290 | 0.084 | 0.229 |
| | | 0.1 | 0.138 | 0.019 | 0.574 |
| | | 0.2 | 0.056 | 0.003 | 0.820 |
| | | 0.5 | -0.044 | 0.002 | 0.857 |
| | | 1 | -0.069 | 0.005 | 0.778 |
| Symptom | DepAll | 0.05 | 0.290 | 0.084 | 0.229 |
| | | 0.1 | 0.138 | 0.019 | 0.574 |
| | | 0.2 | 0.056 | 0.003 | 0.820 |
| | | 0.5 | -0.044 | 0.002 | 0.857 |
| | | 1 | -0.069 | 0.005 | 0.778 |
| Self-report | SelfRepDep | 0.05 | -0.231 | 0.053 | 0.342 |
| | | 0.1 | -0.152 | 0.023 | 0.534 |
| | | 0.2 | -0.172 | 0.029 | 0.482 |
| | | 0.5 | -0.130 | 0.017 | 0.596 |
| | | 1 | -0.138 | 0.019 | 0.572 |
| EMR | ICD10Dep | 0.05 | -0.016 | 0.000 | 0.949 |
| | | 0.1 | -0.321 | 0.103 | 0.180 |
| | | 0.2 | -0.356 | 0.127 | 0.134 |
| | | 0.5 | -0.210 | 0.044 | 0.388 |
| | | 1 | -0.235 | 0.055 | 0.332 |
| DSM-based | LifetimeMDD | 0.05 | 0.448 | 0.200 | 0.055 |
| | | 0.1 | 0.512 | 0.262 | 0.025 |
| | | 0.2 | 0.438 | 0.192 | 0.061 |
| | | 0.5 | 0.425 | 0.181 | 0.070 |
| | | 1 | 0.419 | 0.175 | 0.074 |
| No-MDD | GPNoDep | 0.05 | -0.016 | 0.000 | 0.949 |
| | | 0.1 | -0.321 | 0.103 | 0.180 |
| | | 0.2 | -0.356 | 0.127 | 0.134 |
| | | 0.5 | -0.210 | 0.044 | 0.388 |
| | | 1 | -0.235 | 0.055 | 0.332 |

**Supplemental Table S23: Correlation between prediction Naglekerke's r2 and percentage DSM-MDD cases in PGC cohorts**

This table shows the Pearson correlation coefficient (Pearson r) and $r^2$ (Pearson r2) with P value between PRS prediction Naglekerke's $r^2$ and percentage of DSM-MDD cases in each PGC cohort, for PRS derived from each P value threshold from GWAS of each definition of depression in UKBiobank. This table contains results from PRS build from GWAS on the full dataset.
